# A comprehensive identification protocol for postcranial bones of Southeast Asian Ruminantia. Part 3: Morphological criteria for large-sized species

**DOI:** 10.64898/2026.10.23.761339

**Authors:** Sirikanya Chantasri, Supalak Mithong, Julien Claude, Papavee Seesod, Corentin Bochaton

**Author notes:** Corresponding authors: Correspondence: SM, CB.

## Abstract

Postcranial remains of large-bodied ruminants are abundant in the Late Quaternary record of continental Southeast Asia, yet their taxonomic attribution remains one of the least resolved problems in the zooarchaeology of this biodiversity-rich region. We address it here with an osteological identification protocol covering the six large ruminant species of the mainland: *Bubalus bubalis*, *Bos javanicus*, *Bos sauveli*, *Bos gaurus*, *Budorcas taxicolor* and *Rusa unicolor*, built from a comparative sample of 109 skeletons. A total of 148 discrete characters over 27 skeletal elements representing 442 states were defined on a restricted set of individuals, then scored across the entire sample. Most proved remarkably stable within taxa with 80.0% of the character/taxon combinations showing a dominant state at a frequency above 95%, and a substantial number of states occurred in a single family, subfamily, genus or species. Character combinations for taxonomic identification were then evaluated by the use of two supervised machine learning methods, a random forest model and a naïve Bayes classifier at each of four taxonomic ranks. The two behaved almost identically, with mean accuracies of 95–94% for families, 93% for subfamilies, 89% for genera and 76% for species; only the three closely related and unevenly sampled species of *Bos* resisted consistent separation. The weighted Naïve Bayes models are distributed as a standalone web application in which any combination of states observed on a single element returns attribution probabilities for every rank. Together these results offer a transparent and statistically explicit alternative to expert judgement for the identification of Late Quaternary postcranial mammal remains, in Southeast Asia and elsewhere.

## Introduction

Animal bones represent some of the most commonly recovered archaeological evidence and serve as a major source of information in zooarchaeology, a discipline that integrates archaeology and paleontology to investigate past interactions between human populations and animals (Reitz & Wing, 2008). Most zooarchaeological research relies on the taxonomic identification of frequently fragmented bones and dental remains, a process that typically depends on morphological analysis and comparison with modern reference specimens. This approach can be difficult to reproduce, particularly when multiple closely related or similarly sized taxa are present in the same bone assemblages, and when access to appropriate comparative collections is limited. In biodiverse tropical regions, these challenges are particularly pronounced because archaeological assemblages may contain a wide range of taxa with overlapping morphological characteristics (LeFebvre & Sharpe, 2018) and the appropriate comparative osteological collections, when they exist, are often stored in Western museums far from the area concerned by the study. In addition to these specific issues, the overall identification process remains highly dependent on the quality of reference collections and the experience of the analyst (Reitz & Wing, 2008; Wolverton, 2013), which complicates the comparison and combination of results when identification criteria are not explicitly stated. Combined, these issues make the conduct of good zooarchaeological studies in tropical areas very challenging and most often impossible for local scholars.

These problems call for comprehensive comparative collections, sustained training, and replicable protocols (Driver, 2011; Lyman, 2019; Wolverton, 2013). Biomolecular methods such as ZooMS and ancient DNA can overcome some limitations associated with fragmented or morphologically non-diagnostic remains; however, these approaches remain costly and are constrained by poor collagen or DNA preservation (Buckley, 2018; Buckley et al., 2010; Rizzi et al., 2012). A complementary strategy to tackle this problem more realistically is to develop qualitative diagnostic osteological criteria tested on multiple specimens, an approach already applied to several taxa (Bochaton et al., 2016; Bochaton, Ivanov, et al., 2019; Hanot & Bochaton, 2018; Koolstra et al., 2019; Wang et al., 2020; Zeder & Lapham, 2010; Zhang et al., 2022). Yet such criteria remain largely undeveloped for mammal faunas of high-biodiversity tropical regions.

In continental Southeast Asia, despite its high modern biodiversity and rich archaeological record, both biomolecular (Siripan et al., 2019; Wannajuk et al., 2013) and morphological approaches to taxonomic identification remain relatively underdeveloped (Rodrigo & Manamendra-Arachchi, 2022; Wang et al., 2020). Taxonomic and osteological research on reptiles has recently progressed there (Bochaton et al., 2023, 2025; Bochaton, Hanot, et al., 2019; Claude et al., 2019) but substantial gaps remain for most vertebrate taxa whose postcranial anatomy remains poorly known or insufficiently described. This situation is particularly damaging for zooarchaeological studies that mostly rely on fragmented postcranial bone elements. It could, however, be partly solved with a focus on the taxa that represent the bulk of such assemblages, as human groups tend to exploit a limited number of species. In the case of continental Southeast Asia, several studies have highlighted a predominance of ruminants in prehistoric zooarchaeological assemblages (Van Den Bergh et al., 2001; Auetrakulvit, 2004; Métais et al., 2007; Mudar & Anderson, 2007; Stenhouse, 2019). Ruminants not only played a crucial role in the subsistence strategies of past human populations but also served as valuable indicators of past environmental conditions. Yet, most of the studies incorporating them lack specific osteological criteria to distinguish species within the local representatives of the families Cervidae and Bovidae. The limited availability of reliable postcranial criteria has led researchers to rely heavily on dental remains. Consequently, when teeth are absent, species-level identification becomes more difficult and taxonomic interpretations become less reliable (Amano et al., 2016; Suraprasit et al., 2016a, 2019, 2021, 2025). Defining a standardized protocol for identifying Ruminantia postcranial remains would thus be very helpful to improve the quality and accuracy of species identification in Late Quaternary contexts.

To address this gap, we developed a postcranial identification protocol for the Ruminantia of continental Southeast Asia, built from three complementary studies: osteometry (Seesod et al., 2026), the morphology of small and medium-sized species (Mithong et al., 2026), and the morphology of large-sized species, presented here. In this third and final paper of the series, we focus on six species: water buffalo (*Bubalus bubalis*), banteng (*Bos javanicus*), kouprey (*Bos sauveli*), gaur (*Bos gaurus*), takin (*Budorcas taxicolor*), and sambar (*Rusa unicolor*), whose overlapping body sizes and similar postcranial proportions make them difficult to separate in archaeological assemblages.

Some of these species have been treated before, but only partially: on smaller reference samples, without statistical validation of the proposed criteria, or without including all of the taxa occurring in continental Southeast Asia (Haraniya et al., 2016; Zhang et al., 2022). We describe here readily identifiable, non-metric osteomorphological criteria for their postcranial identification, test them statistically across the full comparative sample, and deploy the resulting models as a standalone web application, so that identifications of the Late Quaternary record of these taxa can be reproduced and their reliability independently assessed.

## Material and Methods

### Investigated species and specimens

The Ruminantia species included in this study were selected based on their documented occurrence in continental Southeast Asia (Lekagul & McNeely, 1988; Francis, 2019) or their occurrence in recent paleontological studies (Suraprasit et al., 2016b). All the large-sized genera currently represented in the region were sampled, with the addition of *Bos sauveli,* a possibly very recently extinct species (Hassanin et al., 2005). The present study focuses on “large-sized” ruminants and excludes all but one of the medium-sized taxa investigated separately in Part 2 of this paper series (Mithong et al., 2026). The latter include the genera: *Axis, Capricornis, Cervus, Elaphodus, Moschus, Muntiacus, Naemorhedus, Pseudois*, and *Rucervus.* The genus *Rusa* has been integrated in both Part 2 and the present study because of its intermediate size between the groups of large and medium-sized species (Seesod et al., 2026). The list of “large-sized” species investigated in this work has been defined based on their skeletal measurements recorded in Part 1 of this paper series (Seesod et al., 2026). They correspond to the genera *Bubalus*, *Bos*, *Budorcas*, and *Rusa*. We hereby investigate osteological characters allowing for the distinction of these different species. A review of the systematics and taxonomy of the considered species, as well as details of the composition of our data set for each of these species, was undertaken in the first paper of this series (Seesod et al., 2026). Most notably, we repeat here that our samples of *Bos javanicus* and *Bubalus bubalis* may include domestic forms in a proportion that we cannot estimate, while our specimens of *Bos gaurus* only correspond to wild individuals.

The sample investigated here comprises 109 specimens representing six species: *Bubalus bubalis (N=24); Bos javanicus (N=17); Bos sauveli (N=3); Bos gaurus (N=18); Budorcas taxicolor (N=18);* and *Rusa unicolor (N=29).* These specimens were obtained from five museum collections in France, Belgium, Germany, and the United States (see full details in Supplementary Material 01; Chantasri et al., 2026).

### Osteological character definition and recording protocol

In the initial step of the study, one or two specimens per species were compared to define meaningful osteological differences among the various taxa. The specimens used in this first step are the following: *Bubalus bubalis* (MNHN-ZM-AC-1857-19); *Bos javanicus (MNHN-ZM-AC-1944- 101; MNHN-ZM-AC-1967-1699)*; *Bos sauveli* (MNHN-ZM-AC-1871-350); *Bos gaurus (MNHN-ZM-AC-2013-27)*; *Budorcas taxicolor (MNHN-ZM-AC-2017-1199; MNHN-ZM-AC-1925-240)*; and *Rusa unicolor* (MNHN-ZM-AC-2013-28). The osteological characters were described following the anatomical nomenclature of Barone (1976). Character observations were initially illustrated through direct visual observation of specimens in the collections and examination of 3D models. These 3D models were produced using an EinScan SP V2 scanner and an Artec Eva 3D scanner, with photogrammetric models generated using Agisoft PhotoScan Professional Edition. These models have been used to illustrate the characters in the present paper with the addition of the following specimens: *Bos gaurus (MNHN-ZM-AC-2017-1204 and ZSM 1982/105)*, *Rusa unicolor* (MNHN-ZM-AC-1878-13). All of the 3D models illustrated are provided as Supplementary Material (Supplementary Material 08; Chantasri et al., 2026).

This initial set of defined osteological characters was then recorded across the entire sample to assess their reliability and relevance. The resulting dataset (Supplementary Material 02, Supplementary Material 03; Chantasri et al., 2026) was then analyzed according to the protocol described below.

### Analytical Protocol of the character dataset

The analytical protocol is the same as the one used in Part 2 of this paper series (Mithong et al., 2026). The initial dataset underwent a preliminary cleaning step before analysis. During this step, all character states that were both very rare (occurring in fewer than 10% of the specimens of every species in which they were observed) and ambiguous character states (which represented 2.7% of the recorded character states) were removed. These represent character states defined in the initial observation but never scored in the dataset. In that case, the character was turned to NA for the concerned specimens. This cleaning step was performed manually on the original scoring table before the analyses described below; the resulting dataset (Supplementary Material 02, Chantasri et al., 2026) thus already reflects these exclusions. The remaining characters and character states were then evaluated by identifying those unique to a given taxon, as well as by using the maximum frequency of each character state within a taxon. This approach provided a general indication of their stability and relevance for taxonomic identification. The characters were already performing very well with this initial evaluation, but in order to make the protocol match that of the previous study (Mithong et al., 2026), we had to conduct more elaborate analyses so the results of the two studies could be comparable and used together.

To estimate the discriminatory power of each skeletal element and character combination, a Random Forest (RF) model was applied to binary-encoded (dummy-variable) character states. Because subfossil skeletal elements are most commonly recovered in isolation, each anatomical element (a complete bone) was analyzed independently. For each element, 1000 independent cross-validation iterations were performed for every possible character subset of size k, ranging from 1 to the maximum number of characters available for that element, as most present a different number of characters. Each iteration was based on a stratified random split retaining 20% of specimens as a test set, with the constraint that at least two taxa must be represented in the training set. Classification accuracy was evaluated across all four taxonomic ranks (Family, Subfamily, Genus, and Species).

Because the deployed identification application must accept any user-defined combination of characters as input, a weighted multinomial Naive Bayes (NB) classifier was also implemented as the inference engine for each skeletal element and taxonomic rank, since it can natively incorporate any subset of the available evidence at prediction time. Its performance was evaluated under the same parameters as the Random Forest model. For this latter approach, likelihood estimates were derived from the observed frequencies of each character state within each taxon and the discriminatory power of each character (estimated by their information gain). In both the RF and NB approaches, for each identification query, analyses are conducted independently for each taxonomic rank (Family, Subfamily, Genus, and Species). While this approach may occasionally produce discrepancies between probability estimates at different ranks, it ensures that the attribution probabilities at each rank are as unbiased and internally consistent as possible given the available reference data. This approach has been preserved in the web application integrating the NB approach (see below).

All the analyses were performed in the software R v. 4.6.0 (R Core Team, 2026) (https://cran.r-project.org), with the packages dplyr (Wickham, François, et al., 2026), stringr (Wickham, 2025), ranger (Wright et al., 2026), readr (Wickham, Hester, et al., 2026), and data.table (Barrett et al., 2026). The NB approach was conducted with a homemade function. Coding assistance was provided by the generative AI model Claude (Anthropic), but all code was reviewed and validated by the authors prior to use. All the code necessary to reproduce these analyses is provided as supplementary information (Supplementary Material 04, Chantasri et al., 2026).

The weighted multinomial Naïve Bayes models were deployed as a web application to perform bone identification based on character-state input from a user (Bochaton et al., 2026). Every reference specimen was submitted to the deployed models for each anatomical element it is scored for. For each element, we tested every number of characters that can be entered at once, from a single character up to the maximum available on that bone, and for each of these we drew up to 100 distinct combinations of the character states recorded, so as to reproduce the range of situations a user may face (Supplementary Material 04, Chantasri et al., 2026). The probabilities returned were then compared with whether the attribution was correct. Reliability rises steeply with the probability returned, and exceeds 90% from 80% upwards at the family, subfamily and genus ranks, and from 90% upwards at the species rank. The application applies these two thresholds and highlights the lowest rank that reaches its own; where none does, no attribution is highlighted. Instructions on how to use this application and how to combine the results with those of the previous osteometric study (Seesod et al., 2026) are provided in the Discussion section and were also already discussed in the previous paper of this series (Mithong et al., 2026).

## Results

### I) Character descriptions

In the following section, we describe and illustrate all the osteological features observed and included in the subsequent analyses. Unless otherwise stated, the illustrated specimens are as follows: *Bubalus bubalis* (MNHN-ZM-AC-1857-19); *Bos javanicus* (MNHN-ZM-AC-1944-101; MNHN-ZM-AC-1967-1699); *Bos sauveli* (MNHN-ZM-AC-1871-350); *Bos gaurus* (MNHN-ZM-AC- 2013-27); *Budorcas taxicolor* (MNHN-ZM-AC-2017-1199; MNHN-ZM-AC-1925-240); and *Rusa unicolor* (MNHN-ZM-AC-2013-28). The side and anatomical position of each bone were recorded using standardized abbreviations as follows: ant. (anterior), post. (posterior), dor. (dorsal), ven. (ventral), prox. (proximal), dis. (distal), med. (medial), lat. (lateral), pal. (palmar), and plan. (plantar).

#### Characters of the scapula

##### 1. Morphology of the scapular spine’s tuberosity in lateral view. (Fig. 1)

**Figure 1.**
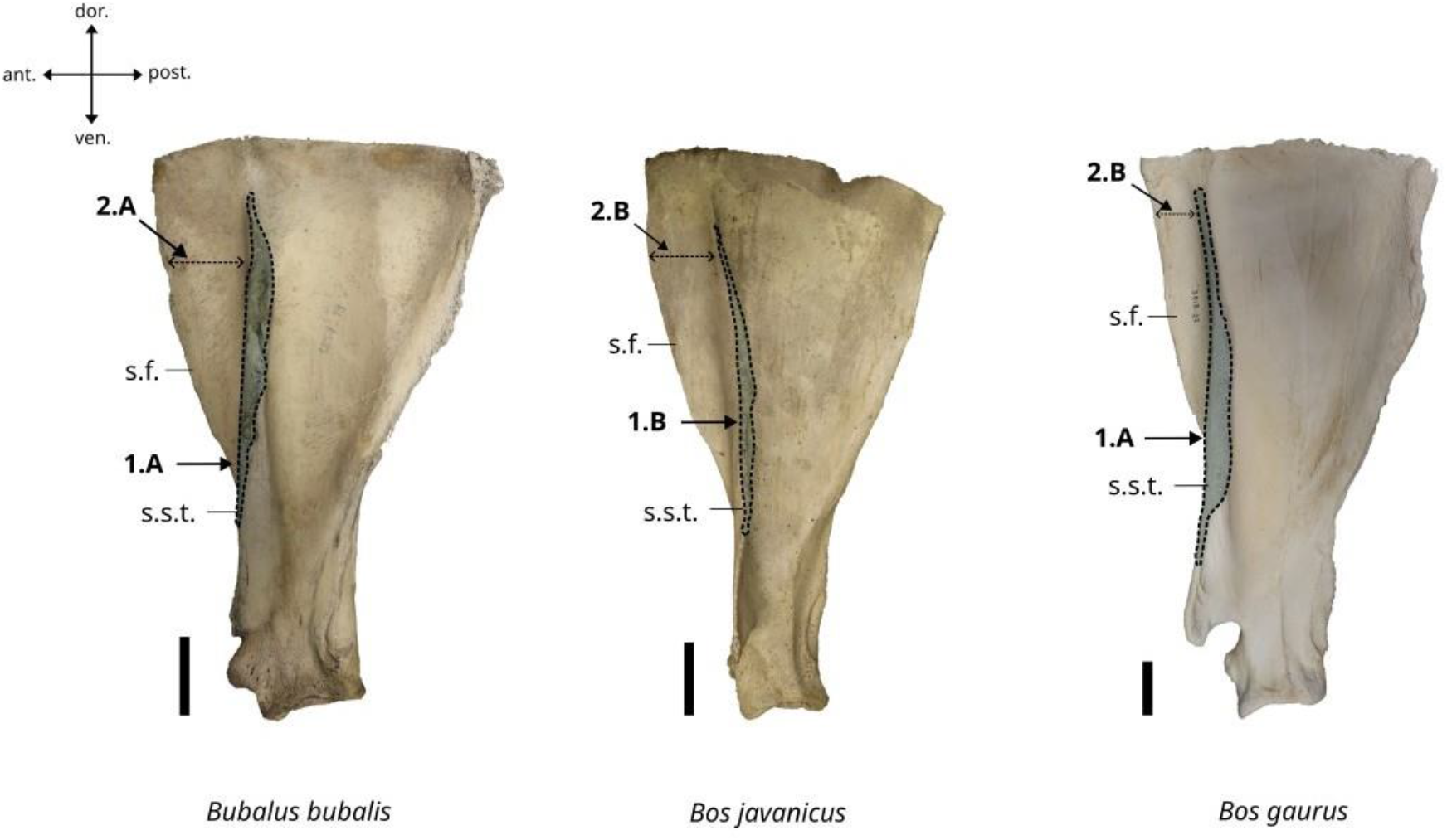
Character states visible on the lateral face of the left scapula (characters 1 and 2). Abbreviations: s.f.: supraspinous fossa, s.s.t.: scapular spine’s tuberosity. Scalebar = 5 cm.

A. The tuberosity is well-developed, and the lateral margin of the spine is thick.
B. The tuberosity is poorly developed, and the lateral margin of the spine is thin.

##### 2. Anterior expansion of the supraspinous fossa beyond the anterior limit of the scapular spine. (Fig. 1)

A. Strongly expanded.
B. Weakly expanded.

##### 3. Morphology of the ventral part of the scapular spine. (Fig. 2)

**Figure 2.**
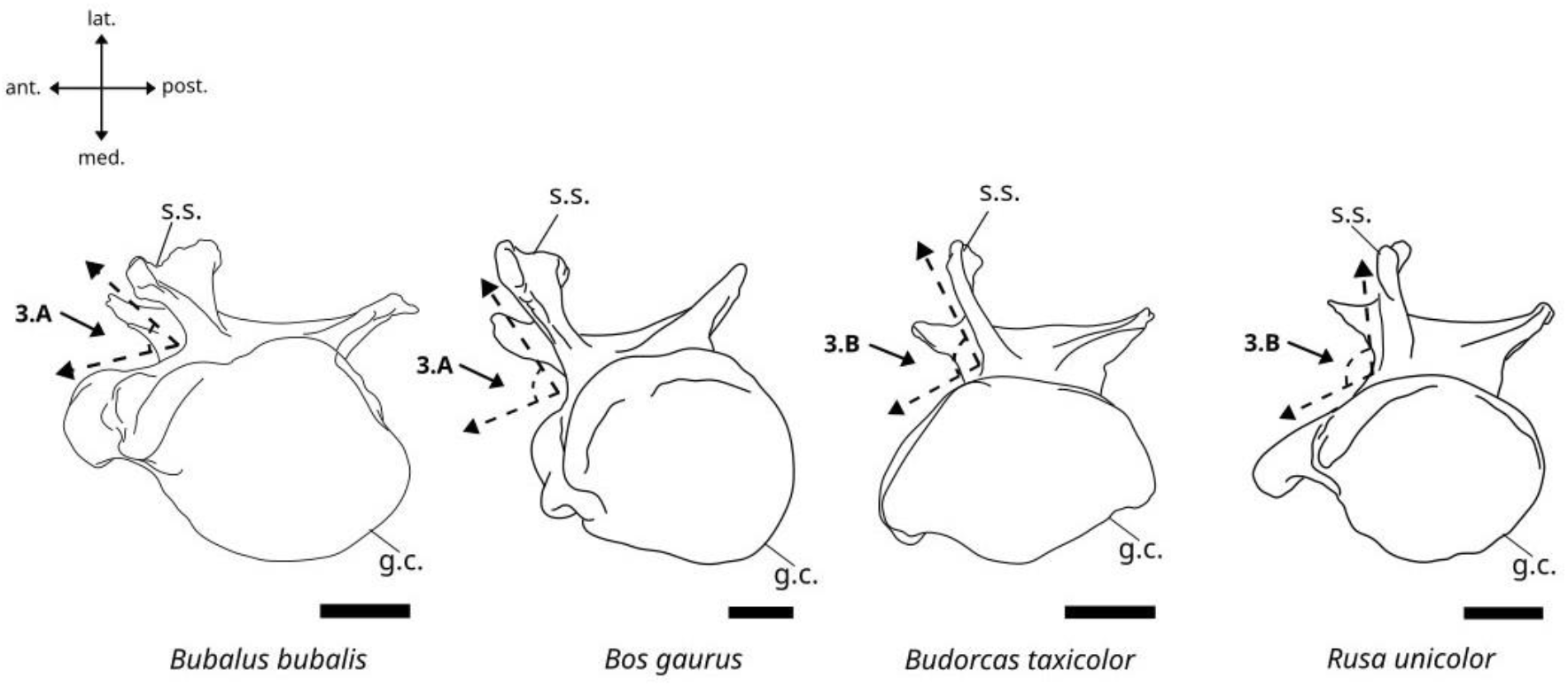
Character states visible on the ventral face of the left scapula (character 3). Abbreviations: s.s.: scapular spine, g.c.: glenoid cavity. Scalebar = 2 cm.

A. In ventral view, the ventral margin of the scapular spine is inclined and forms an angle of 60° with the core of the scapula.
B. In ventral view, the ventral margin of the scapular spine is straight and forms a 90° angle with the core of the scapula.

##### 4. Distance between the scapular spine’s dorsal limit and the scapula’s dorsal limit in lateral view. (Fig. 3)

**Figure 3.**
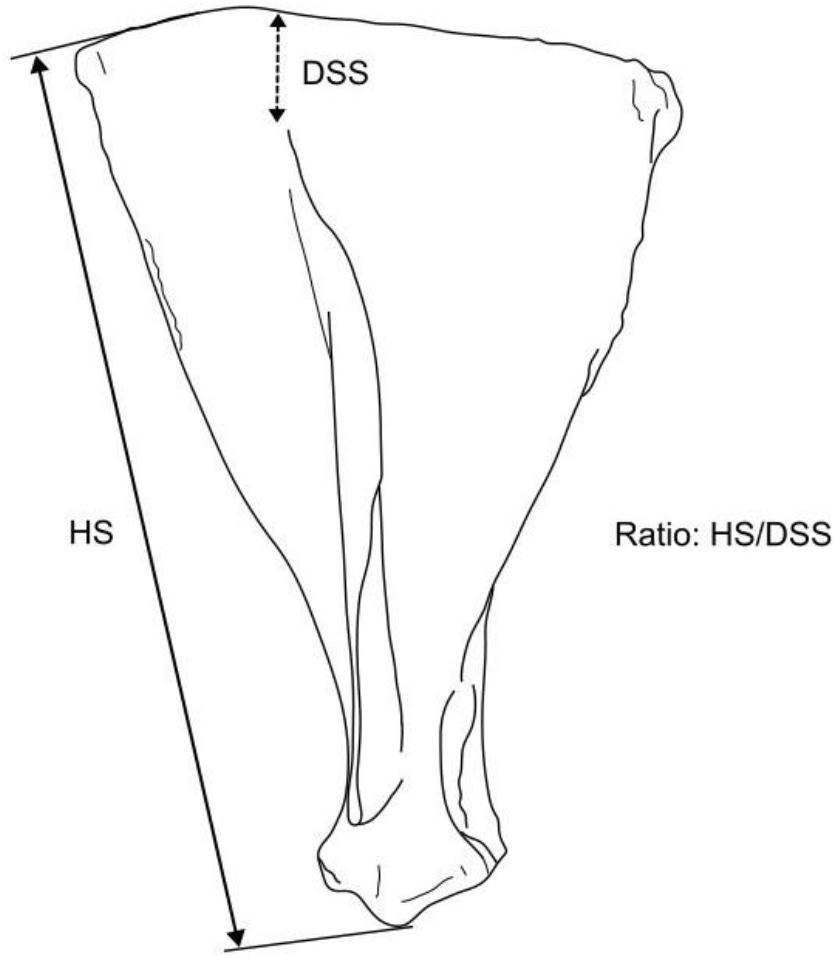
Measurements needed to code character 4.

A. The scapular spine is reaching the dorsal limit of the bone.
B. The scapular spine is nearly reaching the dorsal limit of the bone (Ratio HS/DSS>7).
C. The scapular spine is far from reaching the dorsal limit of the bone (Ratio HS/DSS<7).

##### 5. In ventral view, occurrence of an indentation on the anteromedial margin of the glenoid cavity next to the coracoid process. (Fig. 4)

**Figure 4.**
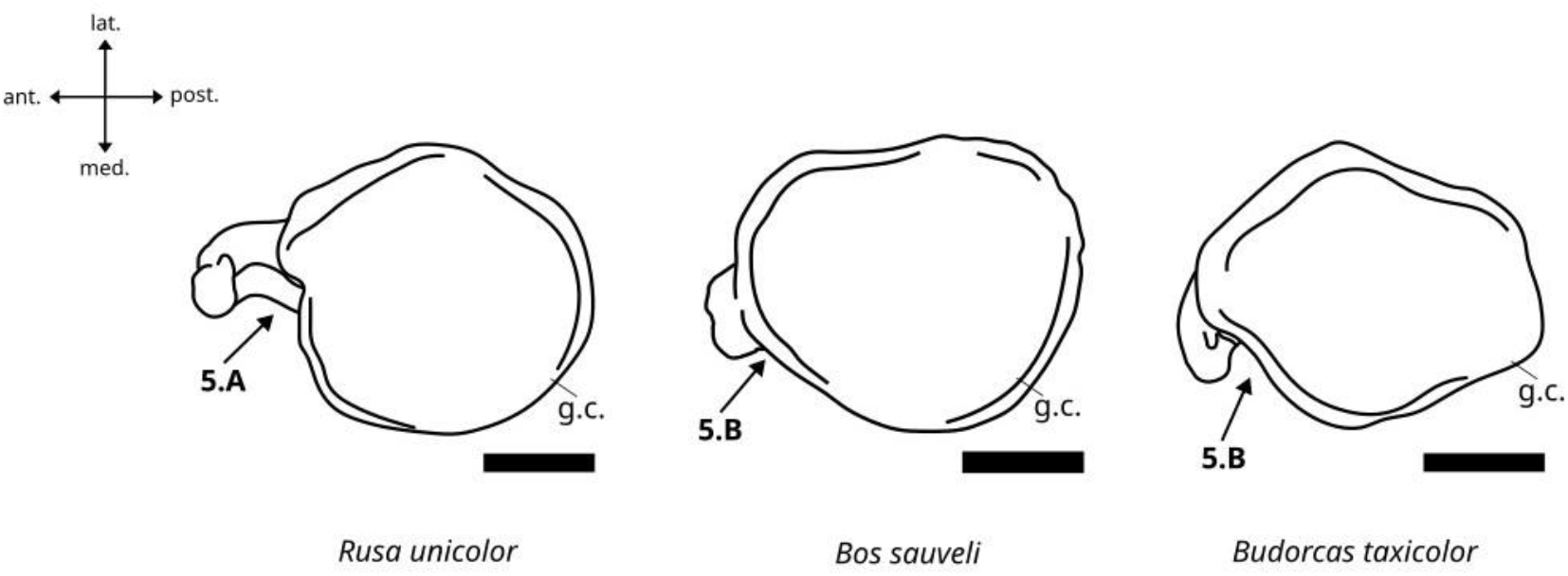
Character states visible on the ventral face of the left scapula (character 5). Abbreviation: g.c.: glenoid cavity. Scalebar = 2 cm.

A. Present and well-marked.
B. Shallow or absent.

##### 6. Morphology of the supraglenoid tubercle in medial view. (Fig. 5)

**Figure 5.**
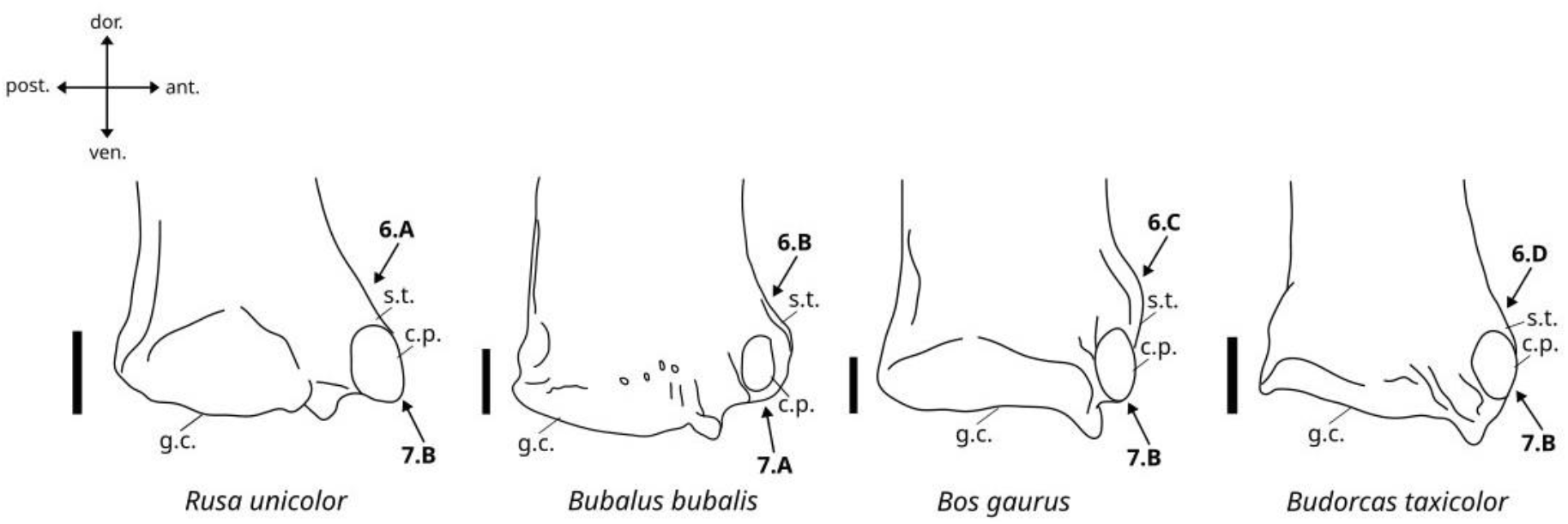
Character states visible on the medial face of the left scapula (characters 6 and 7). Abbreviations: s.t.: supraglenoid tubercle, c.p.: coracoid process, g.c.: glenoid cavity. Scalebar = 2 cm.

A. The supraglenoid tubercle is very well-developed and expanded in anterior direction.
B. The supraglenoid tubercle is moderately developed and expanded in anterior direction.
C. The supraglenoid tubercle is well-developed and expanded in anterior direction.
D. The supraglenoid tubercle is poorly developed.

##### 7. Morphology of the coracoid process in medial view. (Fig. 5)

A. The coracoid process is poorly individualized.
B. The coracoid process is well-individualized.

##### 8. Shape of the glenoid cavity in ventral view. (Fig. 6)

**Figure 6.**
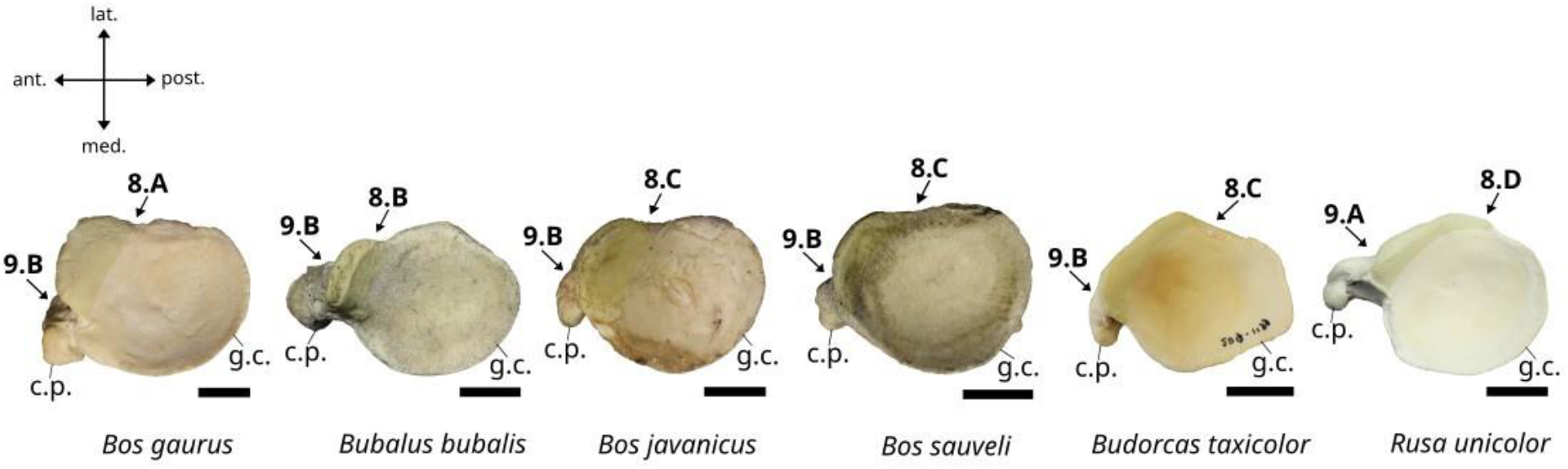
Character states visible on the ventral face of the left scapula (characters 8 and 9). Abbreviations: c.p.: coracoid process, g.c.: glenoid cavity. Scalebar = 2 cm.

A. Well-expanded in antero-lateral and ventral directions and not over the glenoid process area.
B. Well-expanded in antero-ventral direction and over the glenoid process area.
C. Not expanded in antero-ventral direction but expanded over the glenoid process area.
D. Not expanded in antero-ventral direction and not expanded over the glenoid process area.

##### 9. Projection of the coracoid process in ventral view. (Fig. 6)

A. Strongly projected in antero-medial direction.
B. Weakly projected in antero-medial direction.

#### Characters of the humerus

##### 10. Morphology of the medio-dorsal margin of the lesser trochanter. (Fig. 7)

**Figure 7.**
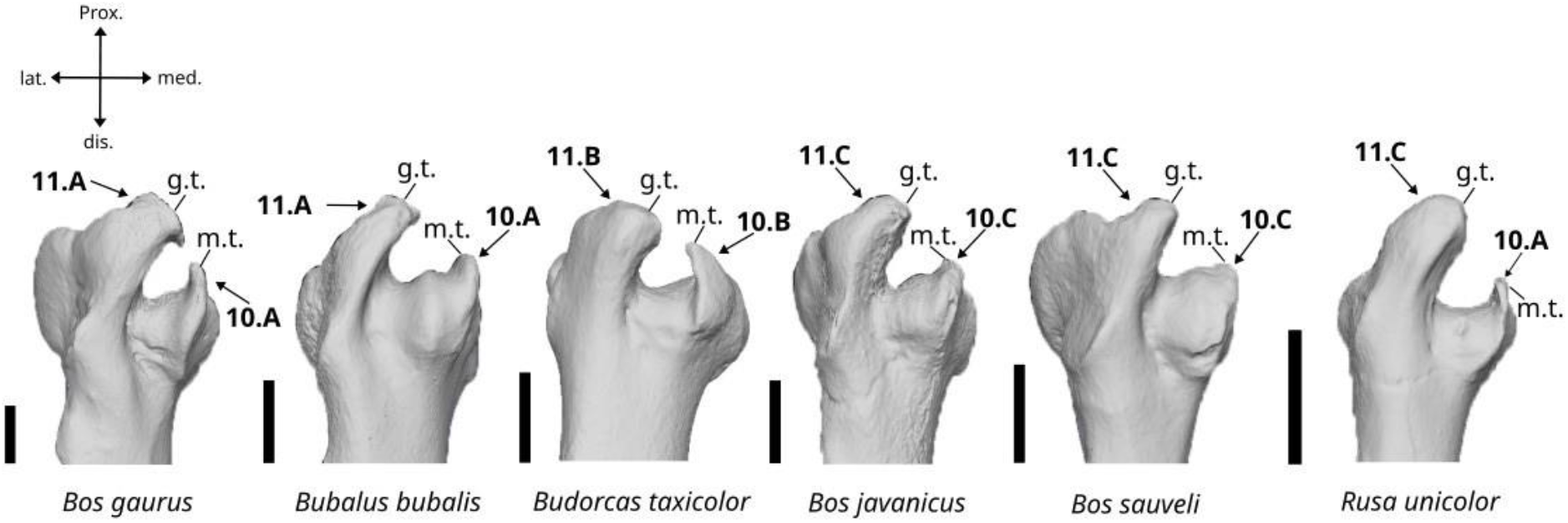
Character states visible on the anterior face of the right humerus (characters 10 and 11). Abbreviations: g.t.: greater trochanter, m.t.: minor trochanter. Scalebar = 5 cm.

A. The medio-dorsal margin of the minor trochanter bears a thin and straight crest.
B. The medio-dorsal margin of the minor trochanter bears a wide crest inclined in lateral direction.
C. The medio-dorsal margin of the minor trochanter bears no crest.

##### 11. Inclination of the greater trochanter. (Fig. 7)

A. The greater trochanter is strongly inclined in medial direction, at an angle greater than 25°.
B. The greater trochanter is moderately inclined in medial direction, at an angle ranging from 10° to 25°.
C. The greater trochanter is weakly inclined in medial direction, at an angle of less than 10°.

##### 12. Expansion and convexity of the greater trochanter. (Fig. 8)

**Figure 8.**
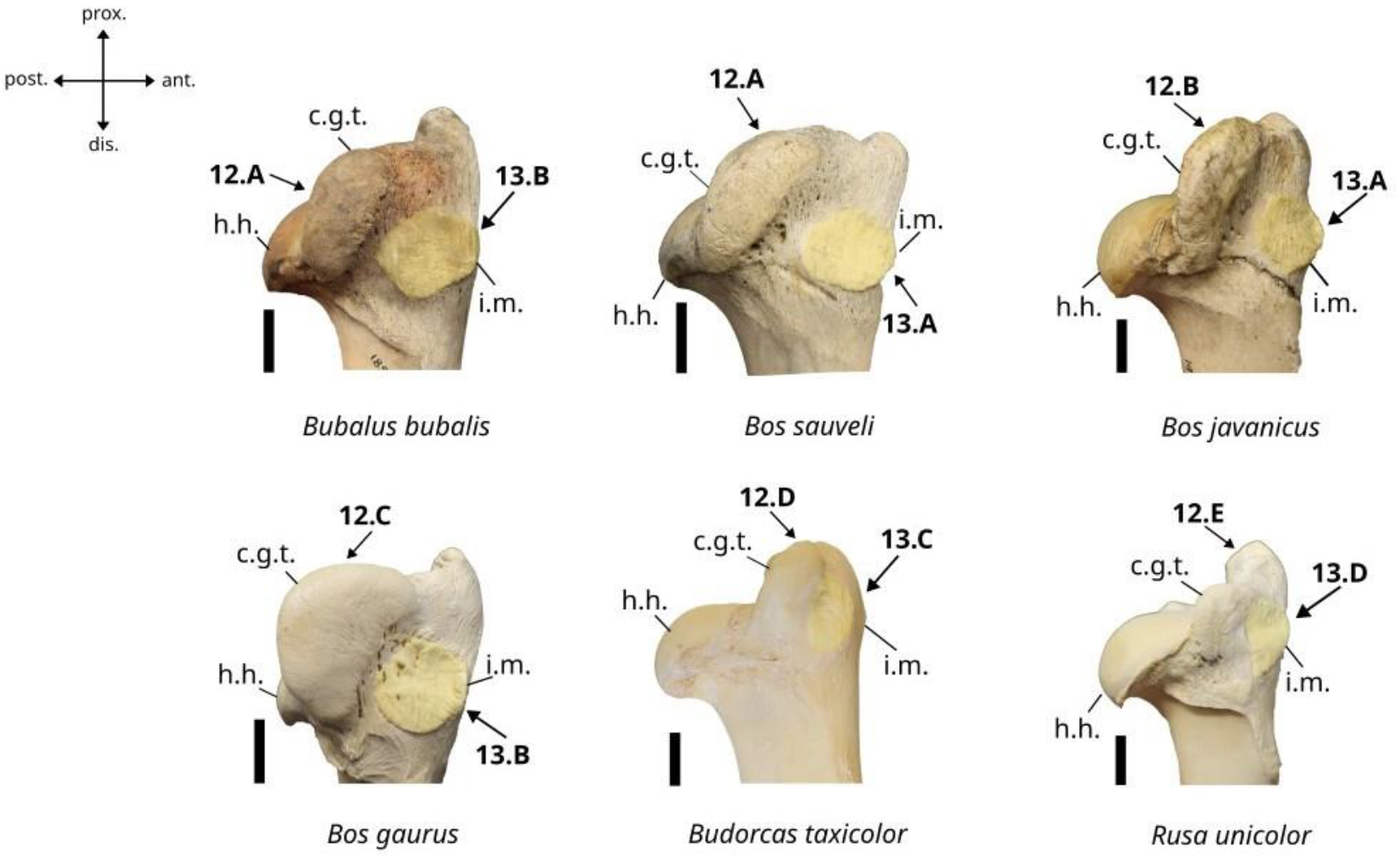
Character states visible on the lateral face of the right humerus (characters 12 and 13). Abbreviations: c.g.t.: convexity of the greater trochanter, h.h.: humeral head, i.m.: *infraspinatus* muscle attachment area. Scalebar = 3 cm.

A. The greater trochanter is wide and short in lateral view and well-expanded in postero-medial direction in dorsal view. Its convexity is well-developed and well-expanded in posterior direction in lateral view.
B. The greater trochanter is elongated in lateral view and moderately expanded in postero-medial direction in dorsal view. Its convexity is moderately developed and moderately expanded in posterior direction in lateral view.
C. The greater trochanter is wide and short in lateral view and moderately expanded in postero- medial direction in dorsal view. Its convexity is very well-developed and strongly expanded in posterior direction in lateral view, to the point that it reaches the posterior limit of the humeral head.
D. The greater trochanter is thin and high in lateral view, and not in posterior direction in dorsal view. Its convexity is small and weakly expanded in posterior direction.
E. The greater trochanter is thin and high in lateral view and expanded in posterior direction in dorsal view. Its convexity is well-developed and moderately expanded in posterior direction in lateral view.

##### 13. Morphology and position of the attachment area of the infraspinatus muscle. (Fig. 8)

A. The attachment area of the *infraspinatus* muscle is well expanded beyond the anterior limit of the bone in lateral view and is located in a low position on the greater tubercle.
B. The attachment area of the *infraspinatus* muscle is weakly expanded beyond the anterior limit of the bone in lateral view and is located in a low position on the greater tubercle.
C. The attachment area of the *infraspinatus* muscle is not expanded beyond the anterior limit of the bone in lateral view and is located in a high position on the greater trochanter.
D. The attachment area of the *infraspinatus* muscle is well expanded beyond the anterior limit of the bone in lateral view and is located in a high position on the greater trochanter.

##### 14. Morphology of the lateral epicondyle in ventral view. (Fig. 9)

**Figure 9.**
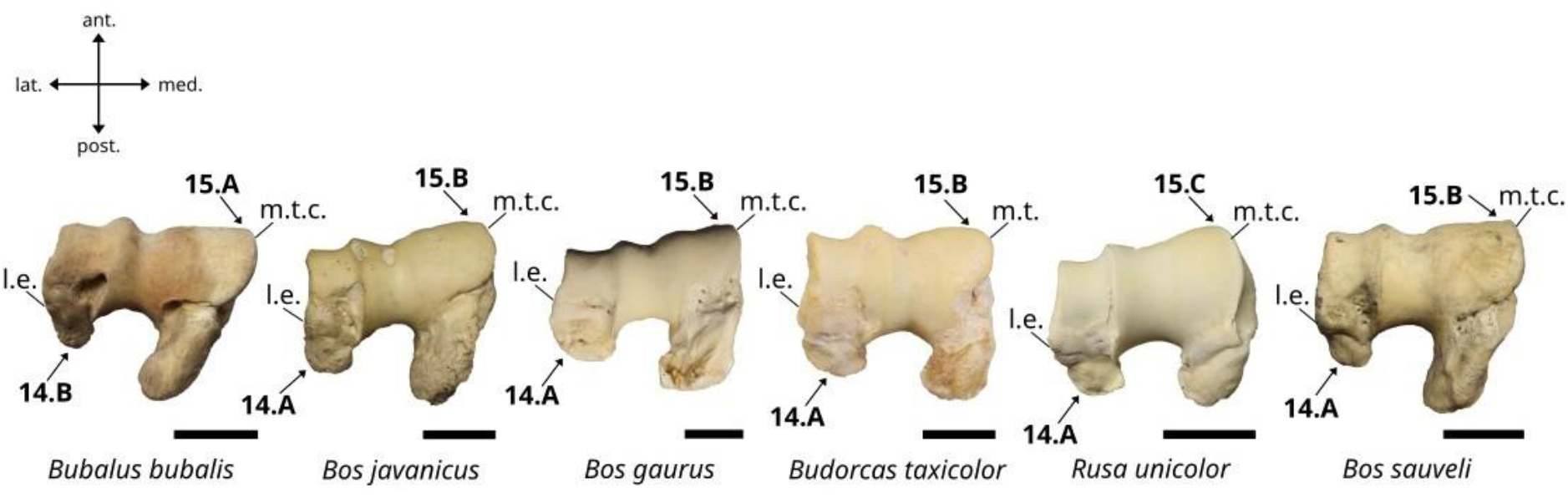
Character states visible on the distal face of the right humerus (characters 14 and 15). Abbreviations: m.t.c.: medial trochlea, l.e.: lateral epicondyle. Scalebar = 3 cm.

A. The lateral epicondyle is wide with a blunt extremity in ventral view.
B. The lateral epicondyle is slender with a more or less pointed posterior extremity in ventral view.

##### 15. Expansion of the medio-distal area of the trochlea in medial direction in ventral view. (Fig. 9)

A. Strongly expanded beyond the medial limit of the bone.
B. Moderately expanded beyond the medial limit of the bone.
C. Not expanded beyond the medial limit of the bone.

##### 16. Inclination of the lateral ridge of the trochlea in posterior view. (Fig. 10)

**Figure 10.**
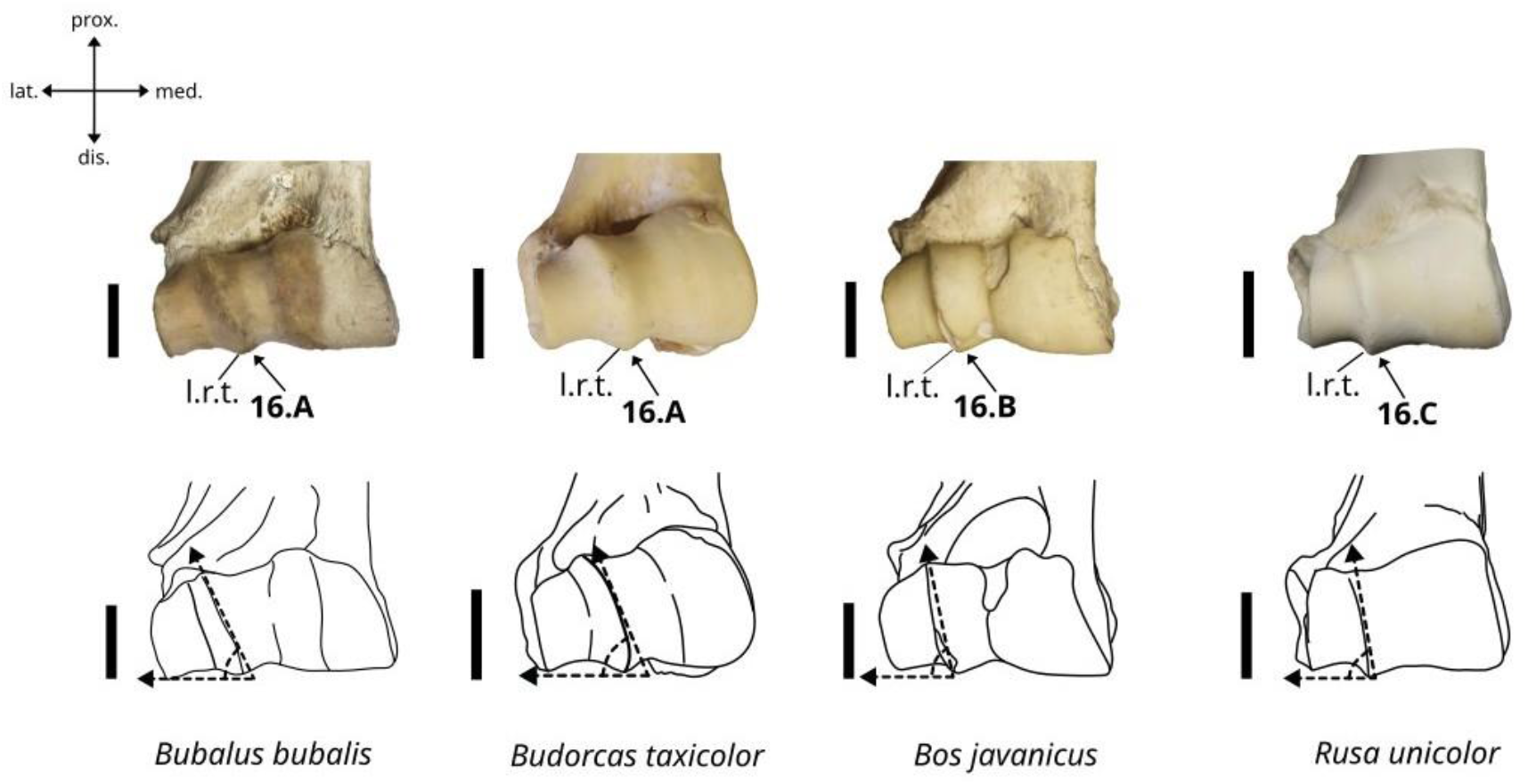
Character states visible on the anterior face of the right humerus (character 16). Abbreviation: l.r.t.: lateral ridge of trochlea. Scalebar = 3 cm.

A. The lateral ridge is strongly inclined in medial direction, at an angle of approximately 20° to 30°.
B. The lateral ridge is moderately inclined in medial direction, at an angle of approximately 20° to 10°.
C. The lateral ridge is weakly inclined in medial direction, appearing nearly straight, at an angle of less than 10°.

##### 17. Morphology of the minor trochanter in dorsal view. (Fig. 11)

**Figure 11.**
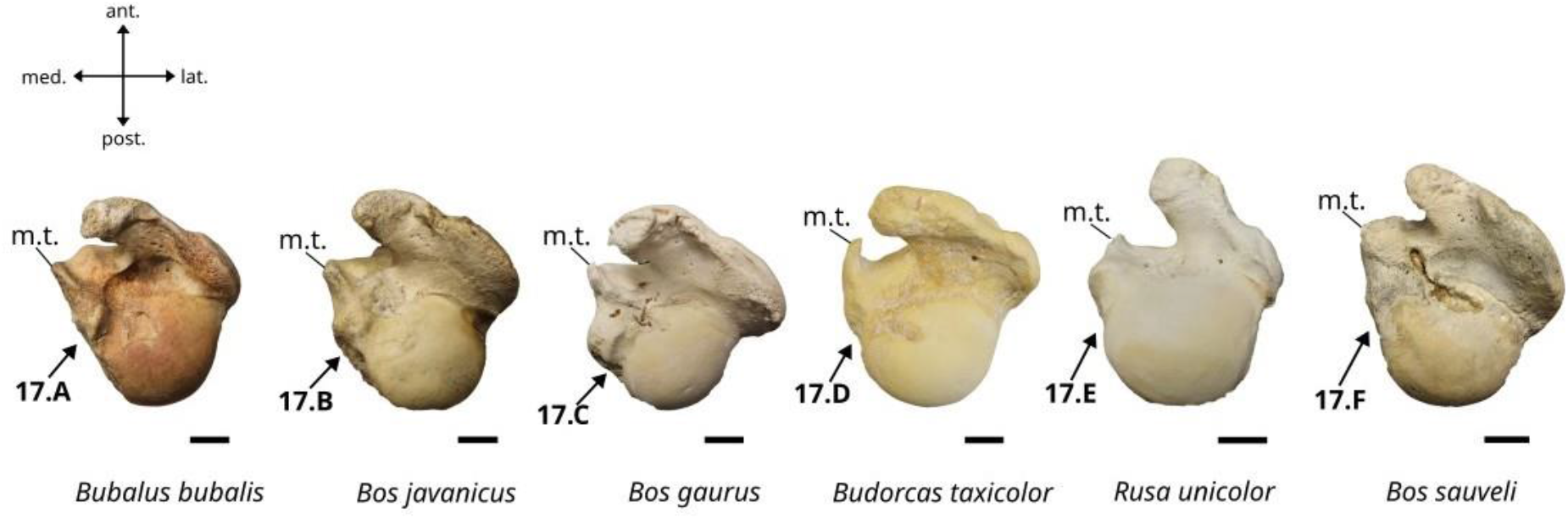
Character states visible on the proximal face of the right humerus (character 17). Abbreviation: m.t.: minor trochanter. Scalebar = 2 cm.

A. The minor trochanter is wide, well-developed, and weakly expanded in posterior direction. Its medial border is oblique.
B. The minor trochanter is wide, well-developed, and weakly expanded in posterior direction. Its medial border is slightly convex.
C. The minor trochanter is wide, well-developed, and expanded in medial direction. Its medial border is slightly concave.
D. The minor trochanter is wide, moderately developed, and presents a thin, elongated posterior projection. Its medial border is slightly concave.
E. The minor trochanter is poorly developed, with its anterior area strongly reduced, and presents a small, pointed posterior projection. Its medial border is slightly concave.
F. The minor trochanter is moderately developed and weakly expanded in medial and posterior directions. Its medial border is oblique.

##### 18. Expansion of the medial epicondyle in lateral view. (Fig. 12)

**Figure 12.**
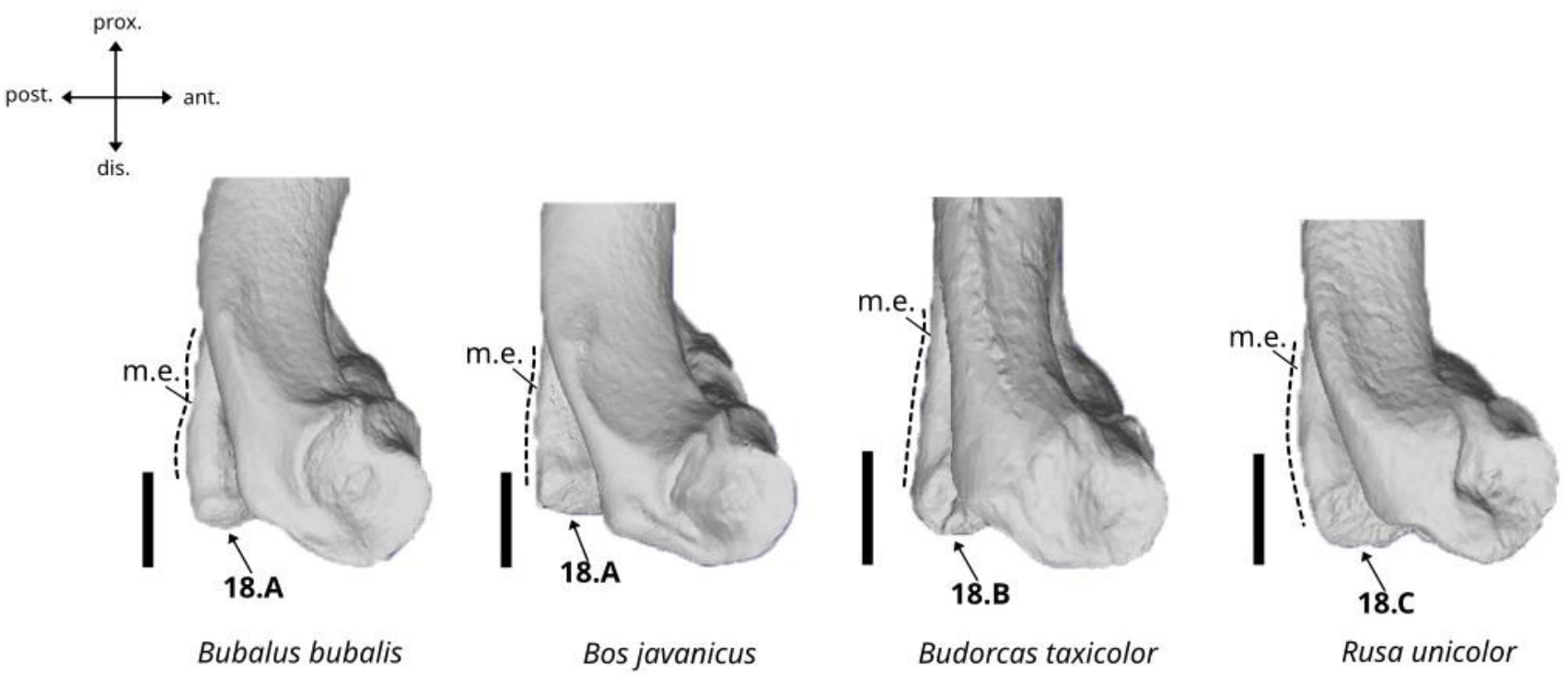
Character states visible on the lateral face of the right humerus (character 18). Abbreviation: m.e.: medial epicondyle. Scalebar = 3 cm.

A. The medial epicondyle is moderately expanded in distal direction, and its posterior border is sinusoidal or curved.
B. The medial epicondyle is moderately expanded in distal direction, and its posterior border is flat and straight.
C. The medial epicondyle is strongly expanded in distal direction, beyond the medio-distal limit of the condyle, and its posterior border is curved.

#### Characters of the ulna

##### 19. Shape of the dorsal margin of the olecranon in lateral view. (Fig. 13)

**Figure 13.**
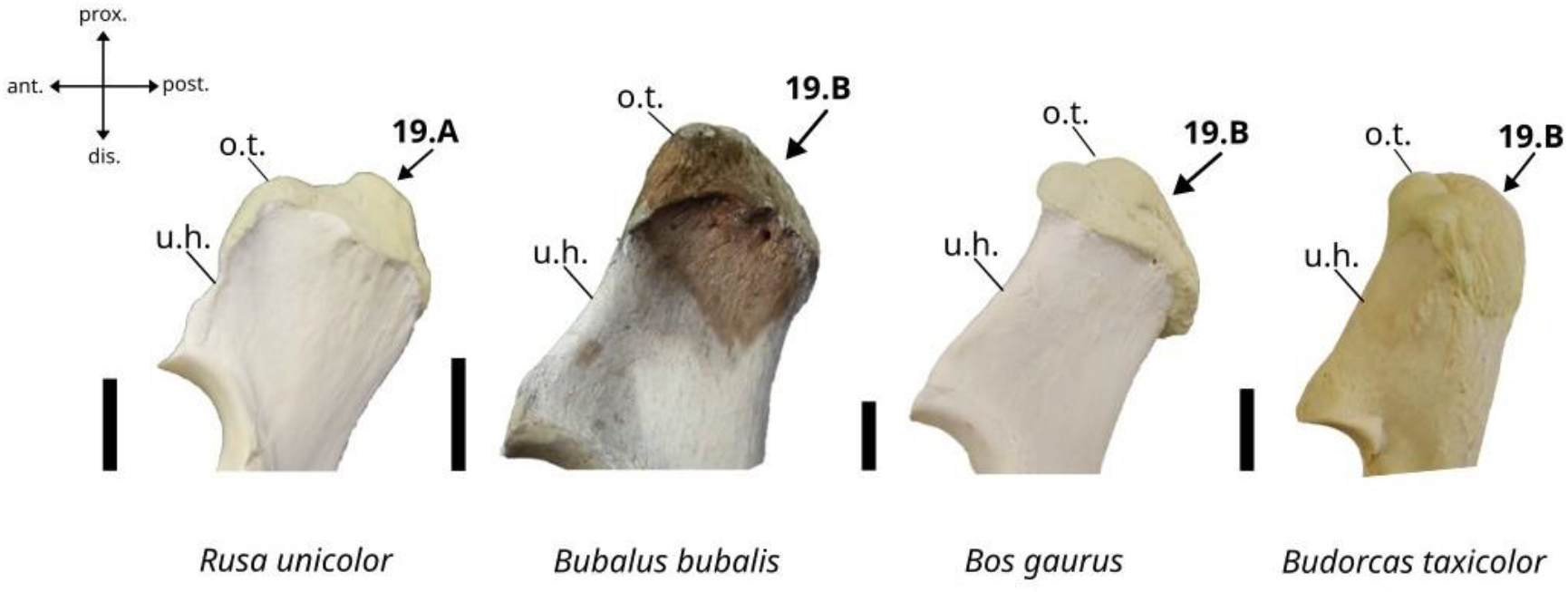
Character states visible on the lateral face of the right ulna (character 19). Abbreviations: o.t.: olecranon tuberosity, u.h.: ulna’s head. Scalebar = 3 cm.

A. It is sinusoidal with a deep notch.
B. It is more or less regularly curved.

##### 20. Development of the lateroventral articular expansion forming a part of the dorsal articular surface of the radius. (Fig. 14)

**Figure 14.**
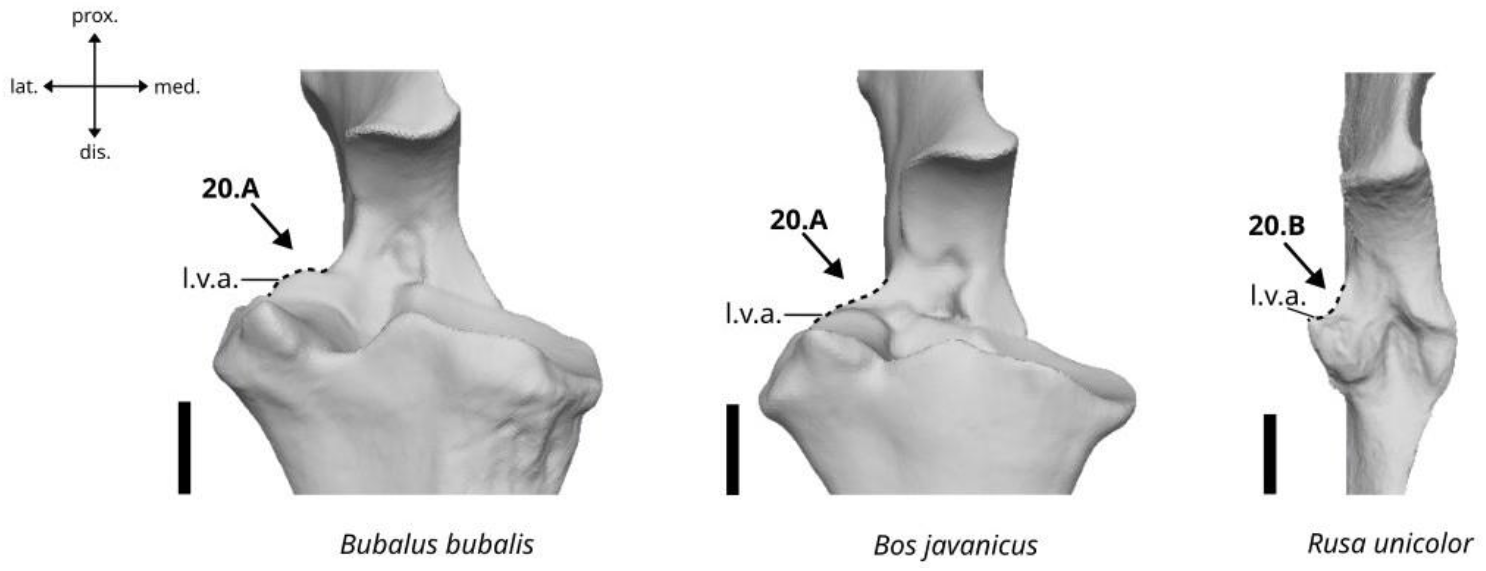
Character states visible on the anterior face of the right ulna (character 20). Abbreviation: l.v.a.: lateroventral articular surface. Scalebar = 2 cm.

A. The articular surface is well-developed and well-expanded beyond the lateral limit of the main body of the bone in anterior view.
B. The articular surface is poorly developed and weakly expanded beyond the lateral limit of the main body of the bone in anterior view.

##### 21. Development of the ulna’s styloid process in distal view. (Fig. 15)

**Figure 15.**
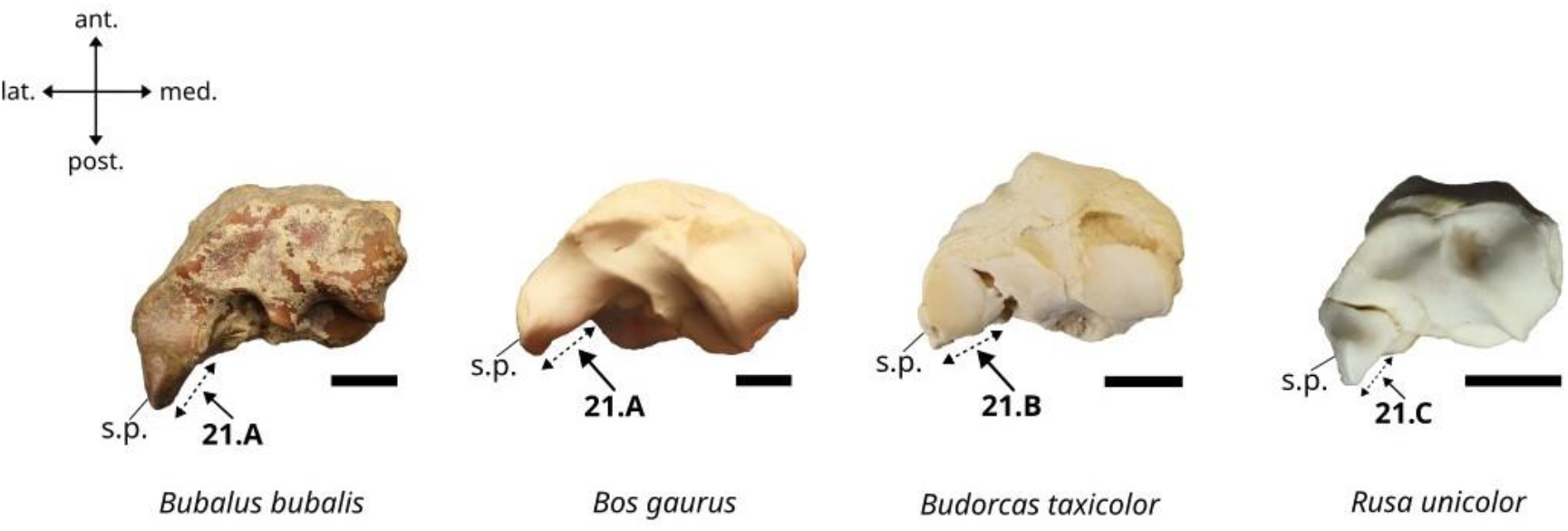
Character states visible on the distal face of the right ulna (character 21). Abbreviation: s.p.: styloid process of ulna. Scalebar = 2 cm.

A. Well-developed.
B. Moderately developed.
C. Poorly developed.

##### 22. Outline of the postero-dorsal margin in lateral view. (Fig. 16)

**Figure 16.**
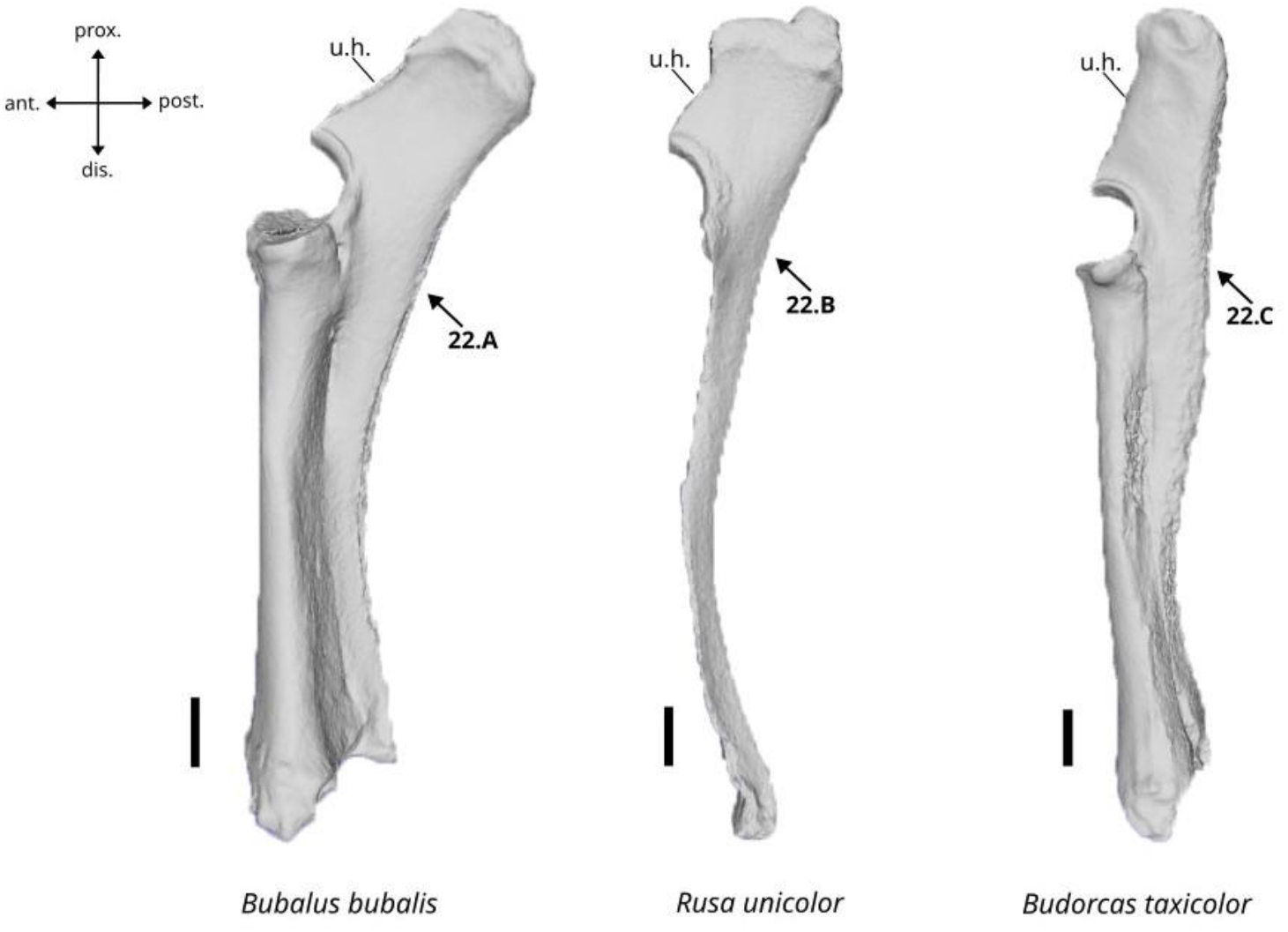
Character states visible on the lateral face of the right ulna (character 22). Abbreviation: u.h.: ulna’s head. Scalebar = 3 cm.

A. Strongly inclined in posterior direction.
B. Moderately inclined in posterior direction.
C. Not inclined in posterior direction and nearly straight.

#### Characters of the radius

##### 23. Presence of a postero-dorsal groove of the radius on the proximal articulation surface. (Fig. 17)

**Figure 17.**
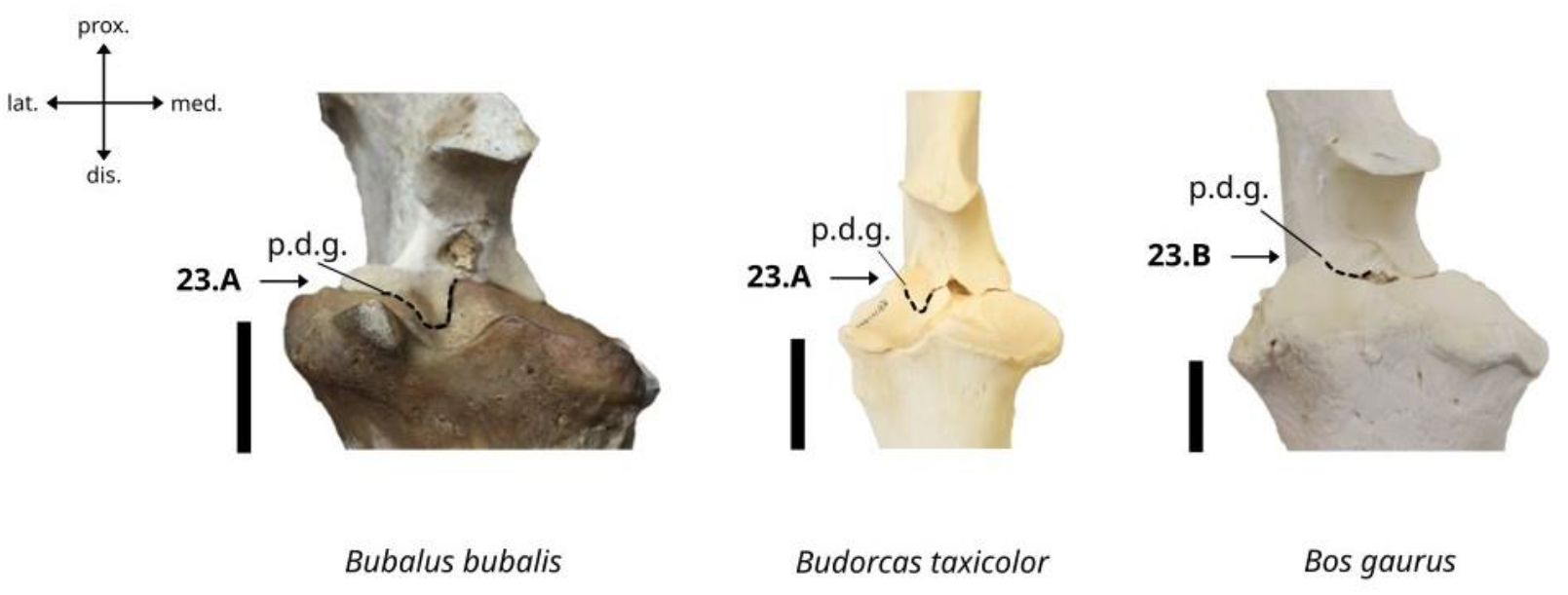
Character states visible on the anterior face of the right ulna (character 23). Abbreviation: p.d.g.: postero-dorsal groove of the radius. Scalebar = 3 cm.

A. The groove is present and well-extended.
B. The groove is either very small or absent.

##### 24. Morphology of the articular surface of the radius in distal view. (Fig. 18)

**Figure 18.**
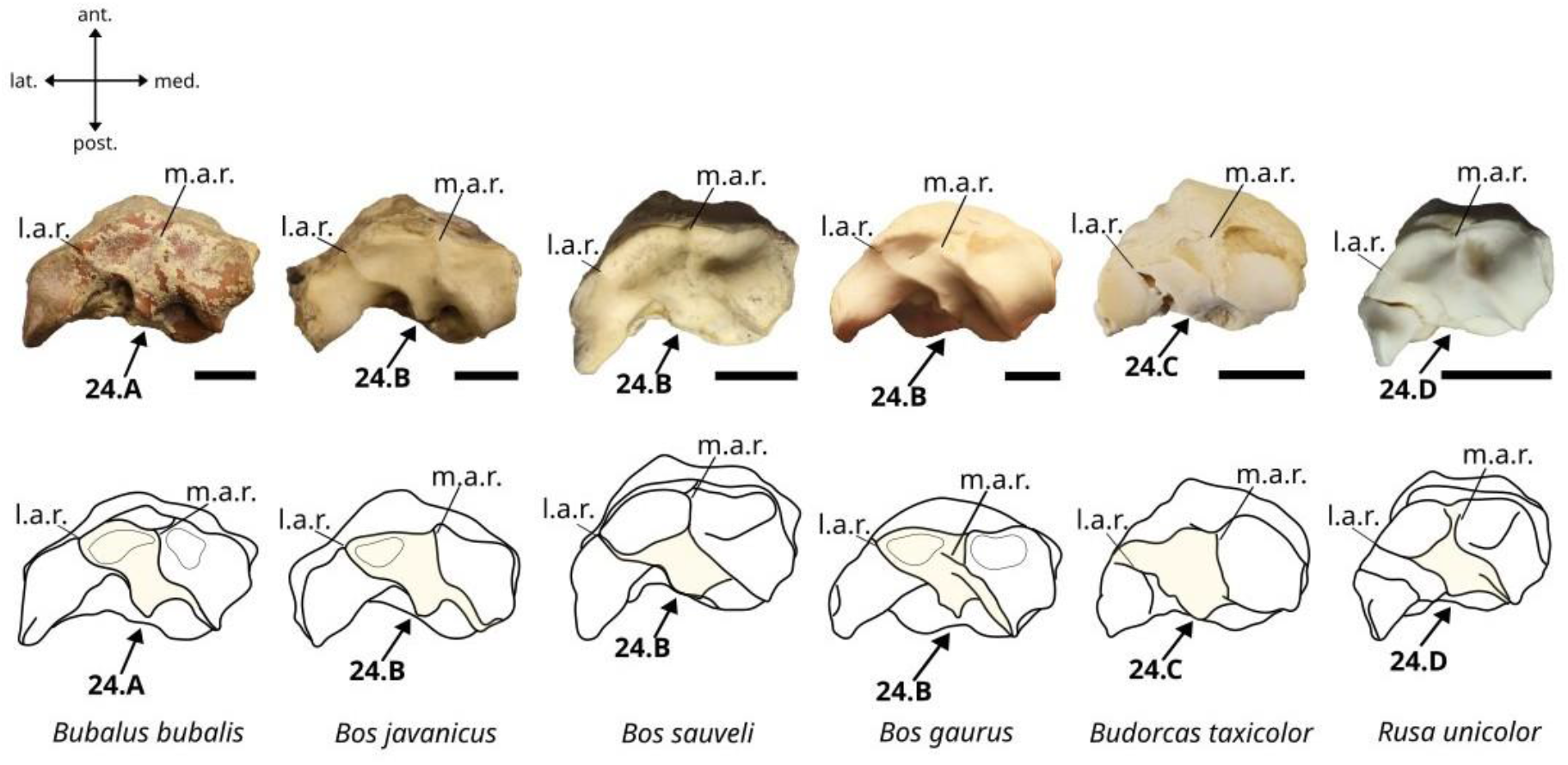
Character states visible on the distal face of the right radius (character 24). Abbreviations: m.a.r.: medial articular ridge, l.a.r.: lateral articular ridge. Scalebar = 3 cm.

A. The lateral and middle articular ridges are both sinusoidal.
B. The lateral articular ridge is sinusoidal, and the middle articular ridge is more or less uniformly curved and inclined in postero-lateral direction.
C. The lateral articular ridge is more or less uniformly curved and inclined in postero-lateral direction, and the middle articular ridge is nearly straight and oriented in antero-posterior direction.
D. The lateral articular ridge is more or less uniformly curved and inclined in postero-lateral direction, and the middle articular ridge is sinusoidal.

##### 25. Morphology of the proximal articular surface of the radius in the proximal view. (Fig. 19)

**Figure 19.**
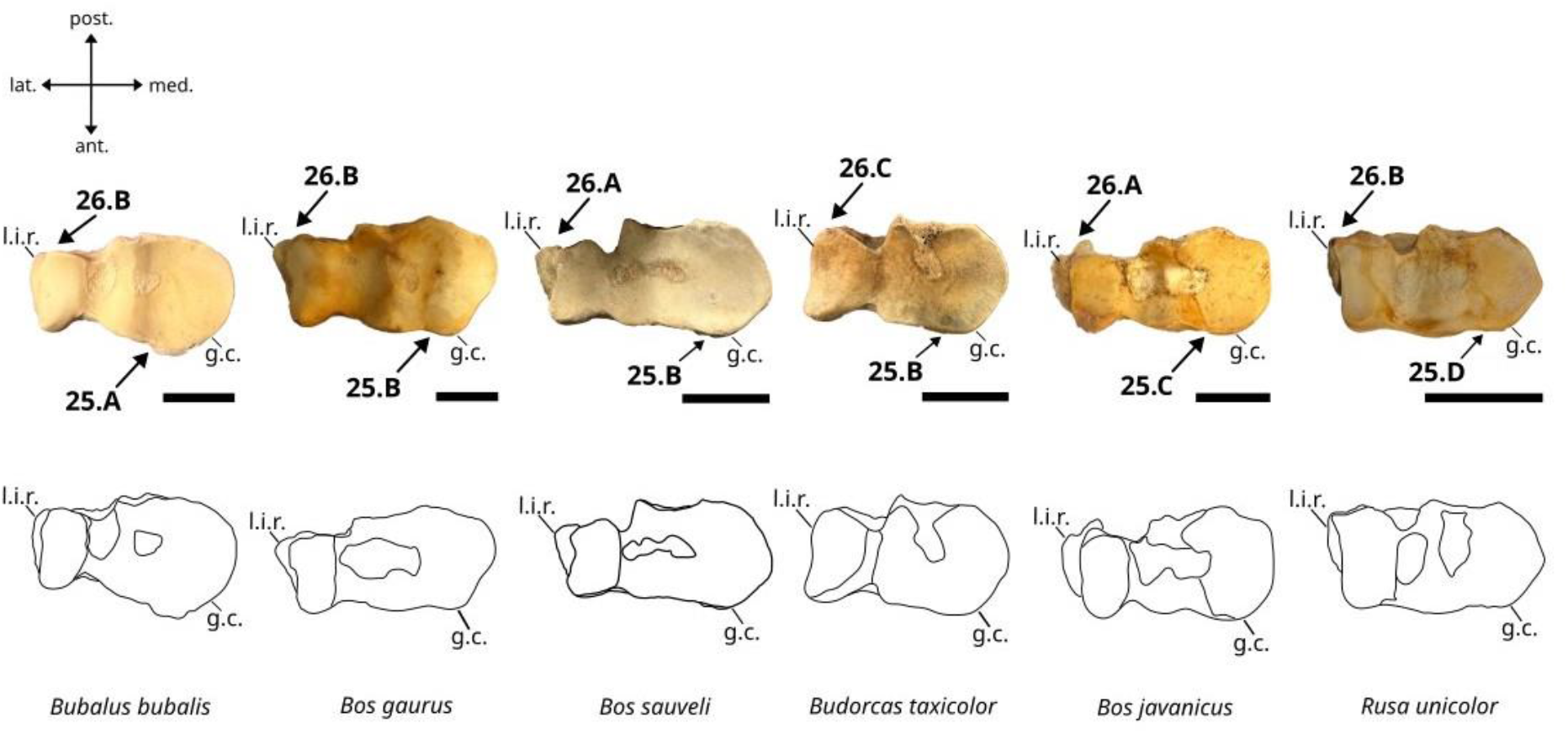
Character states visible on the proximal face of the right radius (character 25 and 26). Abbreviations: l.i.r.: lateral insertion relief, g.c.: glenoid cavity. Scalebar = 3 cm.

A. The glenoid cavity is strongly expanded in anteromedial direction.
B. The glenoid cavity is well expanded in anteromedial direction.
C. The glenoid cavity is weakly expanded in anteromedial direction.
D. The glenoid cavity is not expanded in anteromedial direction.

##### 26. Development of the lateral insertion relief in proximal view. (Fig. 19)

A. It is well developed.
B. It is weakly developed.
C. It is nearly absent.

#### Characters of the scaphoid bone

##### 27. Morphology of the scaphoid bone in lateral view. (Fig. 20)

**Figure 20.**
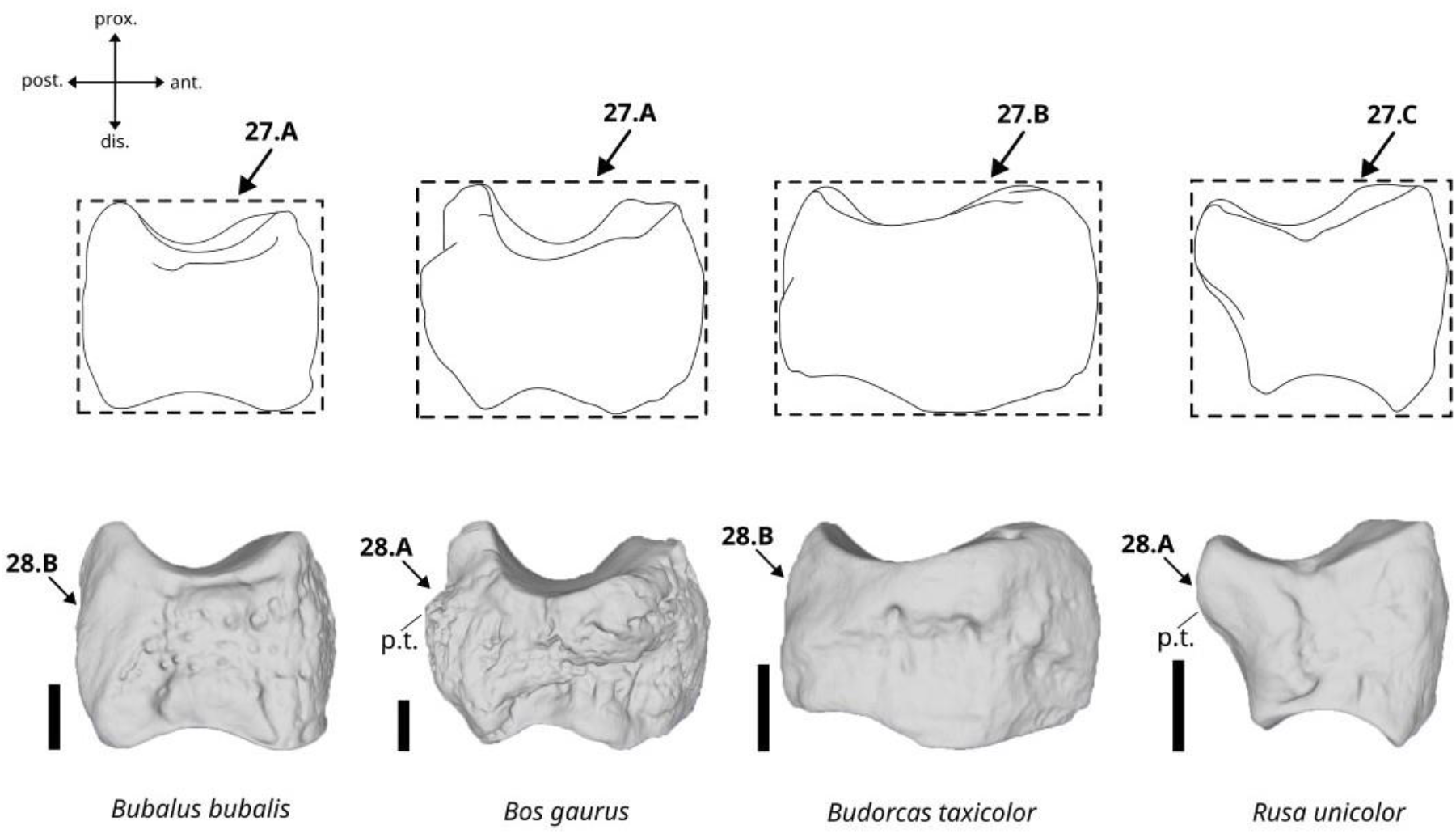
Character states visible on the lateral face of the left scaphoid (characters 27 and 28). Abbreviation: p.t.: posterior tuberosity. Scalebar = 1 cm.

A. It is slightly longer than high.
B. It is clearly longer than high.
C. It is as long as high.

##### 28. Occurrence of a posterior tuberosity in lateral view. (Fig. 20)

A. Present.
B. Absent.

##### 29. Morphology of the distal articular surface in medial view. (Fig. 21)

**Figure 21.**
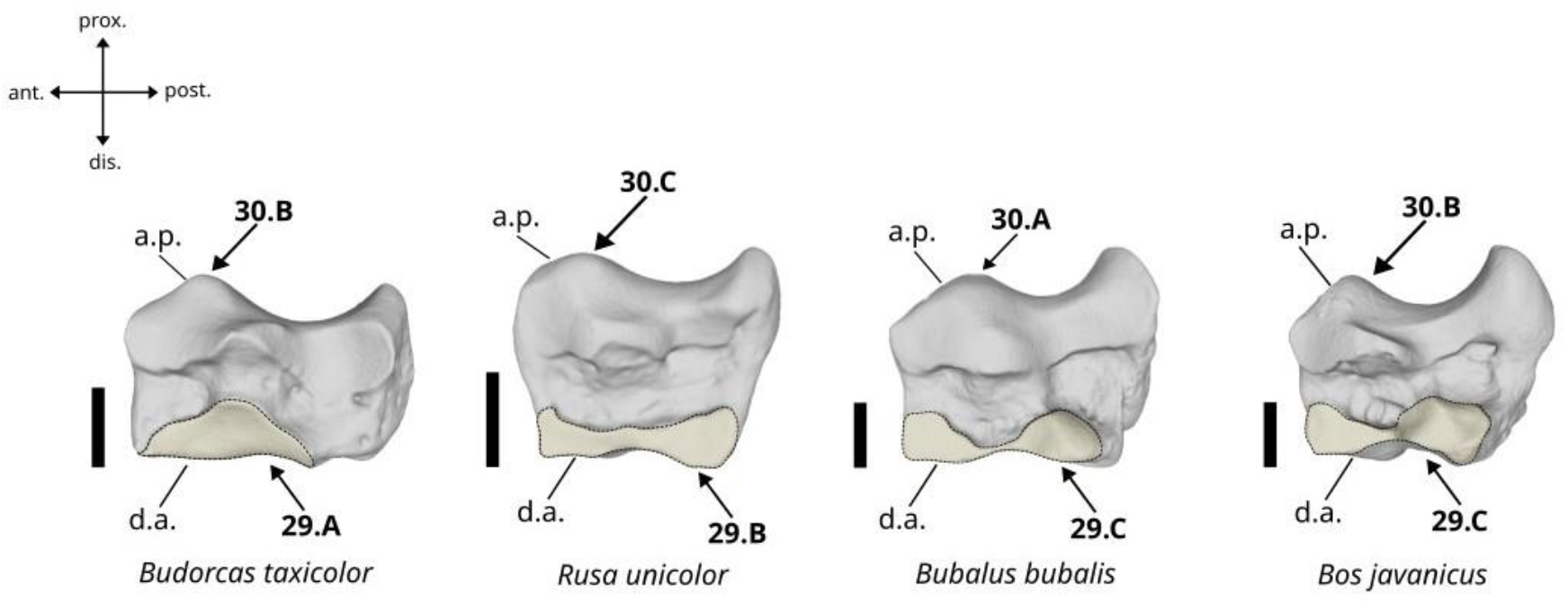
Character states visible on the medial face of the left scaphoid (characters 29 and 30). Abbreviations: a.p.: anterior prominence, d.a.: distal articular surface. Scalebar = 1 cm.

A. Present only on the anterior area of the medial surface of the bone.
B. Expanded evenly along the distal edge of the bone.
C. Expanded mainly in the anterior and posterior areas of the distal edge of the bone and reduced in its median area.

##### 30. Outline of the proximal articular surface in medial view. (Fig. 21)

A. The anterior prominence is triangular and weakly elevated, and the proximal outline of the bone is moderately curved.
B. The anterior prominence is triangular and well-elevated, and the proximal outline of the bone is strongly curved.
C. The anterior prominence is blunt and well-elevated, and the proximal outline of the bone is moderately curved.

##### 31. Shape of the anterior area of the medial border of the proximal articular surface. (Fig. 22)

**Figure 22.**
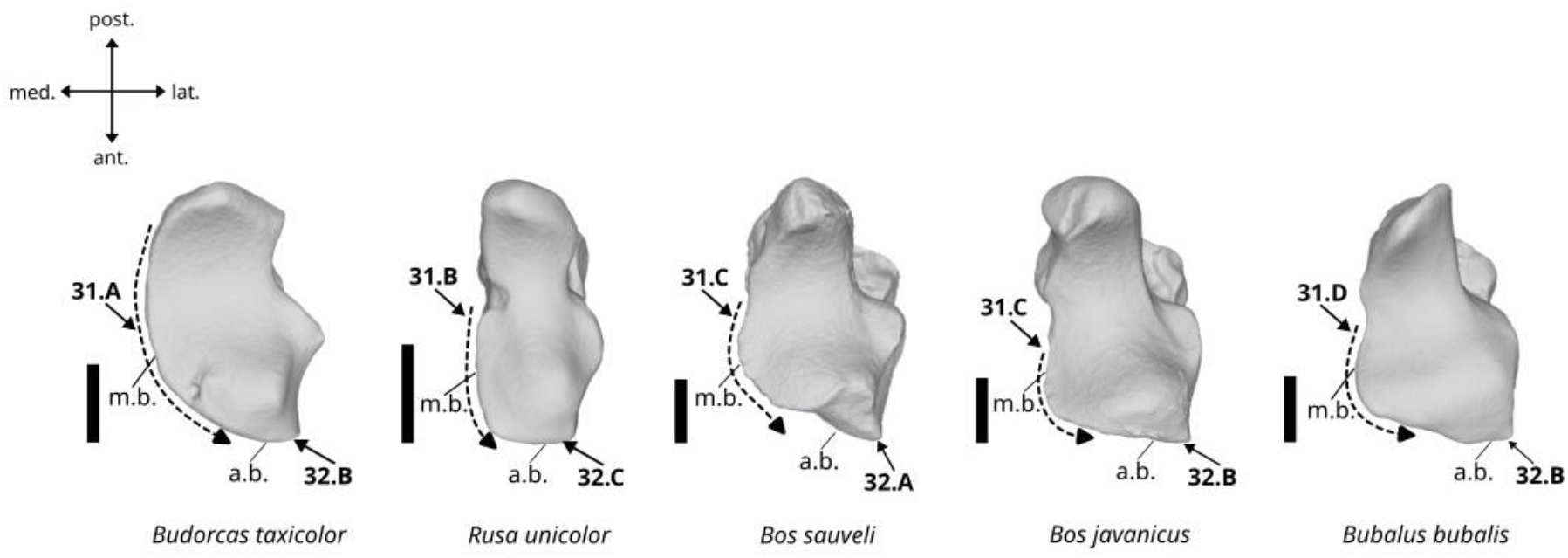
Character states visible on the proximal face of the left scaphoid (characters 31 and 32). Abbreviations: m.b.: medial border, a.b.: anterior border. Scalebar = 1 cm.

A. Medial border regularly convex throughout its length.
B. Medial border straight forming a 90° angle with the anterior border.
C. Medial border convex only in its anterior part with a convexity weakly expanded in medial direction.
D. Medial border convex only in its anterior part with a convexity well-expanded in medial direction.

##### 32. Morphology of the anterolateral angle of the articular surface in proximal view. (Fig. 22)

A. Prolonged into a pointed expansion directed anteriorly.
B. Slightly pointed.
C. Blunt.

##### 33. Shape of the anterior margin in distal view. (Fig. 23)

**Figure 23.**
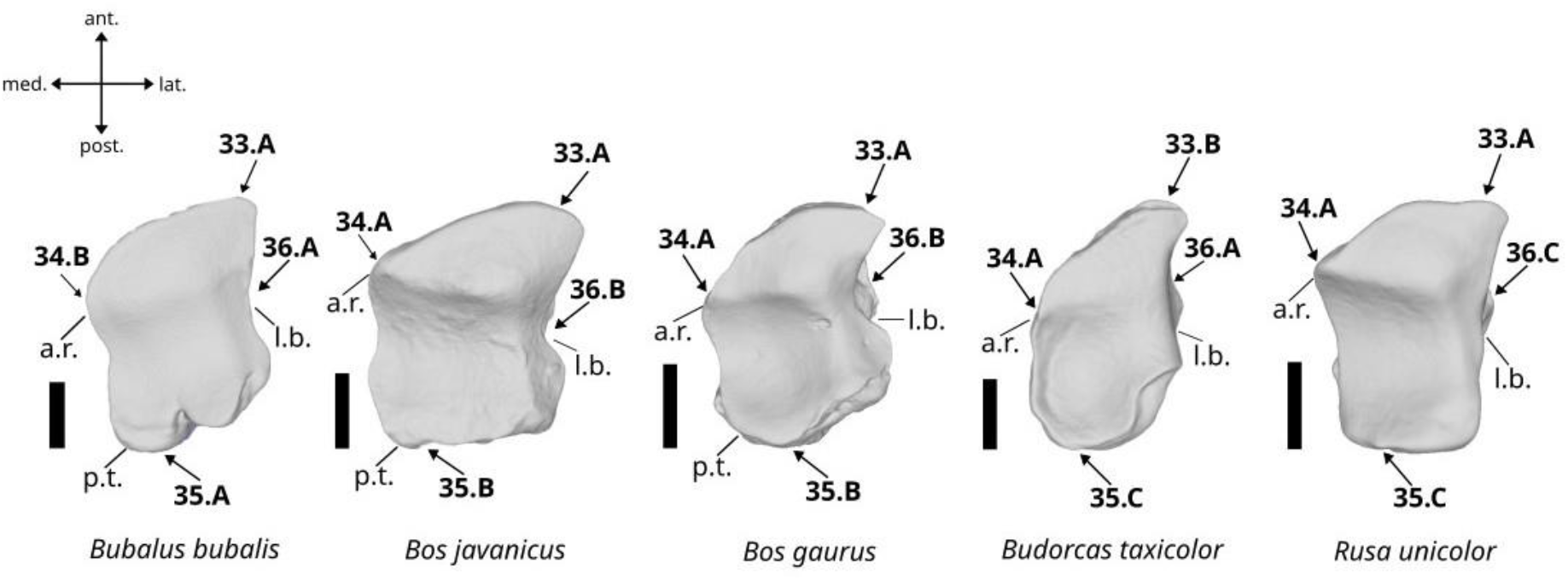
Character states visible on the distal face of the left scaphoid (characters 33, 34, 35, and 36). Abbreviations: l.b.: lateral border, a.r.: anterior ridge, p.t.: posterior tuberosity. Scalebar = 1 cm.

A. It is wide and convex.
B. It is thin and triangular.

##### 34. Development of the anterior ridge in distal view. (Fig. 23)

A. It is well-marked.
B. It is weakly marked.

##### 35. Morphology of the posterior tuberosity in distal view. (Fig. 23)

A. It is well-developed and strongly expanded in posterior direction.
B. It is weakly developed and weakly expanded in posterior direction.
C. It is absent.

##### 36. Shape of the lateral margin in distal view. (Fig. 23)

A. It is slightly concave.
B. It is clearly concave.
C. It is nearly straight.

#### Characters of the lunatum bone

##### 37. Morphology of the anterior border of the proximal articular surface in proximal view. (Fig. 24)

**Figure 24.**
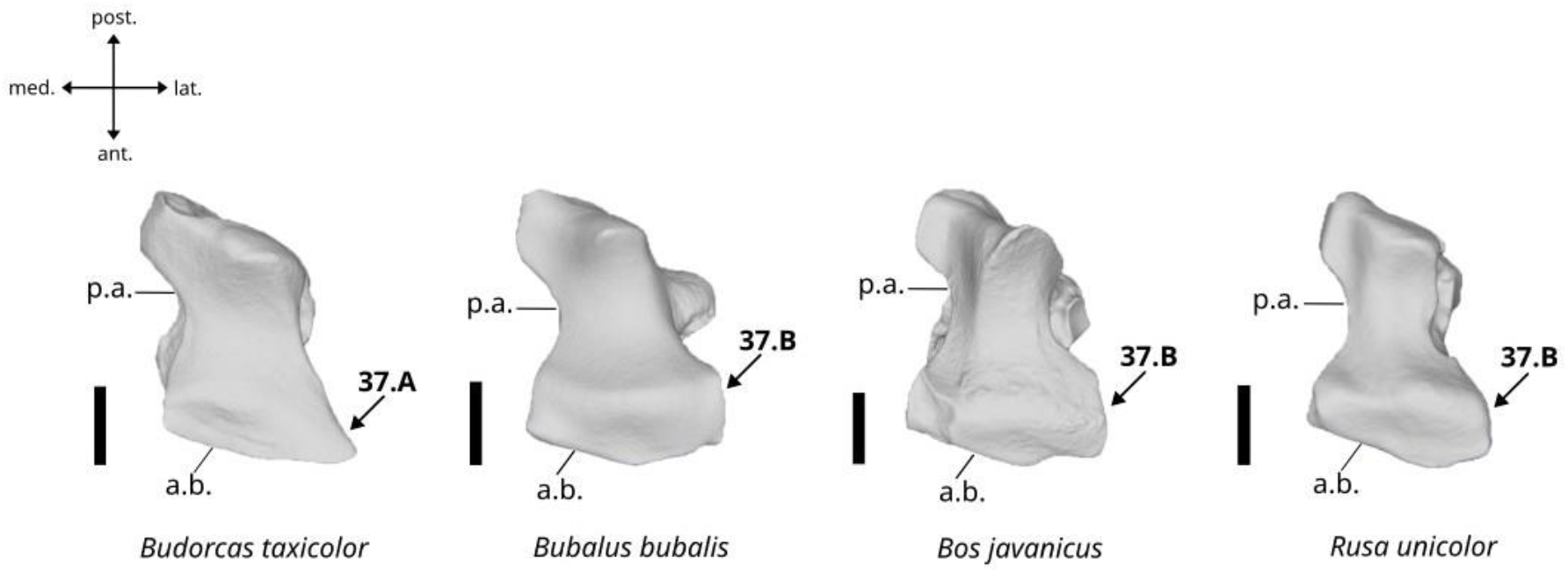
Character states visible on the proximal face of the left lunatum (character 37). Abbreviations: p.a.: proximal articular surface, a.b.: anterior border. Scalebar = 1 cm.

A. Oblique anterior border, with a pointed medial angle.
B. Oblique or irregular anterior border, with a truncated or curved medial angle.

##### 38. Morphology of the medial border of the distal articular surface in distal view. (Fig. 25)

**Figure 25.**
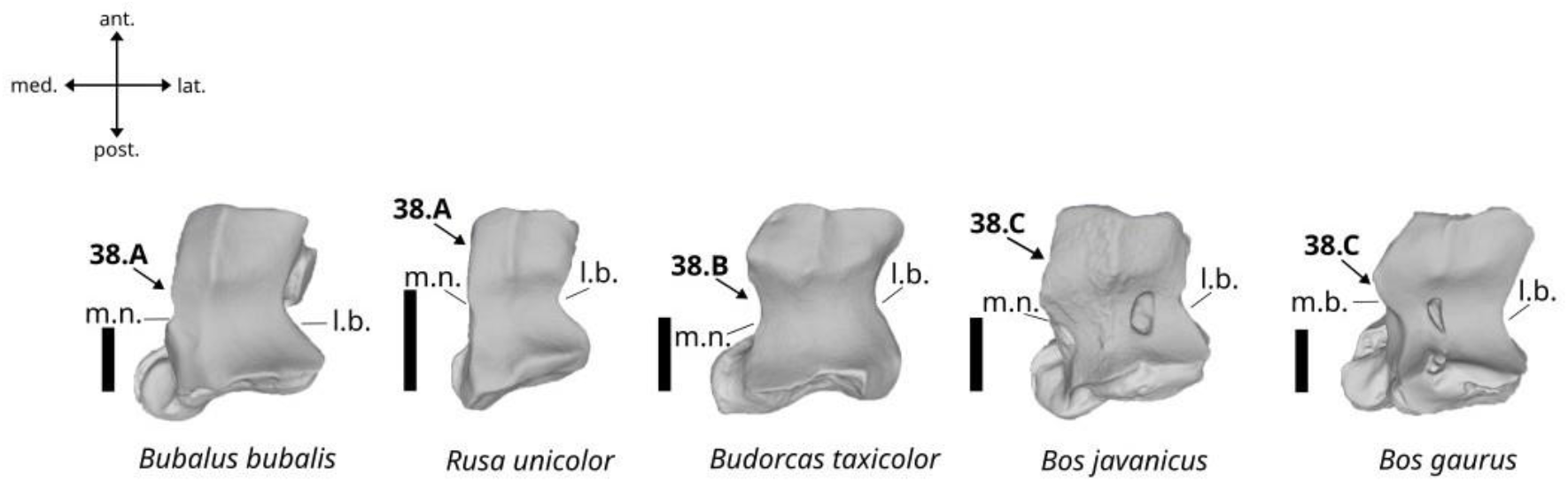
Character states visible on the distal face of the left lunatum (character 38). Abbreviations: m.n.: medial notch, l.b.: lateral border. Scalebar = 1 cm.

A. More or less straight without or with a very shallow medial notch.
B. Slightly sinusoidal with a shallow medial notch.
C. Clearly sinusoidal with a deep medial notch.

##### 39. Shape of the proximal articular surface for the scaphoid bone in medial view. (Fig. 26)

**Figure 26.**
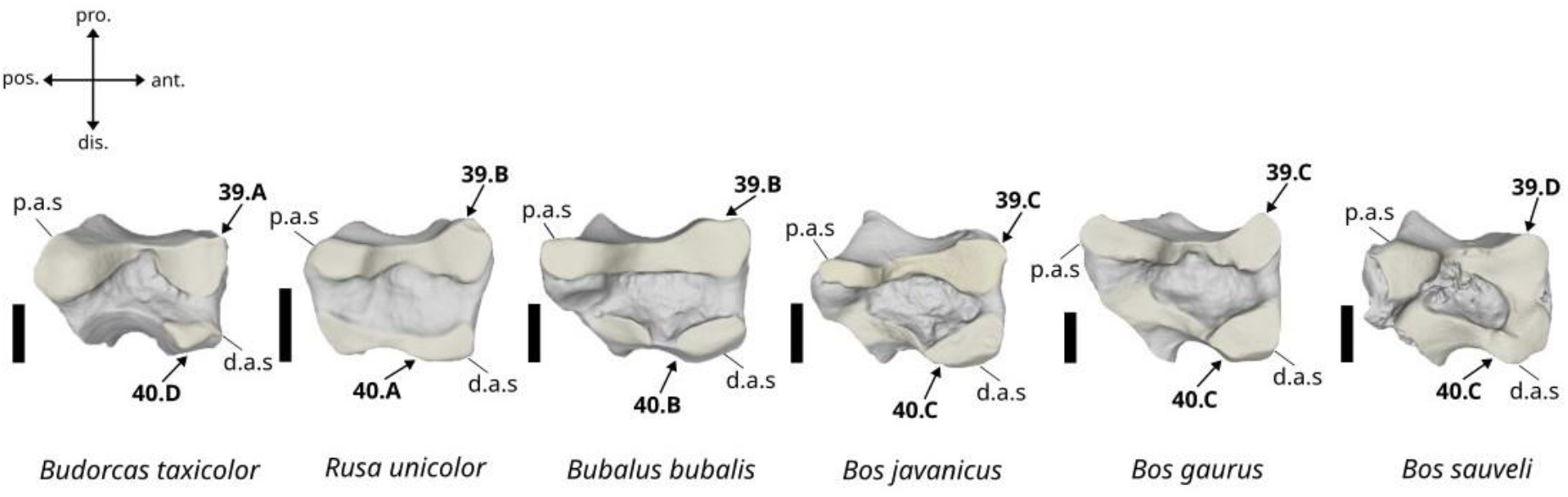
Character states visible on the medial face of the left lunatum (character 39 and 40). Abbreviations: p.a.s.: proximal articular surface for the scaphoid bone, d.a.s.: distal articular surface for the scaphoid bone. Scalebar = 1 cm.

A. Well-expanded in distal direction, forms two distinct proximal facets, and does not contact the distal articular surface.
B. Moderately expanded in distal direction, forms a single elongated proximal facet, and is not linked to the distal articular surface.
C. Weakly expanded in distal direction, forms a single elongated proximal facet, and is not linked to the distal articular surface.
D. Well-expanded in distal direction, forms two distinct proximal facets, and contacts the distal articular surface.

##### 40. Morphology of the distal articular surface of the scaphoid in medial view. (Fig. 26)

A. The articular surface occupies all the distal margin of the bone.
B. The articular surface is composed of an anterior and a posterior facet of similar size.
C. The articular surface is composed of a large posterior facet plunging in lateral direction and of a large anterior facet located in the anteroventral area.
D. The articular surface is reduced to a single small anterior facet.

#### Characters of the pyramidal bone

##### 41. Lateral inclination of the anterior margin of the proximal articular surface in proximal view. (Fig. 27)

**Figure 27.**
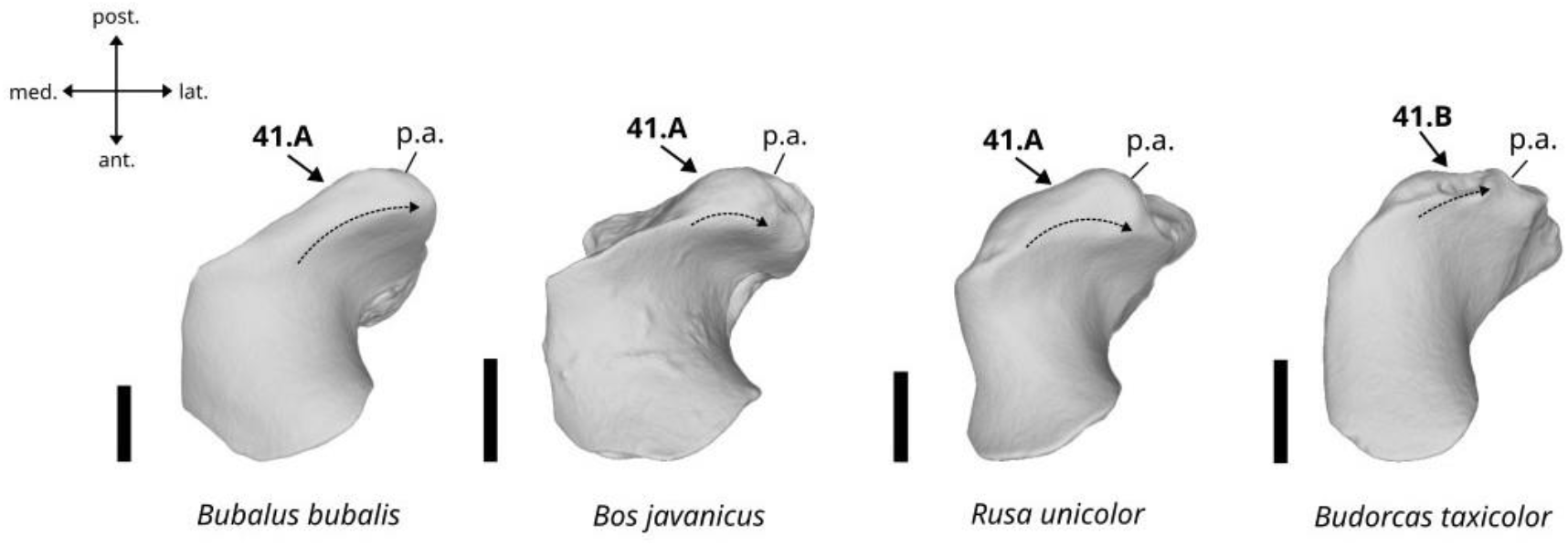
Character states visible on the proximal face of the left pyramidal (character 41). Abbreviation: p.a.: posterior articular surface. Scalebar = 1 cm.

A. Strongly inclined.
B. Weakly inclined.

##### 42. Morphology of the posterior border of the articular surface for the pisiform in lateral view. (Fig. 28)

**Figure 28.**
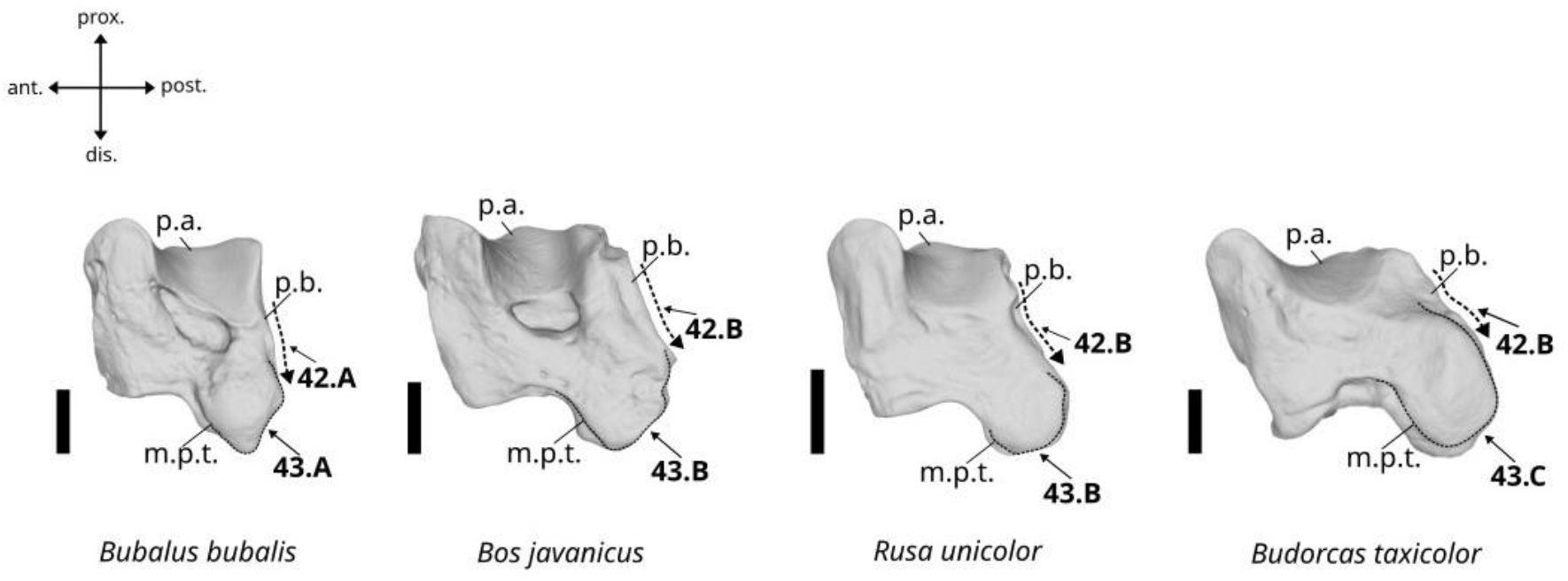
Character states visible on the lateral face of the left pyramidal (characters 42 and 43). Abbreviations: p.a.: proximal articular surface, p.b.: posterior border, m.p.t.: medio-posterior tuberosity. Scalebar = 1 cm.

A. Relatively straight, and not projected in posterior direction.
B. Incurved or sinusoidal, and more or less projected in posterior direction.

##### 43. Expansion of the posteroventral tuberosity of the distal articular surface in lateral view. (Fig. 28)

A. It is well expanded beyond the distal limit of the bone.
B. It is slightly expanded beyond the distal limit of the bone.
C. It does not expand beyond the distal limit of the bone.

##### 44. Morphology of the posteroventral tuberosity of the distal articular surface in medial view. (Fig. 29)

**Figure 29.**
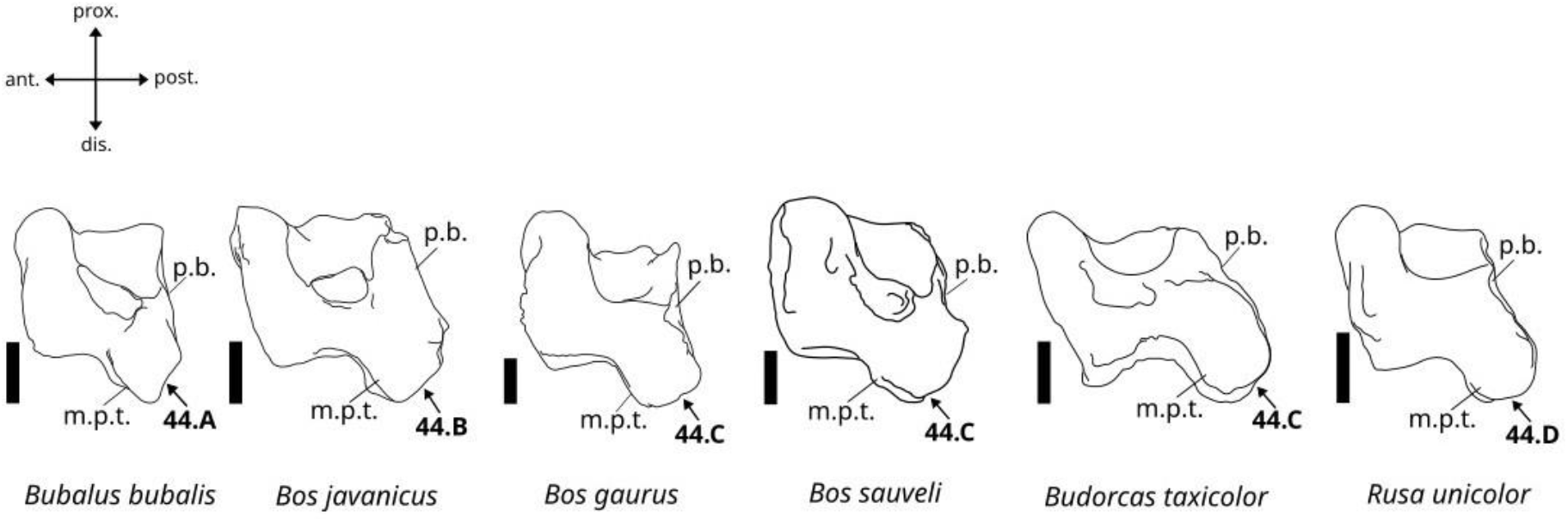
Character states visible on the lateral face of the left pyramidal (character 44). Abbreviations: p.b.: posterior border, m.p.t.: medio-posterior tuberosity. Scalebar = 1 cm.

A. The distal end of the tuberosity is triangular and pointed.
B. The distal end of the tuberosity is triangular with a blunt distal apex.
C. The distal end of the tuberosity is squared.
D. The distal end of the tuberosity is sub-circular.

##### 45. Proximal expansion of the distal articular surface in lateral view. (Fig. 30)

**Figure 30.**
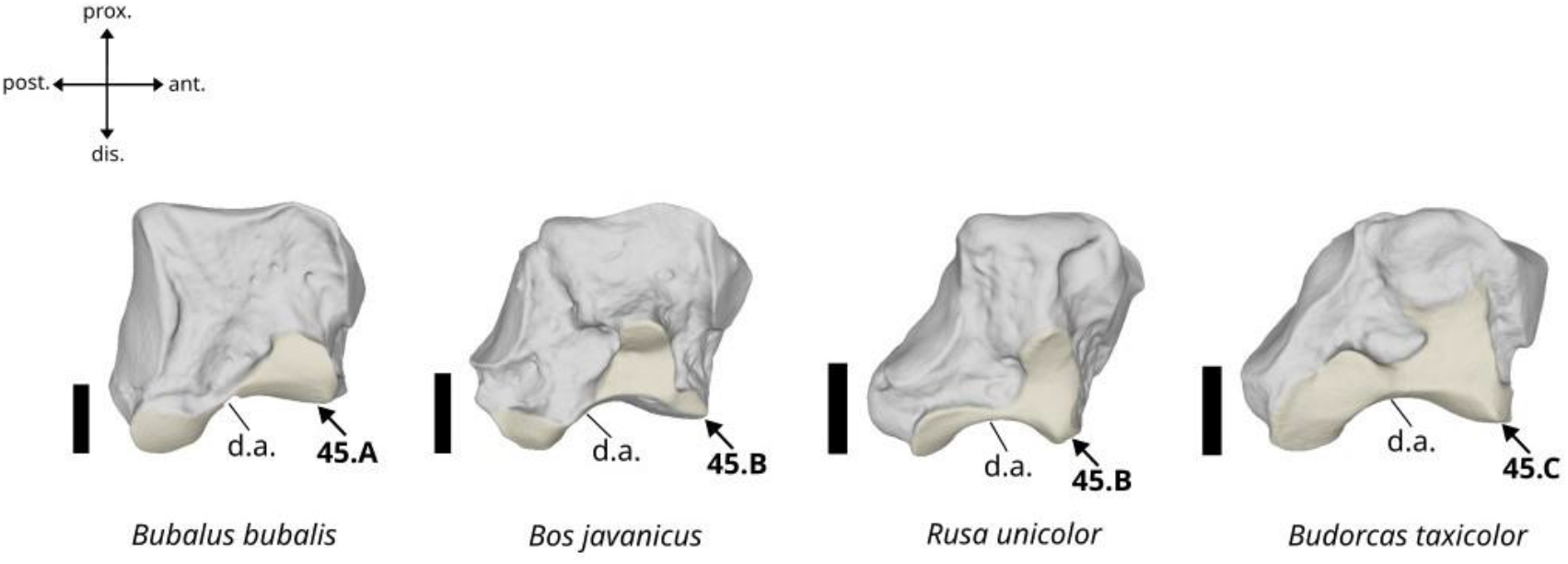
Character states visible on the medial face of the left pyramidal (character 45). Abbreviation: d.a.: distal articular surface. Scalebar = 1 cm.

A. The anterior area is weakly expanded in proximal direction, the posterior area is restricted to the distal part of the bone.
B. The anterior area is strongly expanded in proximal direction, the posterior area is restricted to the distal part of the bone.
C. Both the anterior and posterior areas are strongly expanded in proximal direction.

#### Characters of the pisiform bone

##### 46. Morphology of the pisiform bone in medial view. (Fig. 31)

**Figure 31.**
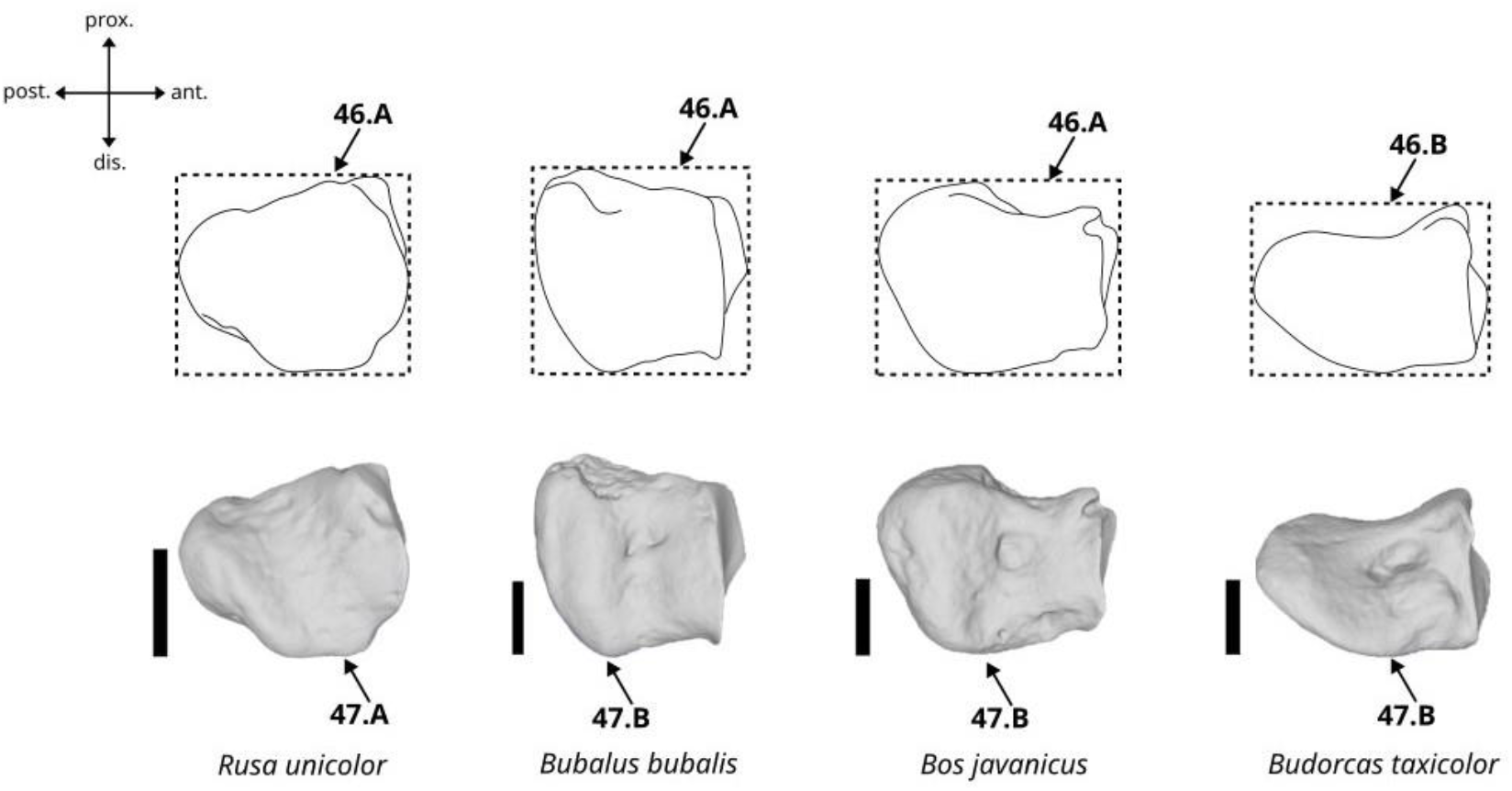
Character states visible on the medial face of the left pisiform (characters 46 and 47). Scalebar = 1 cm.

A. Short.
B. Elongate.

##### 47. Morphology of the distal border in medial view. (Fig. 31)

A. It presents a medio-distal projection and is sinusoidal.
B. It is more or less homogeneously curved.

##### 48. Morphology of the anterior articular surface in anterior view. (Fig. 32)

**Figure 32.**
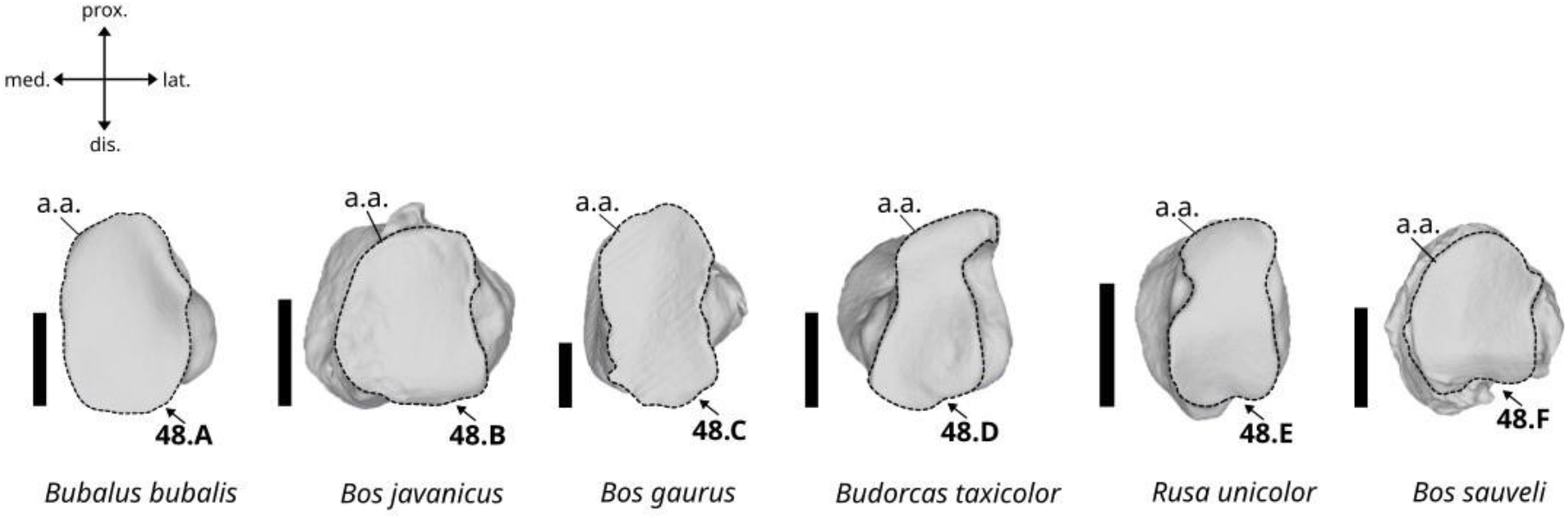
Character states visible on the anterior face of the left pisiform (character 48). Abbreviation: a.a.: anterior articular surface. Scalebar = 1 cm.

A. The articular surface is oval in shape. The lateral border presents a small triangular expansion pointing in lateral direction.
B. The articular surface is semi-circular. The lateral border presents a shallow median notch.
C. The articular surface is oval and narrow. The lateral border presents a sinusoidal margin.
D. The articular surface is hexagonal and elongated. The lateral border presents a deep notch in its proximal part, giving the articular surface a beak shape in this area.
E. The articular surface is oval and narrow in shape. The lateral and medial borders present well-marked notches.
F. The articular surface is sub-circular. The medial border is not expanded in medial direction, and the lateral and distal borders present small notches in their median areas.

#### Characters of the magnum bone

##### 49. Morphology of the posteromedial border of the articular surface for the metacarpus in distal view. (Fig. 33)

**Figure 33.**
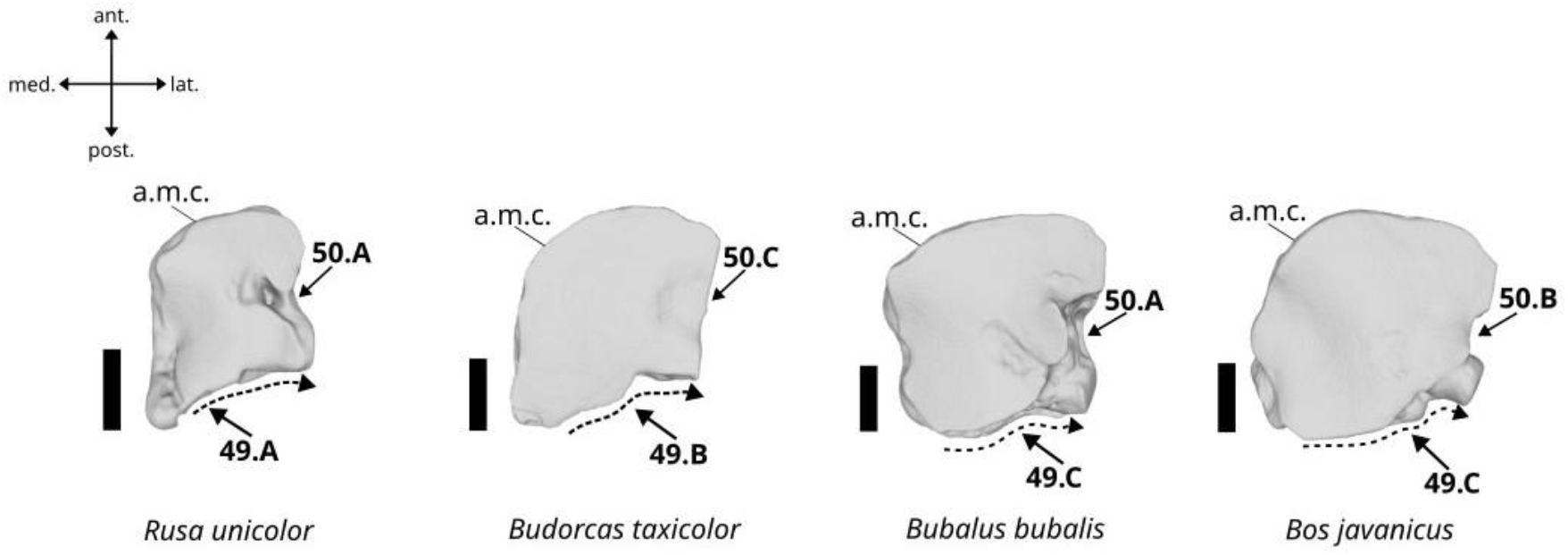
Character states visible on the distal face of the left magnum (characters 49 and 50). Abbreviations: a.m.c.: articular surface for the metacarpus (distal). Scalebar = 1 cm.

A. It is strongly projected in posterior direction and forms a moderately elongated expansion with a blunt apex.
B. It is strongly projected in posterior direction and is of wide, truncated triangular shape.
C. It is weakly projected in posterior direction, with a more or less uniformly curved margin.

##### 50. Occurrence of a notch in the medial margin of the articular surface for the metacarpus in distal view. (Fig. 33)

A. Occurrence of a deep notch.
B. Occurrence of a shallow notch.
C. Absence of notch.

##### 51. Depth of the proximal margin of the proximal articular surface in posterior view. (Fig. 34)

**Figure 34.**
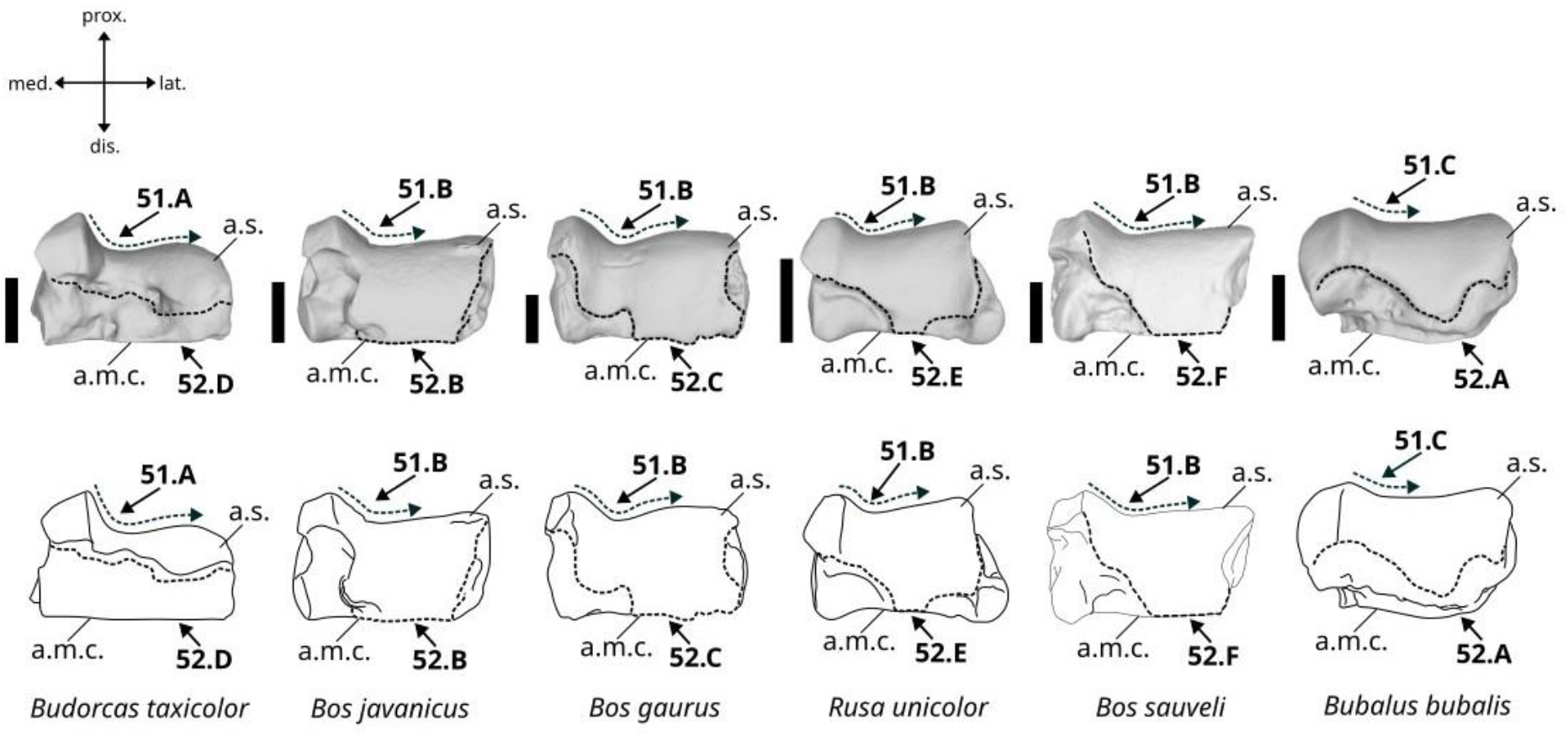
Character states visible on the posterior face of the left magnum (characters 51 and 52). Abbreviations: a.s.: articular surface for the scaphoid (proximal), a.m.c.: articular surface for the metacarpus (distal). Scalebar = 1 cm.

A. Very deep.
B. Moderately deep.
C. Shallow.

##### 52. Shape of the limits of the articular surface for the scaphoid in posterior view. (Fig. 34)

A. The articular surface is moderately expanded in distal direction and does not contact the articular surface for the metacarpus. It presents a pointed distal apex.
B. The articular surface is expanded in distal direction and contacts the articular surface for the metacarpus in all of its lateral half. The articular surface is moderately extended on the medioventral area of the bone, and the medial border of the articular surface is concave.
C. The articular surface is expanded in distal direction and contacts the articular surface for the metacarpus in all of its lateral half. The articular surface is well-extended on the medioventral area of the bone, and the medial border of the articular surface is sinusoidal.
D. The articular surface is weakly expanded in distal direction and does not contact the articular surface for the metacarpus.
E. The articular surface is expanded in distal direction and contacts the articular surface for the metacarpus. The contact area with the metacarpus is only made on a small median area, as the posterior and anterior areas of the distal border of the articular surface are concave and moderately expanded in distal direction.
F. The articular surface is expanded in distal direction and contacts the articular surface for the metacarpus in all of its lateral half. The articular surface does not extend on the medioventral area of the bone, and the medial border of the articular surface is nearly straight and oblique.

##### 53. Shape of the posterior area of the articular surface for the unciform bone in medial view. (Fig. 35)

**Figure 35.**
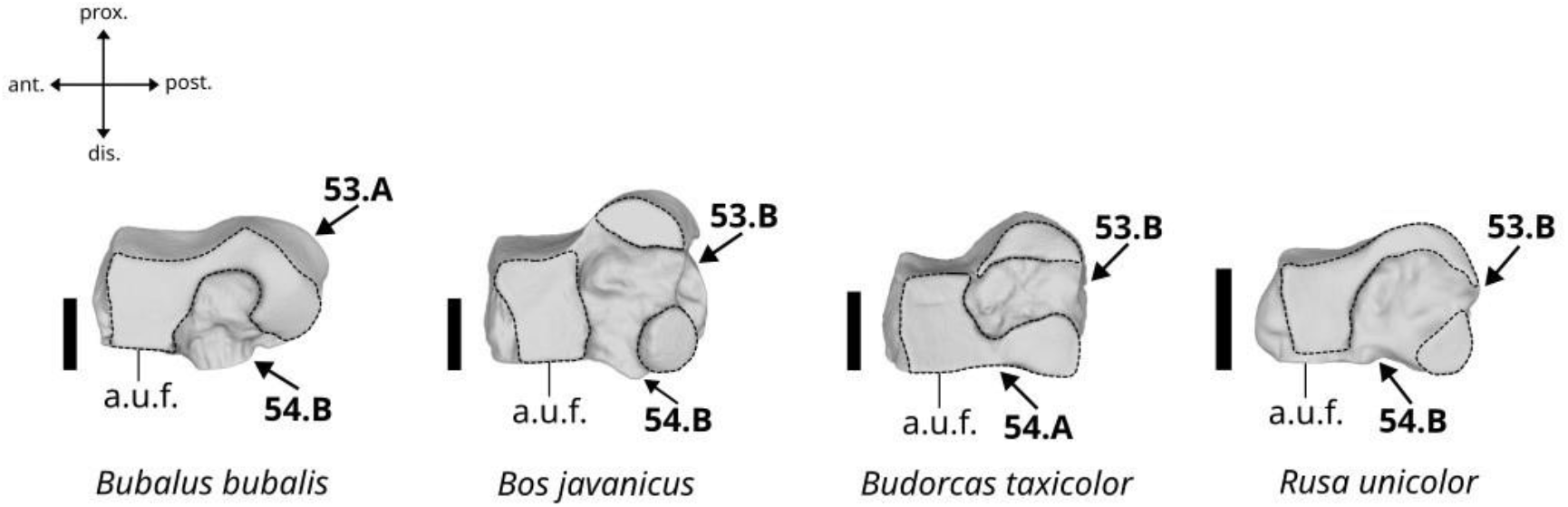
Character states visible on the lateral face of the left magnum (characters 53 and 54). Abbreviations: a.u.f.: articular surface for the unciform bone (medial). Scalebar = 1 cm.

A. Continuous.
B. Divided into two areas.

##### 54. Shape of the distal area of the articular surface for the unciform bone in medial view. (Fig. 35)

A. Continuous along the distal edge.
B. Divided into two areas.

#### Characters of the unciform bone

##### 55. Morphology of the posterior border of the articulation surface for the pisiform in proximal view. (Fig. 36)

**Figure 36.**
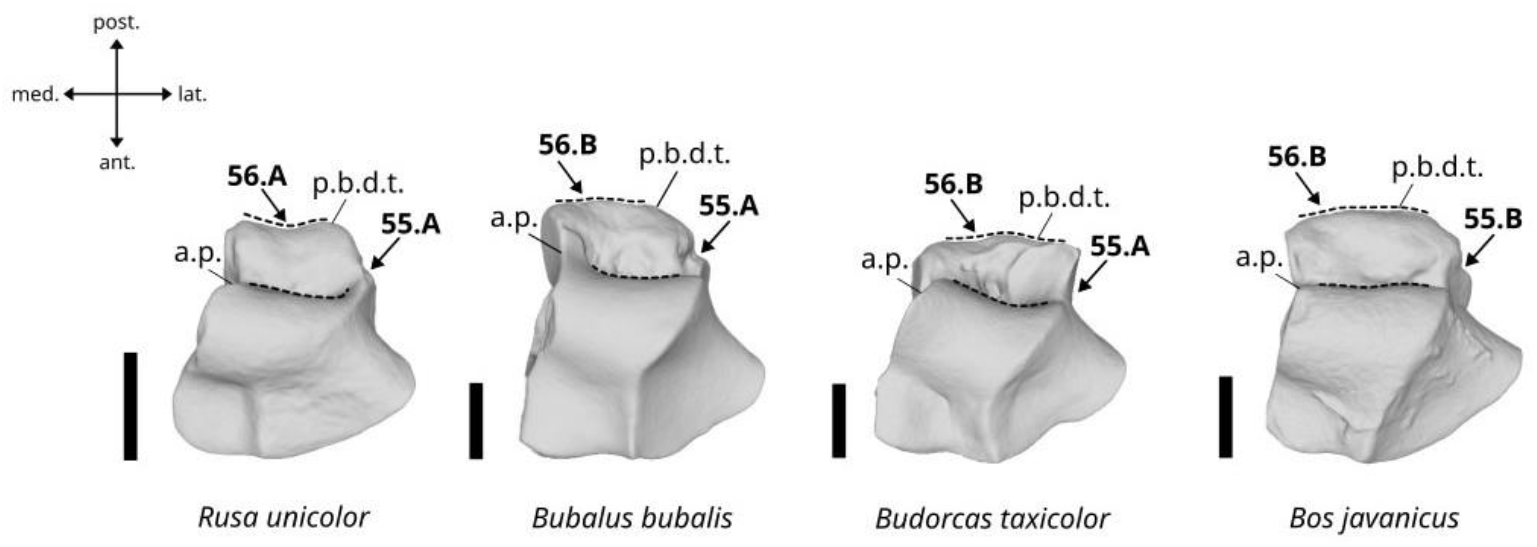
Character states visible on the proximal face of the left unciform (characters 55 and 56). Abbreviations: a.p.: articular surface for the pisiform, p.b.d.t.: posterior border of the distal tuberosity. Scalebar = 1 cm.

A. Concave.
B. More or less straight.

##### 56. Morphology of the posterior border of the distal tuberosity in proximal view. (Fig. 36)

A. Concave.
B. Straight or slightly convex.

##### 57. Morphology and orientation of the posterolateral apex of the distal articular surface in distal view. (Fig. 37)

**Figure 37.**
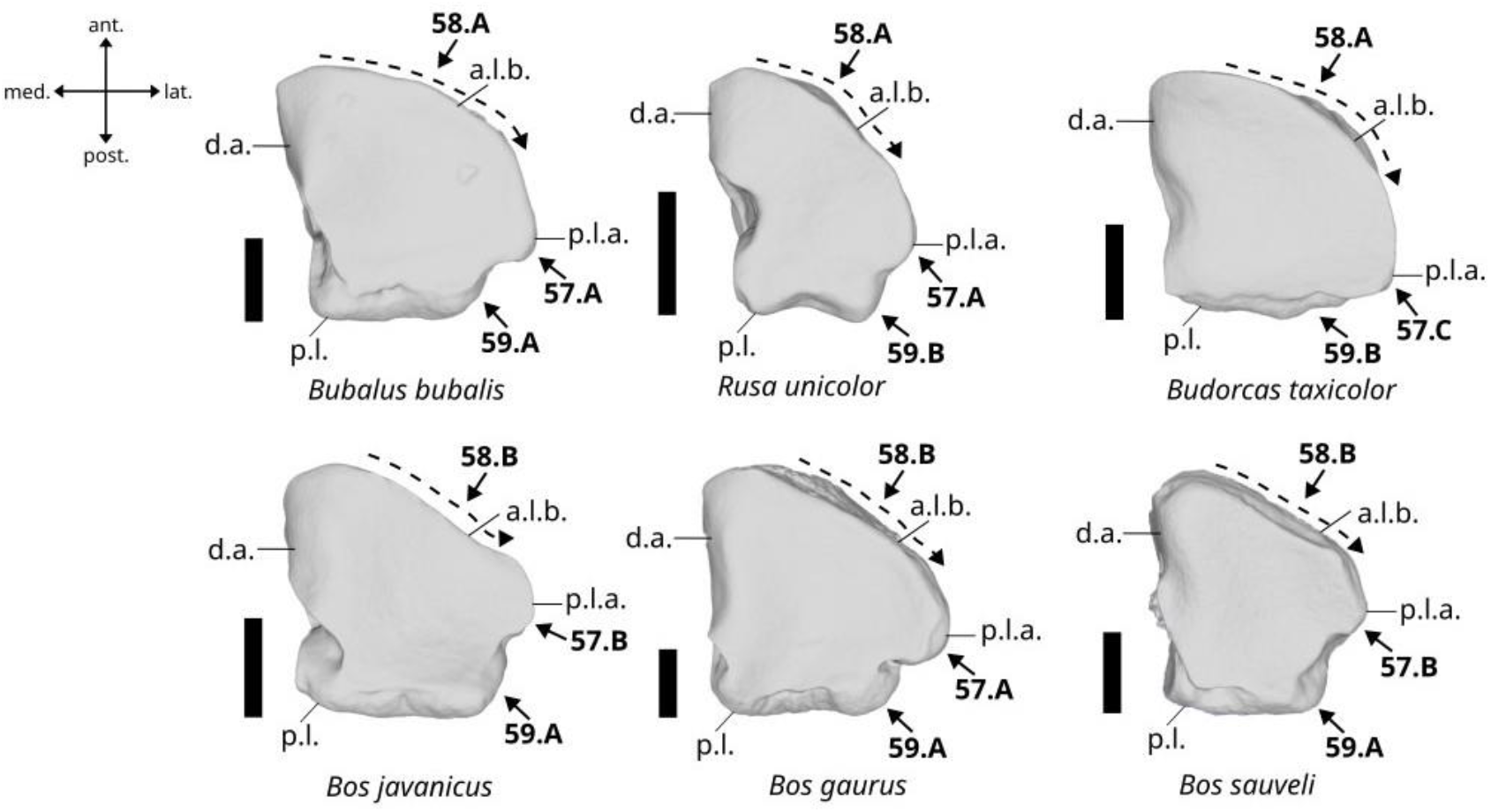
Character states visible on the distal face of the left unciform (character 57, 58, and 59). Abbreviation: a.l.b.: anterolateral border, p.l.a.: posterolateral apex, p.l.: posterior limit, d.a.: distal articular surface. Scalebar = 1 cm.

A. It is more or less individualized and oriented in posterolateral direction.
B. It is more or less individualized and oriented in lateral direction.
C. It is not individualized.

##### 58. Morphology of the antero-lateral border of the bone in distal view (Fig. 37)

A. It is curved.
B. It is straight.

##### 59. Expansion of the distal articular surface in distal view. (Fig. 37)

A. The articular surface nearly reaches the posterior limit of the bone.
B. The articular surface reaches the posterior limit of the bone.

##### 60. Morphology of the distal area of the medial articular facet in medial view. (Fig. 38)

**Figure 38.**
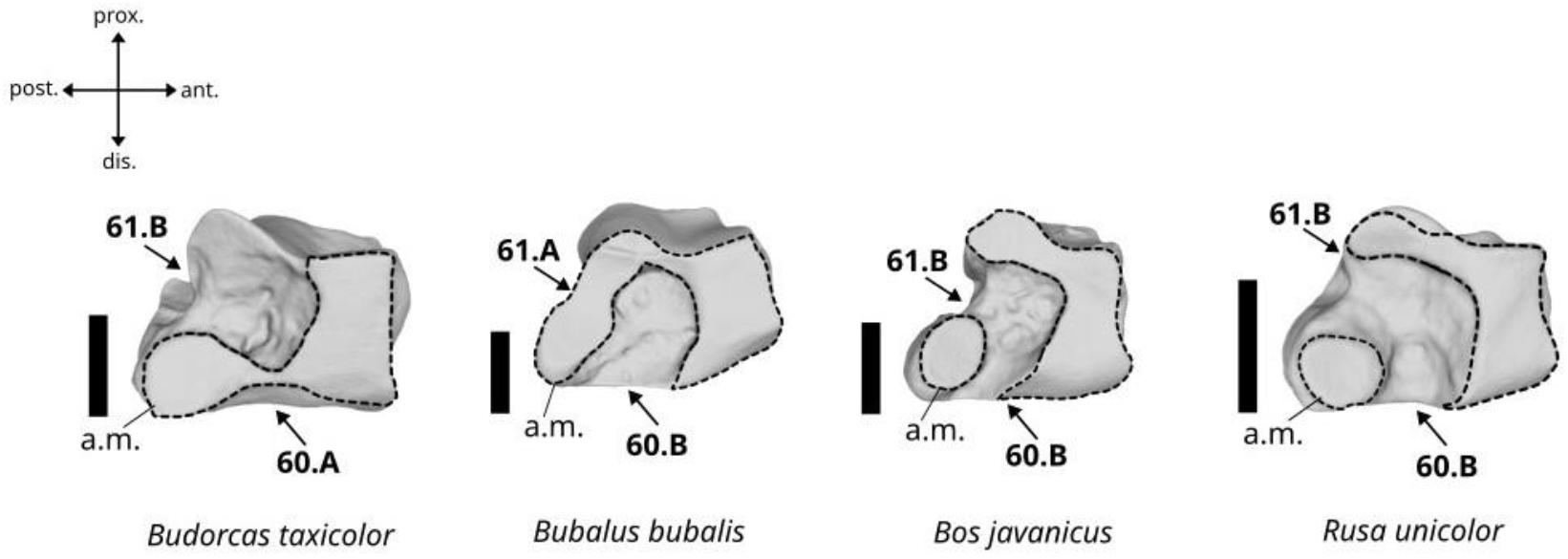
Character states visible on the medial face of the left unciform (characters 60 and 61). Abbreviation: a.m.: articular surface for the magnum bone (medial). Scalebar = 1 cm.

A. Continuous all along the distal edge of the bone.
B. Anterior and posterior articular areas separated by a gap.

##### 61. Morphology of the posterodistal area of the medial articular surface in medial view. (Fig. 38)

A. In contact with the proximal part of the articular facet.
B. Separated from the proximal part of the articular facet.

#### Characters of the metacarpus

##### 62. Morphology of the metacarpus in anterior view. (Fig. 39)

**Figure 39.**
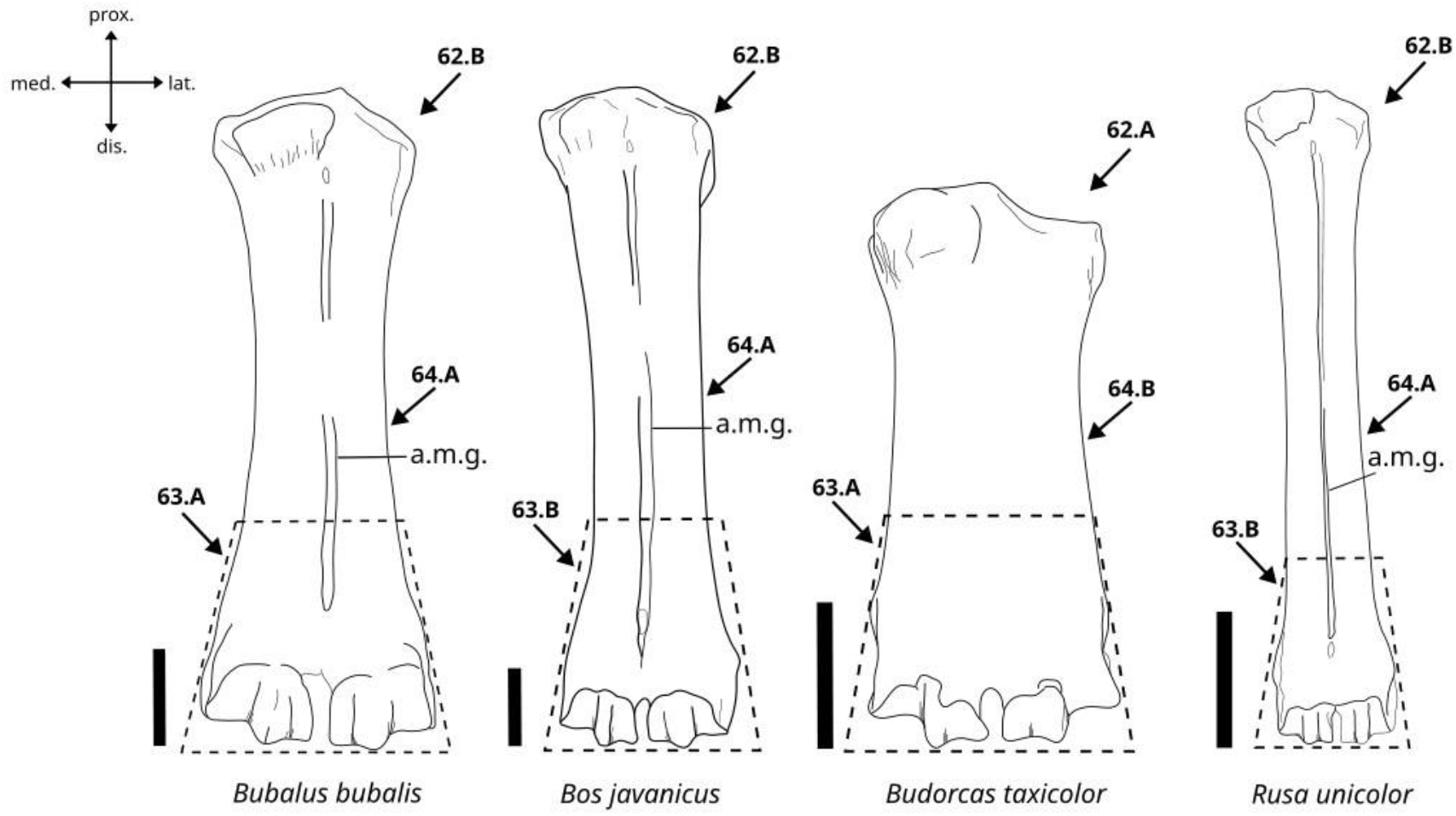
Character states visible on the anterior face of the left metacarpus (characters 62, 63, and 64). Abbreviation: a.m.g.: anteromedian groove. Scalebar = 3 cm.

A. It is very short and wide.
B. It is elongate and slender.

##### 63. Morphology of the distal part of the metacarpus in anterior view. (Fig. 39)

A. It is broad.
B. It is slender.

##### 64. Morphology of the anteromedian groove in anterior view. (Fig. 39)

A. Present.
B. Absent or very weakly marked.

##### 65. Shape of the articular surface for the magnum bone in proximal view. (Fig. 40)

**Figure 40.**
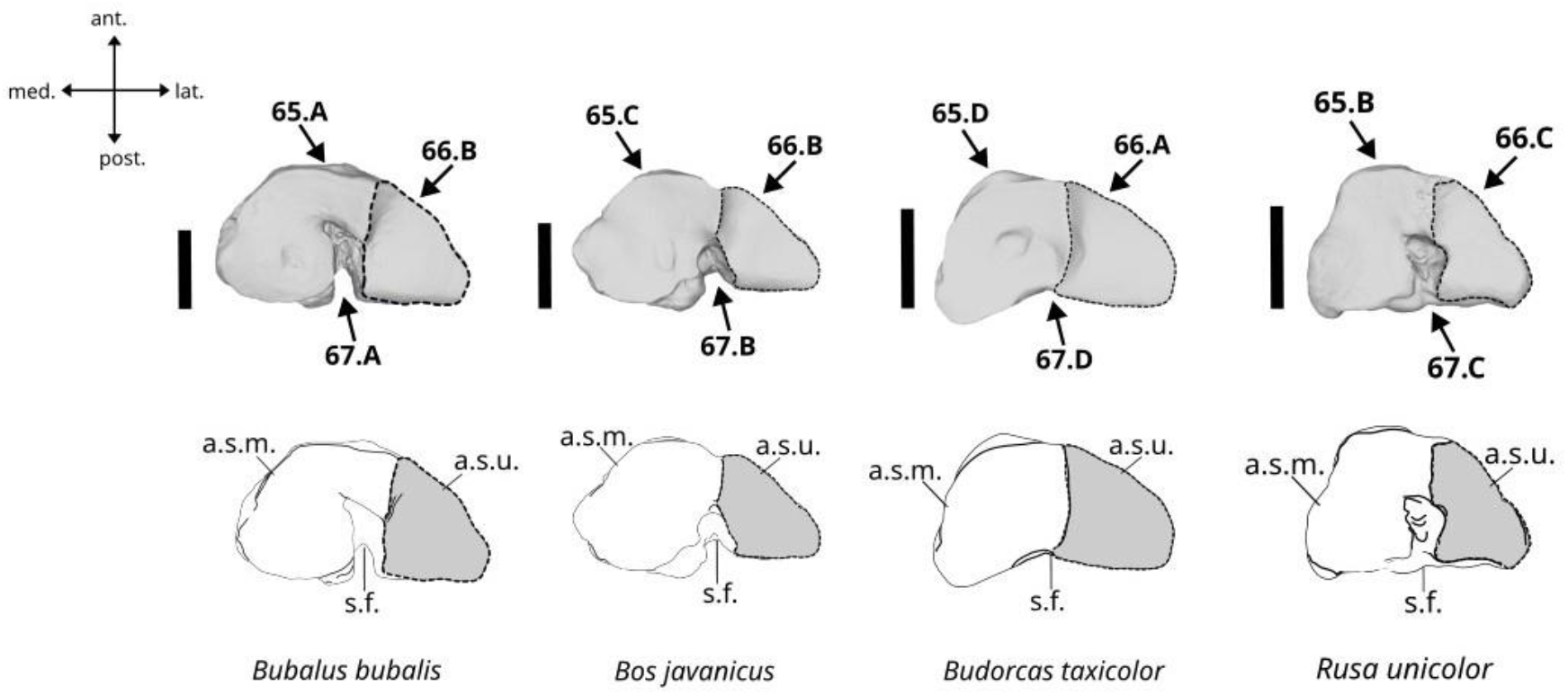
Character states visible on the proximal face of the left metacarpus (characters 65, 66, and 67). Abbreviations: a.s.m.: articular surface for the magnum bone, a.s.u.: articular surface for the unciform bone, s.f.: synovial fossa. Scalebar = 2 cm.

A. The articular surface is roughly as long as wide, and its lateral border is curved.
B. The articular surface is triangular, and its lateral border is sinuous.
C. The articular surface is roughly as long as wide, and the lateral border is sinuous.
D. The articular surface is ovoid and elongated, and the lateral border is curved.

##### 66. Medial expansion of the articular surface for the unciform bone in proximal view. (Fig. 40)

A. Strongly expanded.
B. Moderately expanded.
C. Weakly expanded.

##### 67. Morphology of the synovial fossa and the posterior tuberosity in proximal view. (Fig. 40)

A. The synovial fossa is deep, and its distal wall is open.
B. The synovial fossa is shallow, and its distal wall is open.
C. The synovial fossa is deep, and its distal margin is closed.
D. There is no synovial fossa.

##### 68. Morphology of the medial tuberosity in anterior view. (Fig. 41)

**Figure 41.**
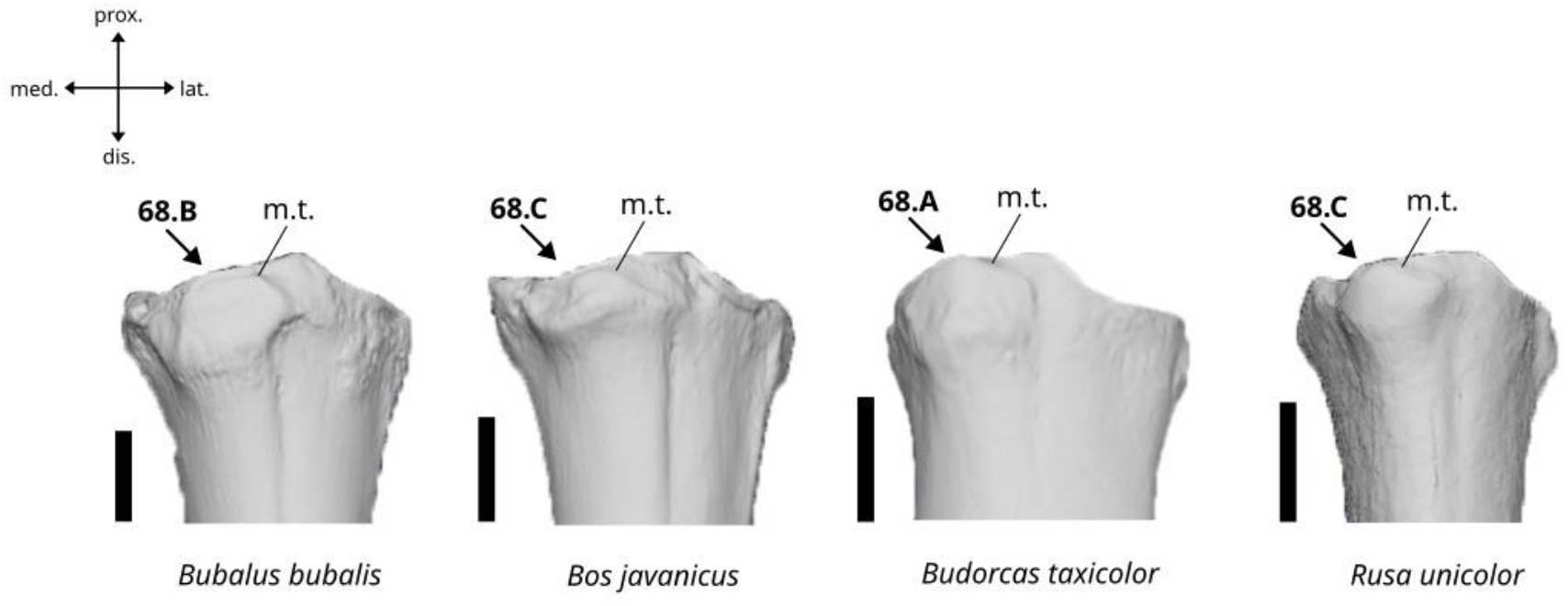
Character states visible on the anterior face of the left proximal metacarpus (character 68). Abbreviation: m.t.: medial tuberosity. Scalebar = 2 cm.

A. Strongly developed and expanded beyond the proximal limit of the proximal articular surface.
B. Strongly developed but not expanded beyond the proximal end of the proximal articular surface.
C. Moderately developed and not expanded beyond the proximal end of the proximal articular surface.

#### Characters of the coxal bone

##### 69. Occurrence of a notch in the antero-ventral part of the semi-lunar area of the acetabulum. (Fig. 42)

**Figure 42.**
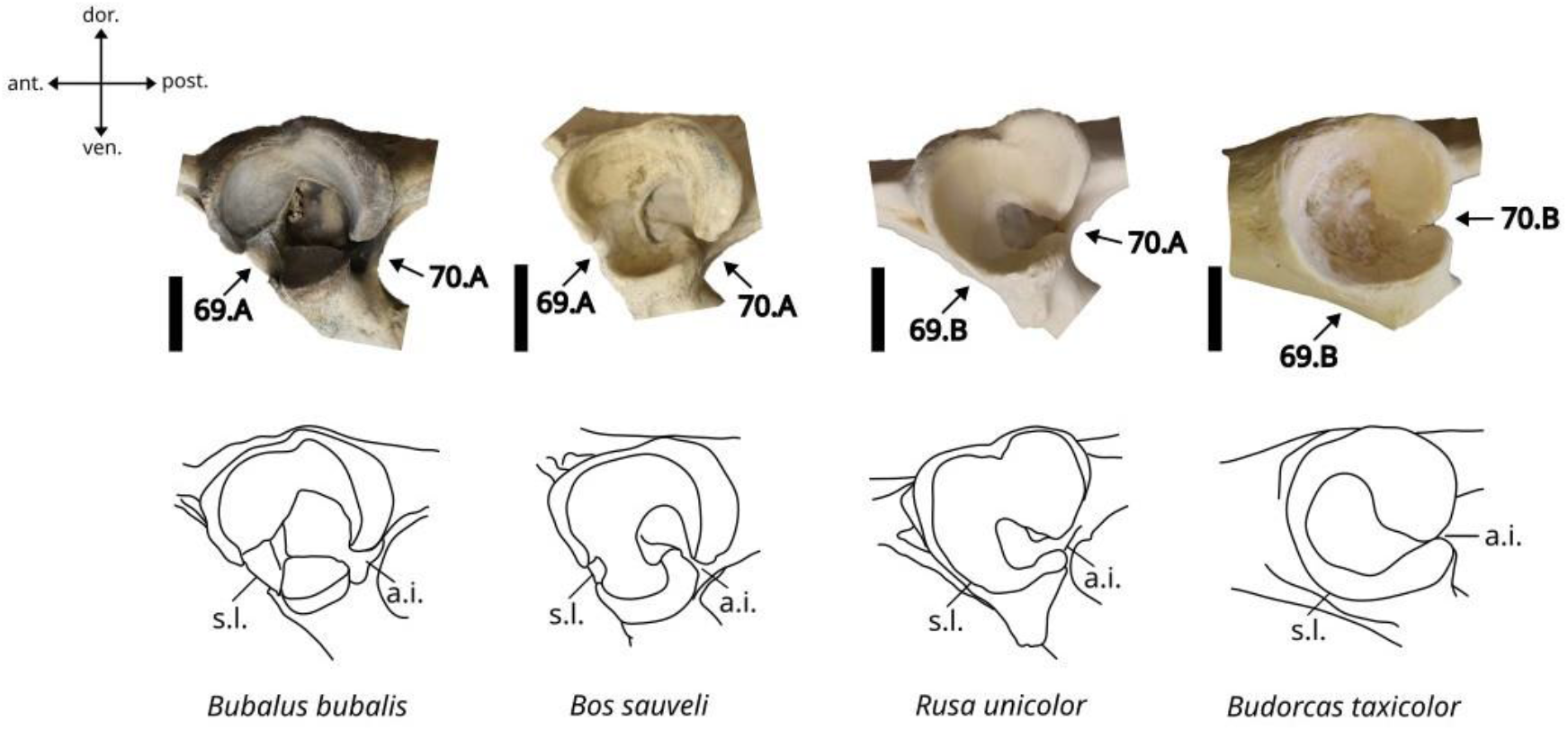
Character states visible on the lateral face of the left acetabulum (characters 69 and 70). Abbreviations: a.i.: acetabulum incisor, s.l.: semi-lunar area of the acetabulum. Scalebar =3 cm.

A. Yes.
B. No.

##### 70. Morphology of the acetabulum incisor. (Fig. 42)

A. Narrow but clearly open.
B. Almost closed.

##### 71. Morphology of the dorsal border of the acetabulum in dorsal view. (Fig. 43)

**Figure 43.**
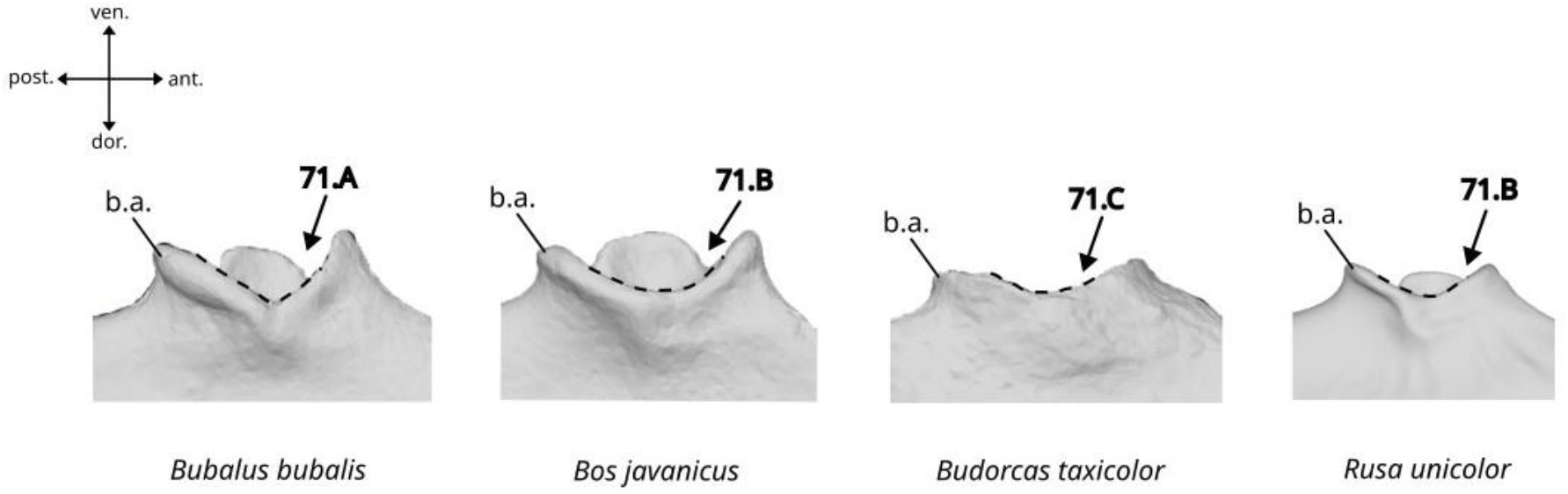
Character states visible on the dorsal face of the left acetabulum (character 71). Abbreviations: b.a.: lateral border of the acetabulum.

A. Forming a deep angular medial depression.
B. Forming a deep and curved medial depression.
C. Forming a shallow medial depression.

##### 72. Morphology of the tuberosity of the iliac spine. (Fig. 44)

**Figure 44.**
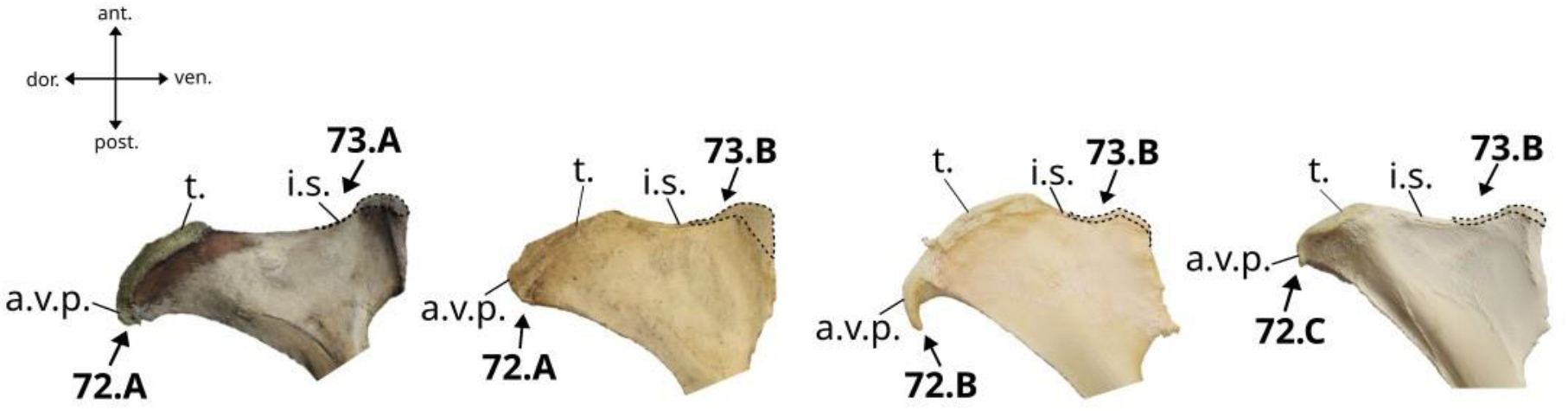
Character states visible on the lateral face of the left iliac spine of the coxal bone (characters 72 and 73). Abbreviations: t.: tuberosity, i.s.: iliac spine.

A. It is well-extended on the anterior margin of the bone and does not present an anteroventral process.
B. It is well-extended on the anterior margin of the bone and presents an antero-ventral process incurved in posteroventral direction.
C. It is moderately extended on the anterior margin of the bone and does not present an anteroventral process.

##### 73. Length of the iliac spine in lateral view. (Fig. 44)

A. Elongated
B. Shortened

##### 74. Shape of the ischial tuberosity in posterior view. (Fig. 45)

**Figure 45.**
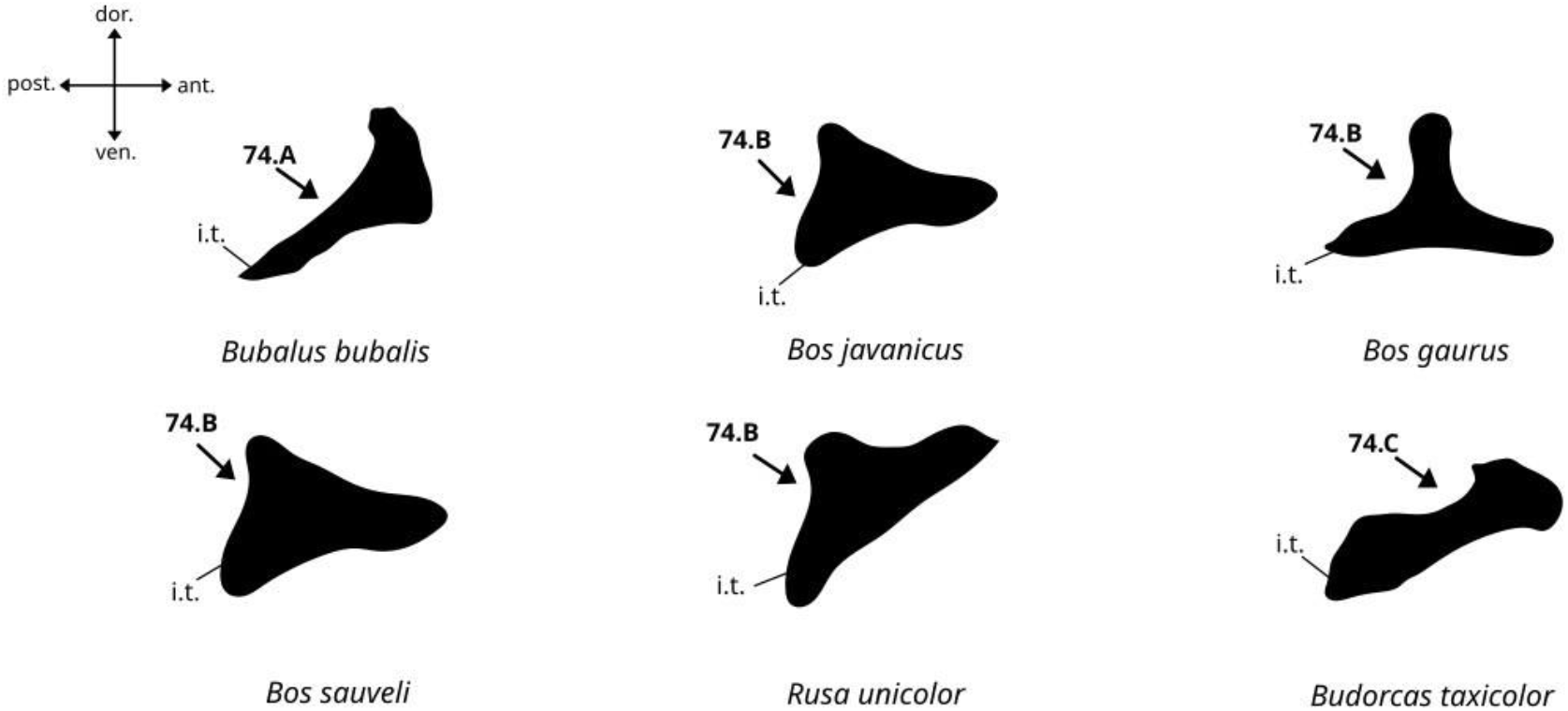
Character states visible on the posterior face of the left coxal bone (character 74). Abbreviation: i.t.: ischial tuberosity.

A. Slender and more or less triangular.
B. Short, wide, and more or less triangular.
C. Elongated, wide, and more or less quadrangular.

##### 75. Morphology of the ischial tuberosity in anterolateral view. (Fig. 46)

**Figure 46.**
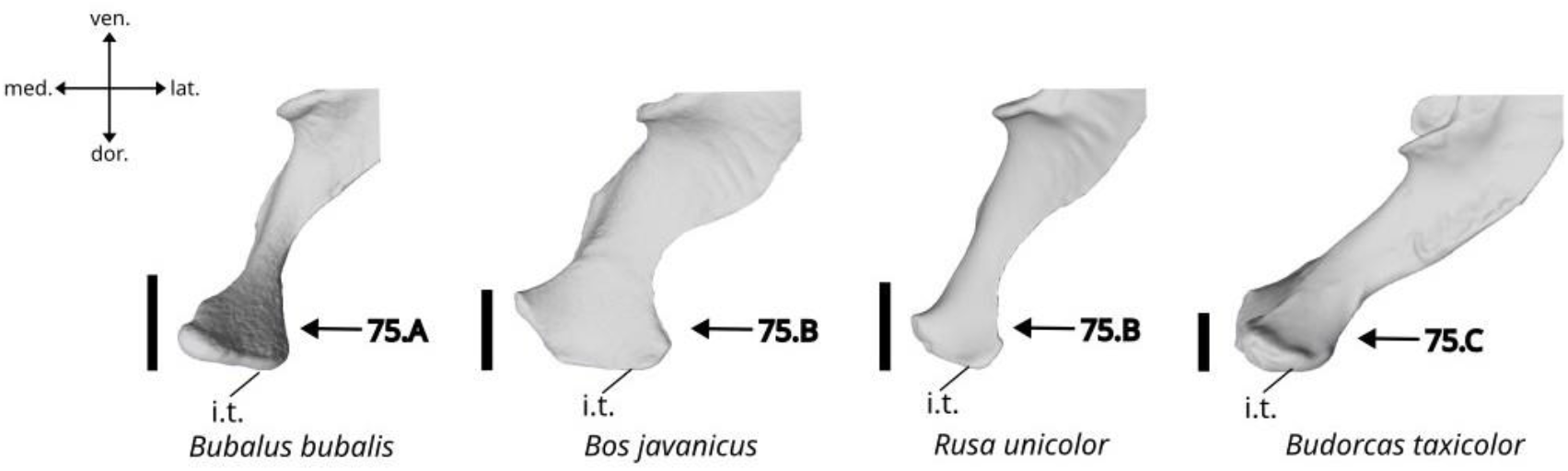
Character states visible on the anterolateral face of the left coxal bone (character 75). Abbreviation: ischial tuberosity. Scalebar = 5 cm.

A. Short and broad, and well-individualized.
B. Long and thin, and well-individualized.
C. Poorly individualized.

##### 76. Length of the iliac neck. (Fig. 47)

**Figure 47.**
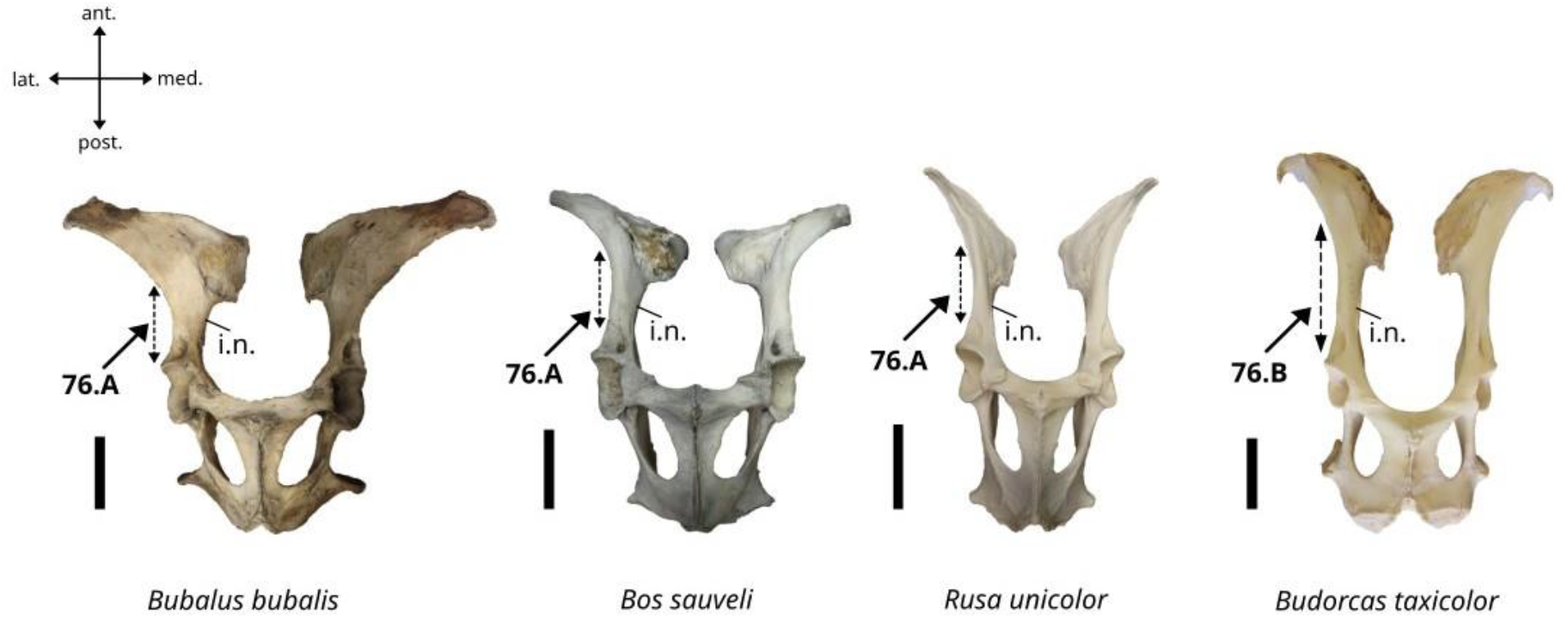
Character states visible on the ventral face of the coxal bone (character 76). Abbreviation: i.n.: iliac neck. Scalebar = 10 cm.

A. Short.
B. Long.

#### Characters of the femur

##### 77. Morphology of the intertrochanteric crest. (Fig. 48)

**Figure 48.**
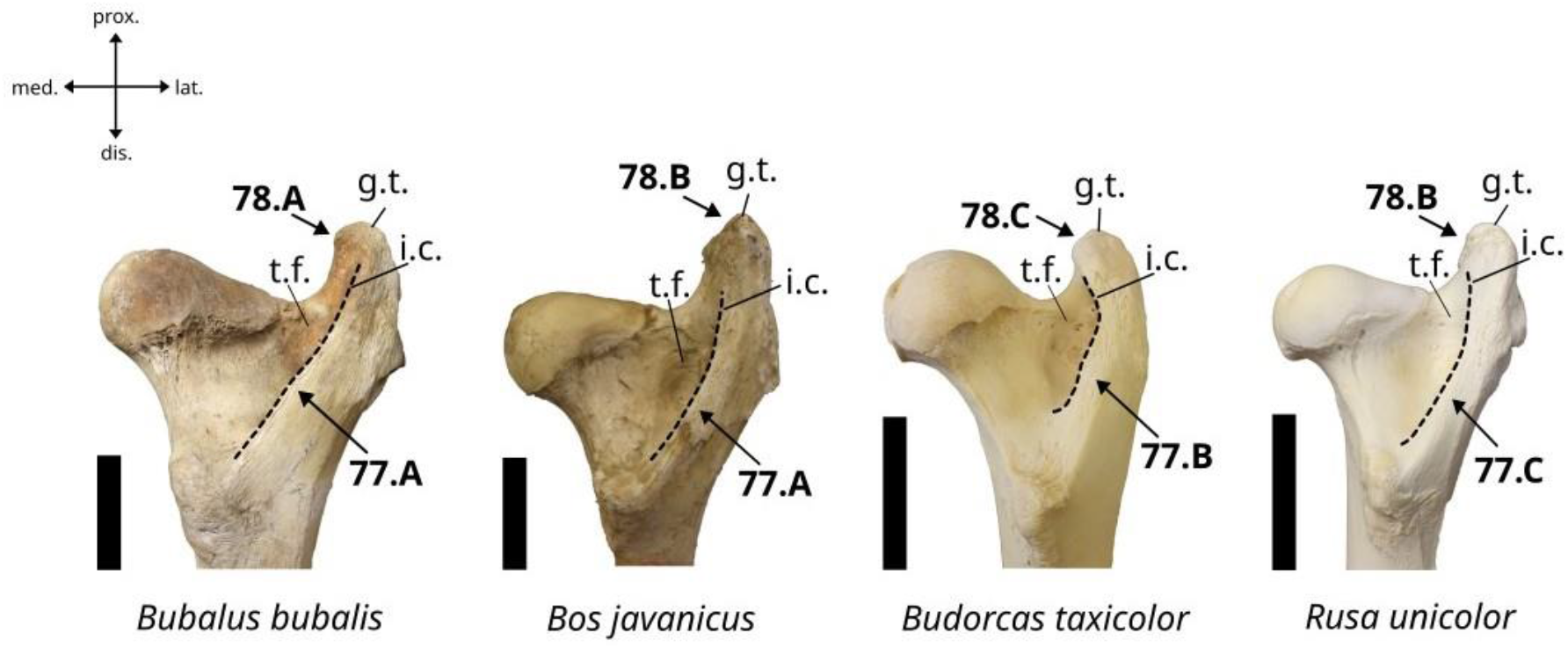
Character states visible on the posterior face of the right femur (characters 77 and 78). Abbreviations: g.t.: greater trochanter, i.c.: intertrochanteric crest, t.f.: trochanteric fossa. Scalebar = 5 cm.

A. The intertrochanteric crest is well expanded over the trochanteric fossa in posterior view. The margin of this crest is thick and blunt.
B. The intertrochanteric crest is not expanded over the trochanteric fossa in posterior view. The margin of the crest is thick and blunt.
C. The intertrochanteric crest does not expand over the trochanteric fossa in posterior view. The margin of this crest is thin and sharp.

##### 78. Morphology of the top of the greater trochanter in posterior view. (Fig. 48)

A. The top of the greater trochanter is slightly higher than the top of the femur head and, the trochanter is straight.
B. The top of the greater trochanter is clearly higher than the top of the femur head, and the trochanter is straight.
C. The top of the greater trochanter is slightly higher than the top of the femur head, and the trochanter is slightly inclined in medial direction.

##### 79. Delimitation of the attachment area of the gluteus medium muscle. (Fig. 49)

**Figure 49.**
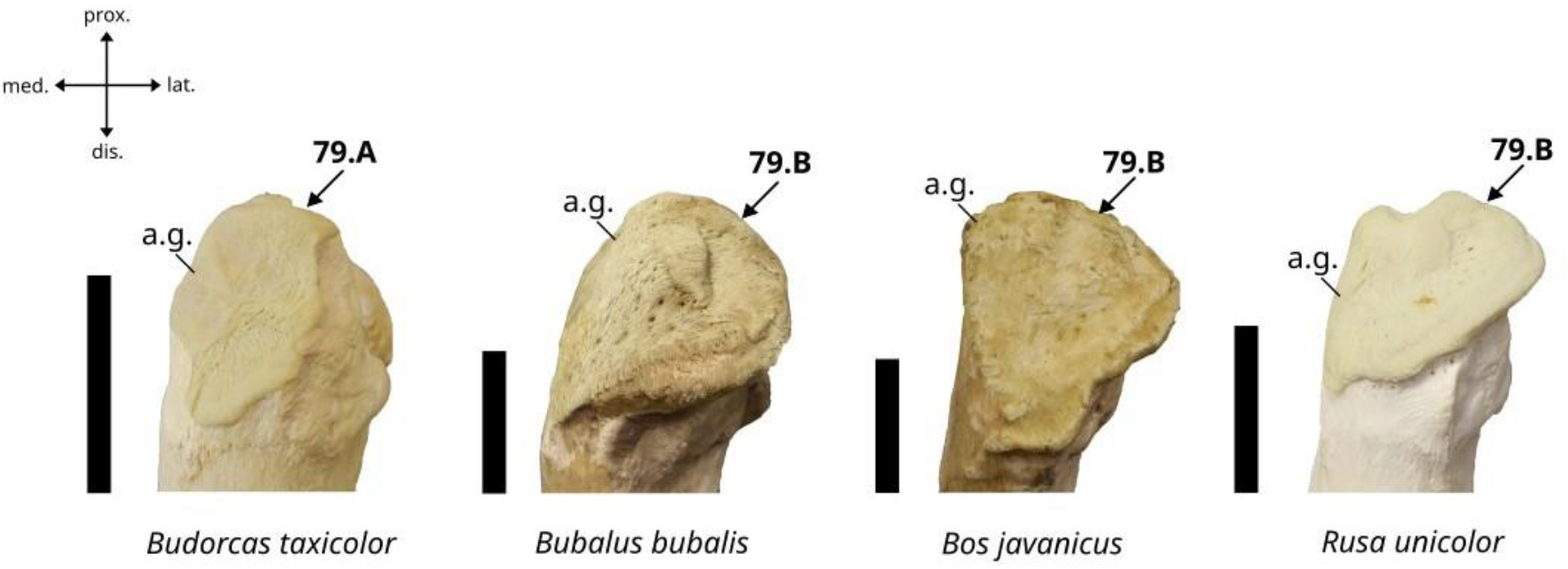
Character states visible on the lateral face of the right femur (character 79). Abbreviation: a.g.: attachment area of the gluteus medium muscle. Scalebar = 5 cm.

A. Poorly defined with an irregular outline.
B. Well defined with a regular outline.

##### 80. Morphology of the trochlea in anterior view. (Fig. 50)

**Figure 50.**
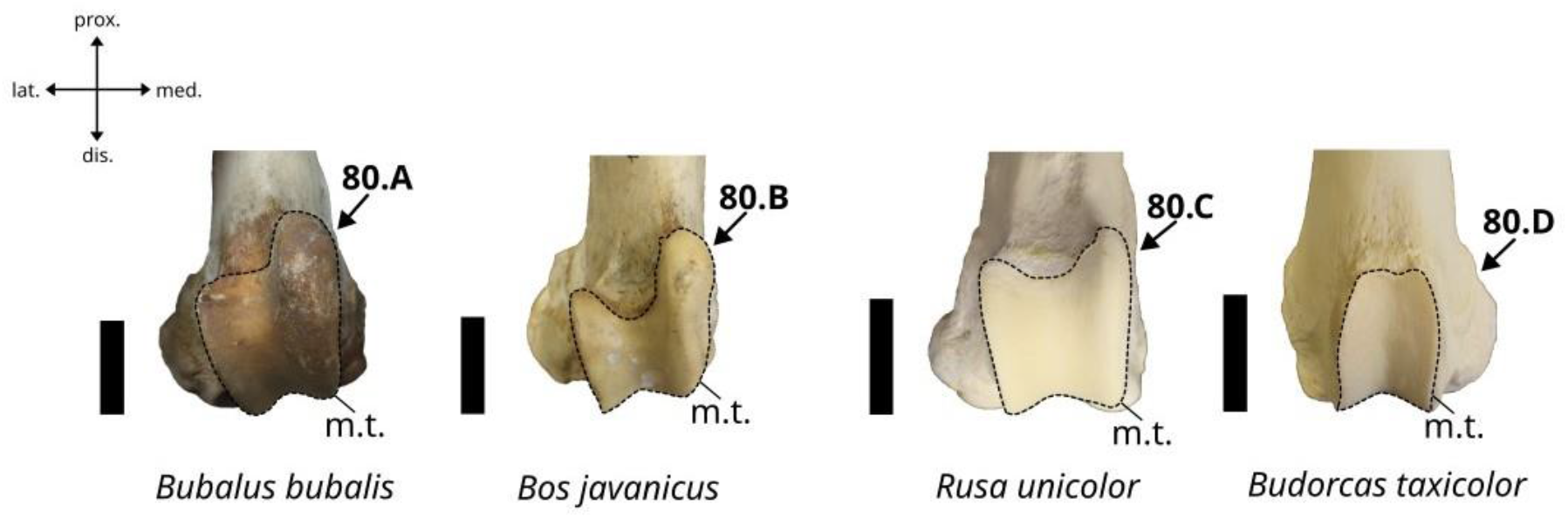
Character states visible on the anterior face of the right femur (character 80). Abbreviation: m.t.: medial trochlea. Scalebar = 5 cm.

A. The lateral and medial tubercles are moderately individualized, with the medial one being considerably higher than the lateral one.
B. The lateral and medial tubercles are strongly individualized, separated by a well-marked notch, with the medial one being considerably higher than the lateral one.
C. The lateral and medial tubercles are moderately individualized, with the medial one being slightly higher than the lateral one.
D. The trochlea is thin and the lateral and medial tubercles are not individualized.

#### Characters of the patella

##### 81. Occurrence of a groove between the base of the anterior tuberosity and the medial angle of the bone in anterior view. (Fig. 51)

**Figure 51.**
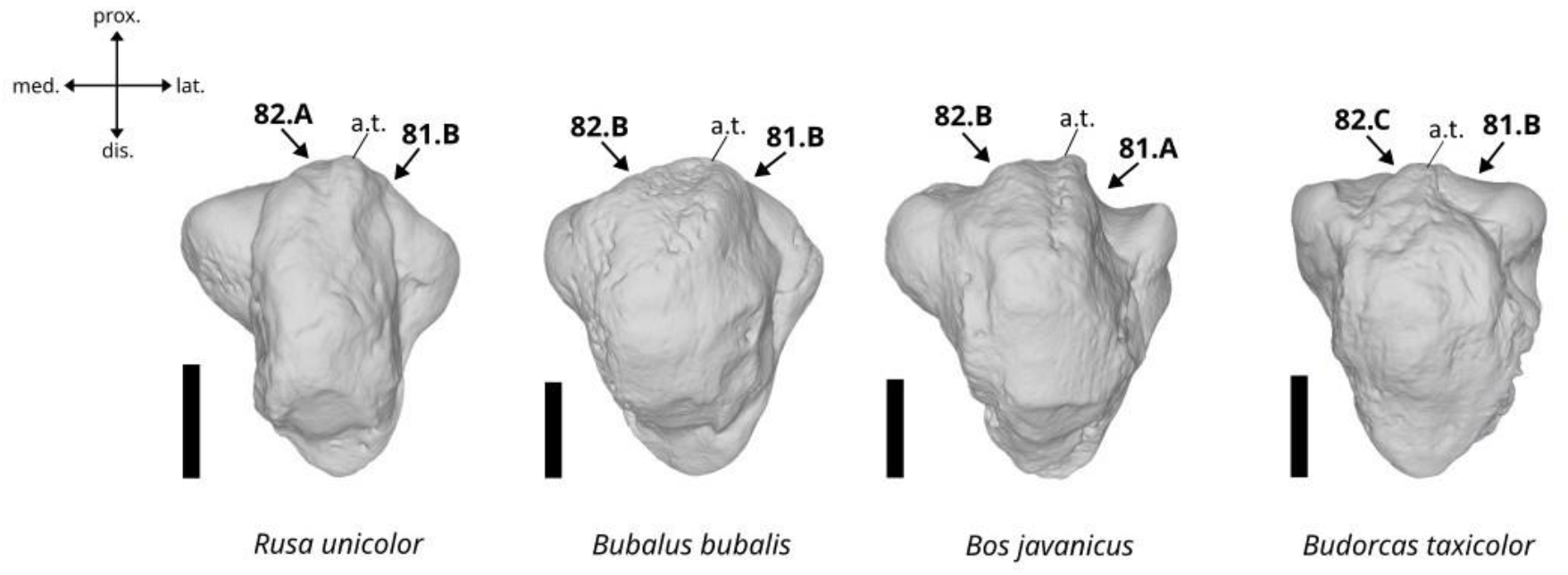
Character states visible on the anterior face of the left patella (characters 81 and 82). Abbreviations: a.t.: anterior tuberosity. Scalebar = 2 cm.

A. Present.
B. Absent.

##### 82. Expansion of the anterior tuberosity in anterior view. (Fig. 51)

A. The tuberosity is strongly expanded beyond the proximal border of the bone.
B. The tuberosity is weakly expanded beyond the proximal border of the bone.
C. The tuberosity is not expanded beyond the proximal border of the bone.

##### 83. Morphology of the medial angle in posterior view. (Fig. 52)

**Figure 52.**
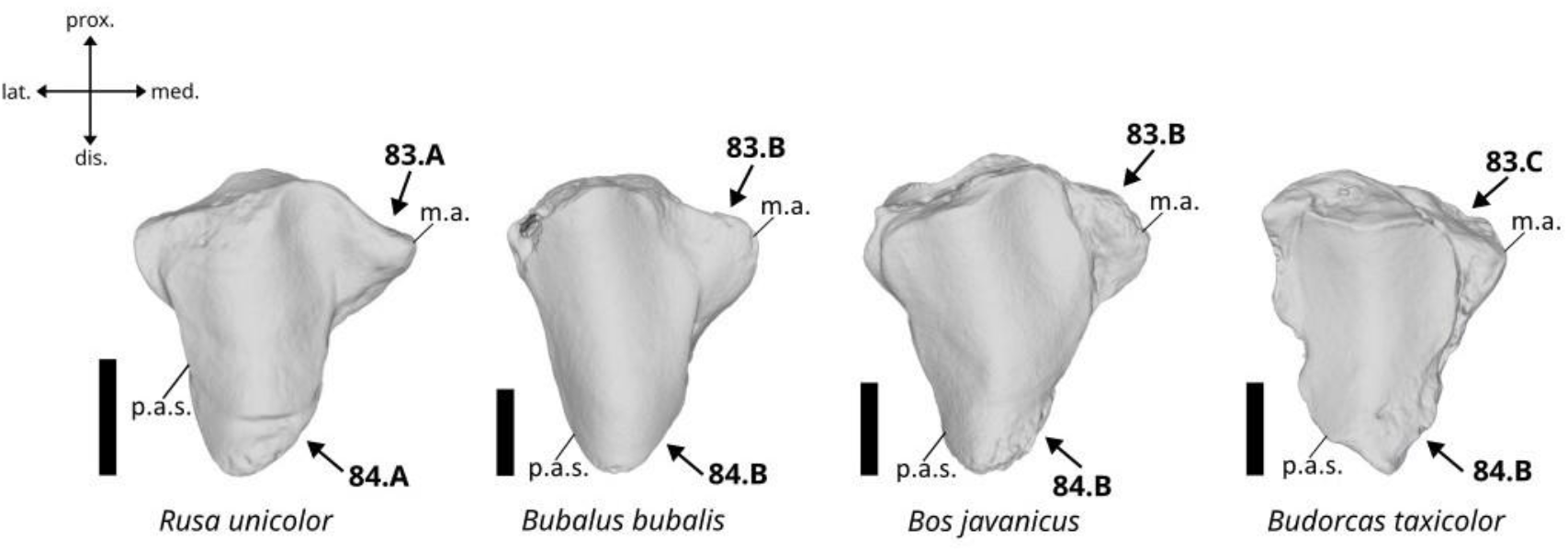
Character states visible on the posterior face of the left patella (characters 83 and 84). Abbreviations: m.a.: medial angle, p.a.s.: posterior articular surface. Scalebar = 2 cm.

A. It is well-developed in medial direction, triangular with a pointed apex, and is slightly inclined in posterior direction.
B. It is well-developed in medial direction, of truncated triangular shape, and not inclined in posterior direction.
C. It is poorly developed.

##### 84. Morphology of the distal limit of the posterior articular surface in posterior view. (Fig. 52)

A. It is located at a high position far away from the distal limit of the bone.
B. It reaches the distal limit of the bone.

#### Characters of the tibia

##### 85. Morphology of the latero-proximal area of the tibial crest in anterior view. (Fig. 53)

**Figure 53.**
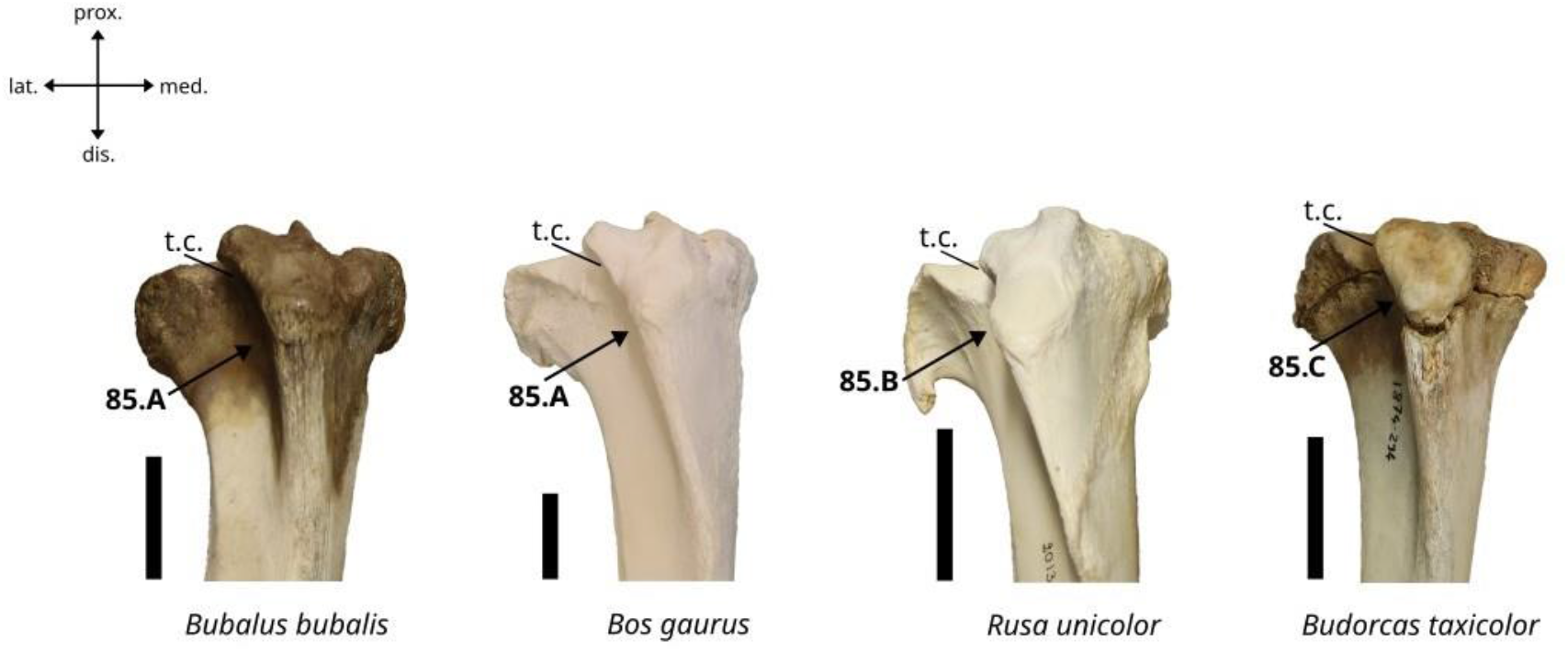
Character states visible on the anterior face of the right tibia (character 85). Abbreviation: t.c.: tibial crest. Scalebar = 5 cm.

A. It is well-developed, inclined in lateral direction, and its lateral border is more or less straight.
B. It is well-developed, slightly inclined in lateral direction, with a straight lateral border presenting a shallow but very well-marked notch in its dorsal area.
C. It is reduced, very slightly inclined in lateral direction, and its lateral border is straight.

##### 86. Morphology of the posterolateral extremity of the proximal articular surface in proximal view. (Fig. 54)

**Figure 54.**
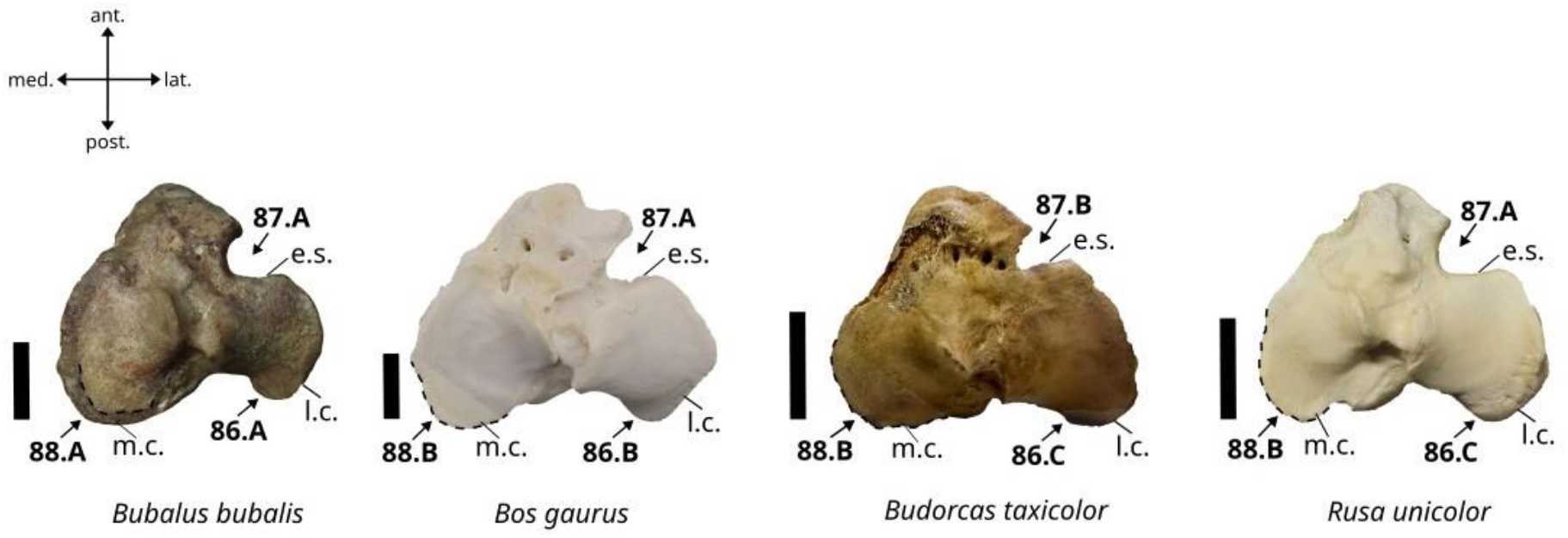
Character states visible on the proximal face of the right tibia (characters 86, 87, and 88). Abbreviations: l.c.: lateral condyle, m.a.: medial condyle, e.s.: extensor sulcus. Scalebar = 3 cm.

A. It is well delimited, and its posterior margin is of truncated triangular outline.
B. It is moderately delimited, and its posterior margin is curved.
C. It is poorly delimited.

##### 87. Shape of the extensor sulcus in proximal view. (Fig. 54)

A. Wide and deep.
B. Narrow and short.

##### 88. Shape of the medial condyle of the tibia in proximal view. (Fig. 54)

A. The medial articular surface of the tibia does not reach the medial and posterior margins of the bone.
B. The medial articular surface of the tibia reaches the medial and posterior margins of the bone.

##### 89. Occurrence of an articular surface for the malleolar bone in anterolateral position in addition to the one occurring in posterolateral position in distal view. (Fig. 55)

**Figure 55.**
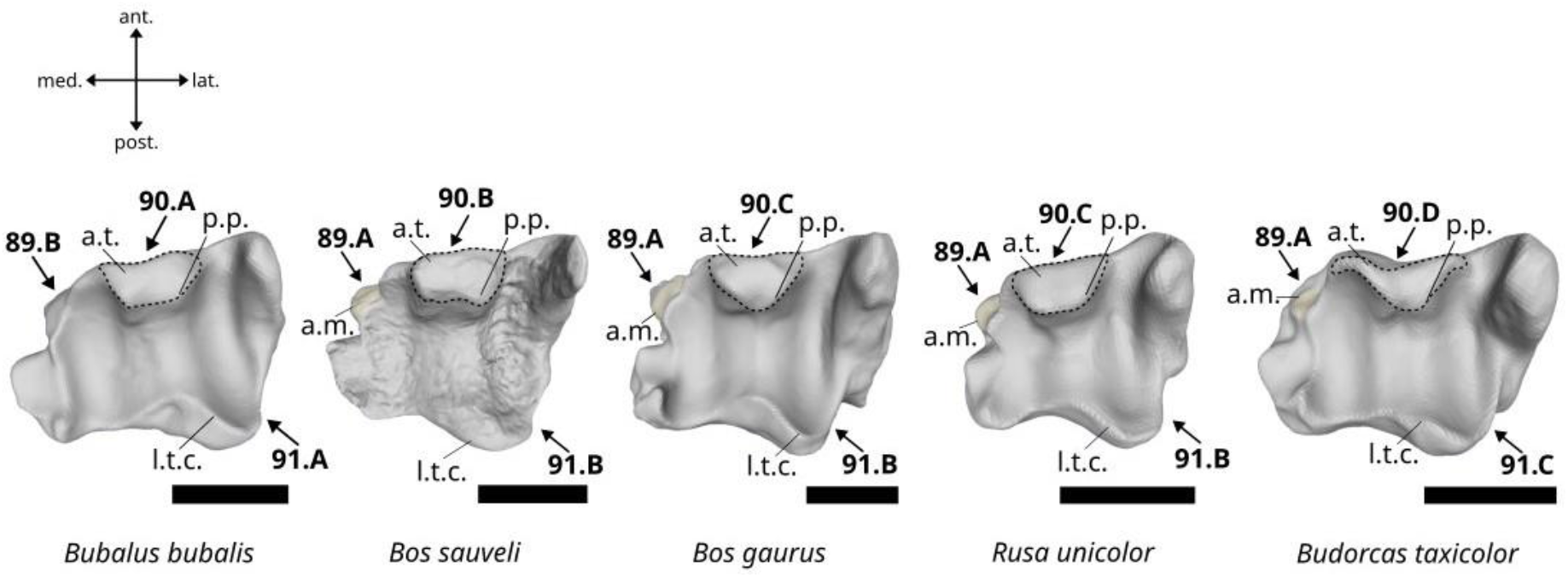
Character states visible on the distal face of the right tibia (characters 89, 90, and 91). Abbreviations: a.m.: anterior articular surface for the malleolar bone, a.t.: anterior tubercle, l.t.c.: lateral tibial cochlea, p.p.: posterior projection. Scalebar = 3 cm.

A. Present.
B. Absent.

##### 90. Shape of anterior tubercle in distal view. (Fig. 55)

A. It is squared with a single small posterior projection.
B. It is oblong with two small posterior projections.
C. It is semi-circular with a single small posterior projection.
D. It is crescent-shaped.

##### 91. Shape of the lateral tibial cochlea in distal view. (Fig. 55)

A. It is well-extended in posterolateral direction in comparison to the medial tibial cochlea.
B. It is moderately extended in posterolateral direction in comparison to the medial tibial cochlea.
C. The lateral tibial cochlea is nearly at the same level as the medial tibial cochlea.

##### 92. Morphology of the medial tuberosity in medial view. (Fig. 56)

**Figure 56.**
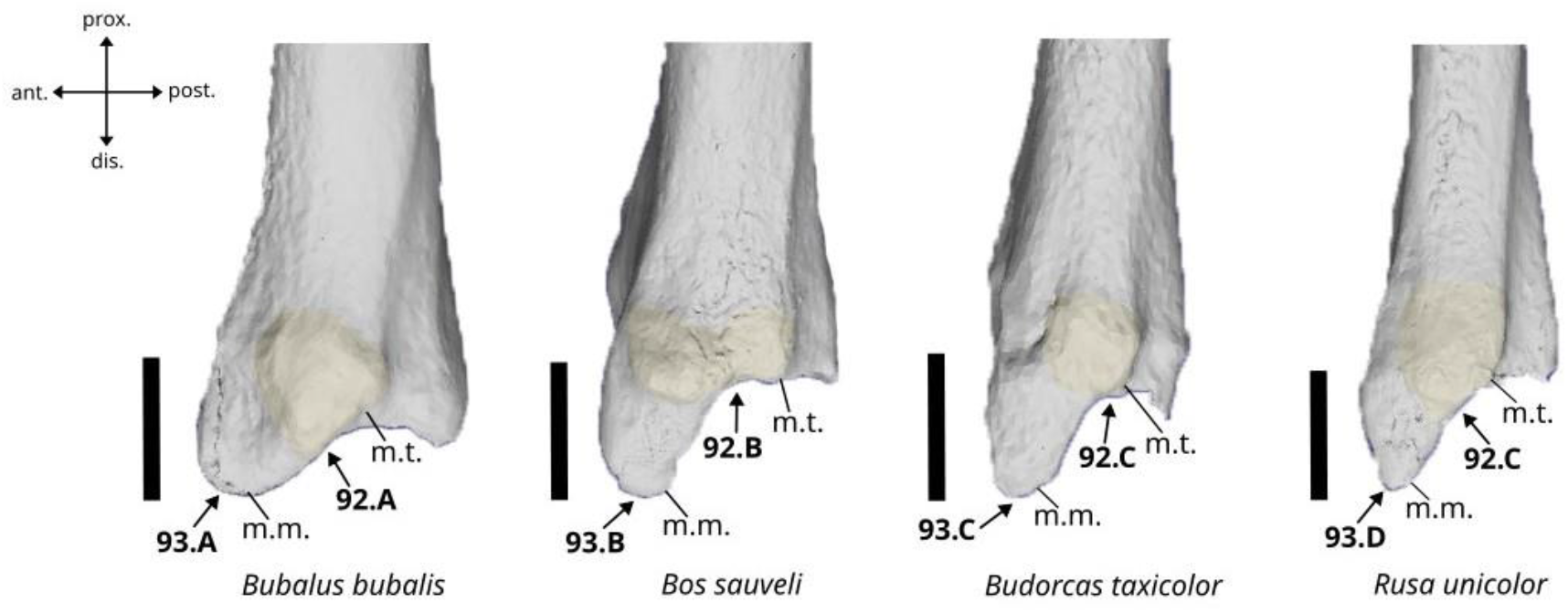
Character states visible on the lateral face of the right tibia (character 92 and 93). Abbreviations: m.m.: medial malleolar bone, m.t.: medial tubercle. Scalebar = 3 cm.

A. The medial tuberosity is moderately developed.
B. The medial tuberosity is well-developed.
C. The medial tuberosity is poorly developed.

##### 93. Morphology of the medial malleolar process in medial view. (Fig. 56)

A. It is projected in latero-distal direction with a rounded apex.
B. It is projected in distal direction with a blunt apex.
C. It is projected in distal direction with a truncated apex.
D. It is projected in distal direction with a pointed apex.

##### 94. Morphology of the postero-medial tubercle in posterior view. (Fig. 57)

**Figure 57.**
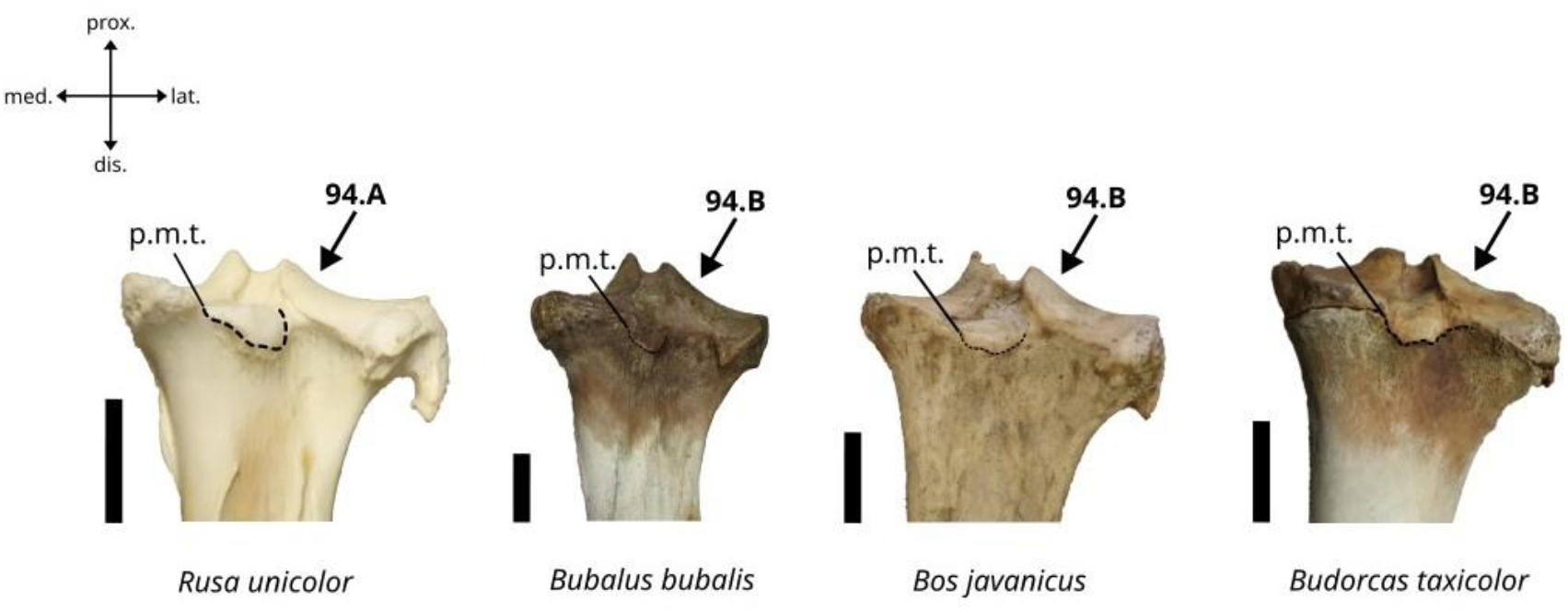
Character states visible on the posterior face of the right tibia (character 94). Abbreviation: p.m.t.: postero-medial tubercle. Scalebar = 3 cm.

A. Well expanded below the distal limit of the proximal epiphysis.
B. Moderately expanded below the distal limit of the proximal epiphysis.

#### Characters of the malleolar bone

##### 95. Morphology of the proximal spine in lateral view. (Fig. 58)

**Figure 58.**
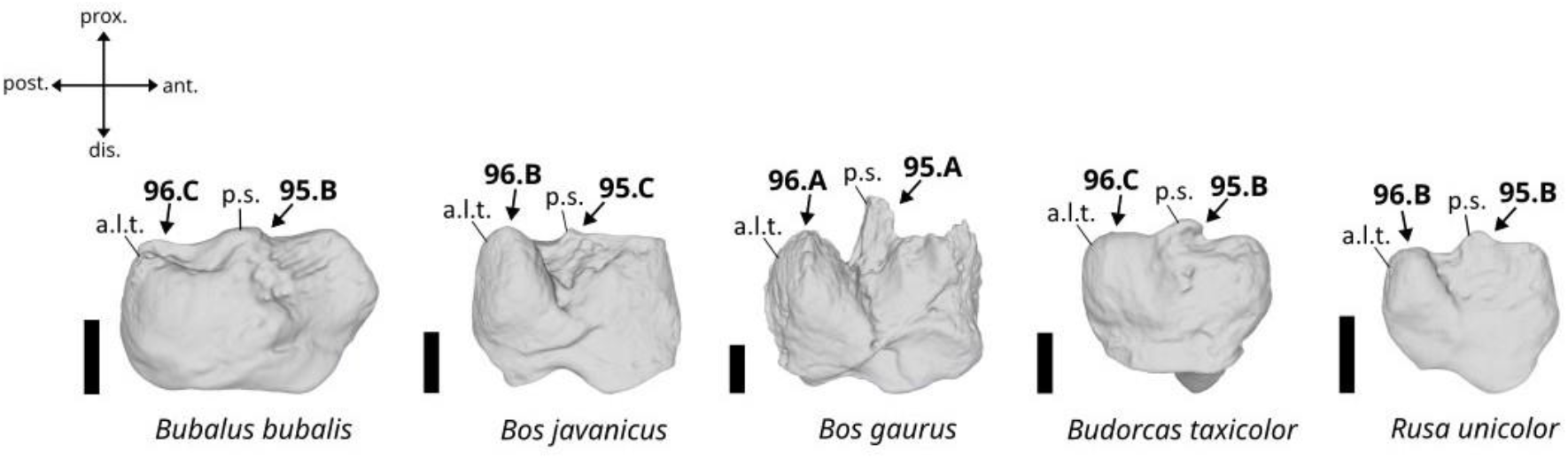
Character states visible on the lateral face of the right malleolar bone (characters 95 and 96). Abbreviations: p.s.: proximal spine, a.l.t.: anterolateral tuberosity. Scalebar = 1 cm.

A. It is well-developed.
B. It is moderately developed.
C. It is nearly absent.

##### 96. Morphology of the anterolateral tuberosity in lateral view. (Fig. 58)

A. Well-expanded beyond the proximal border of the bone.
B. Slightly expanded beyond the proximal border of the bone.
C. Not expanded beyond the proximal border of the bone.

##### 97. Morphology of the anterodistal area of the bone in medial view. (Fig. 59)

**Figure 59.**
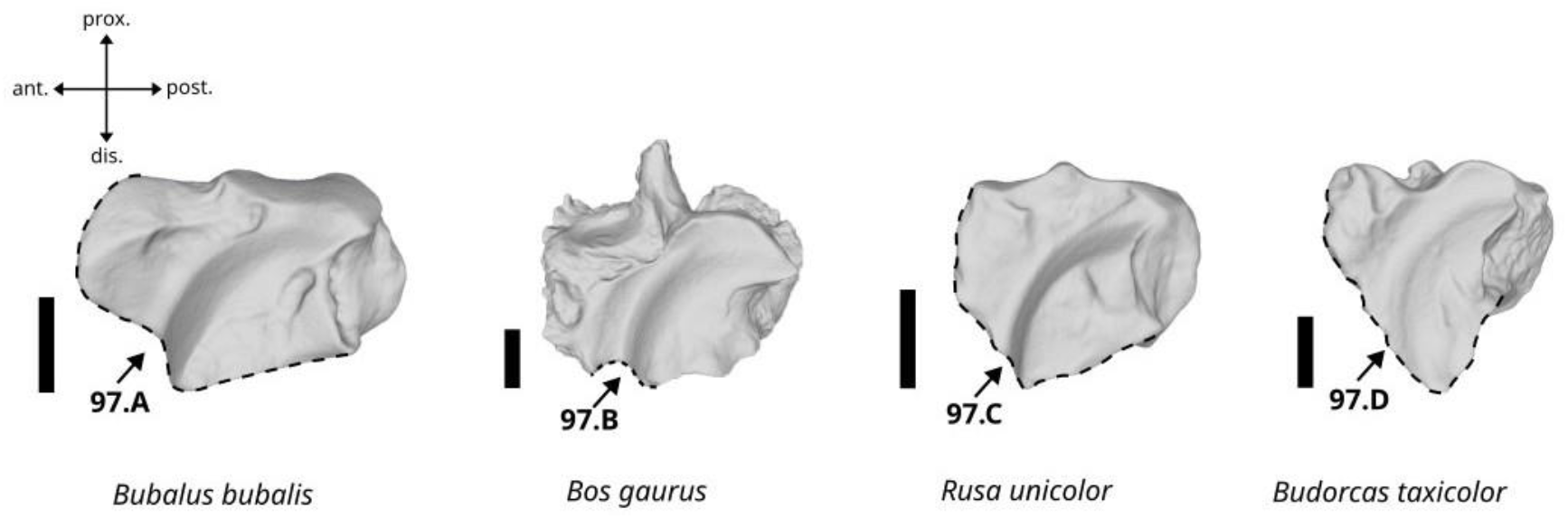
Character states visible on the medial face of the right malleolar bone (character 97). Scalebar = 1 cm.

B. It presents a small notch central notch.
A. It is sinusoidal.
C. It has a truncated triangular shape.
D. It is triangular with a blunted apex.

##### 98. Morphology of the distal articular surface in distal view. (Fig. 60)

**Figure 60.**
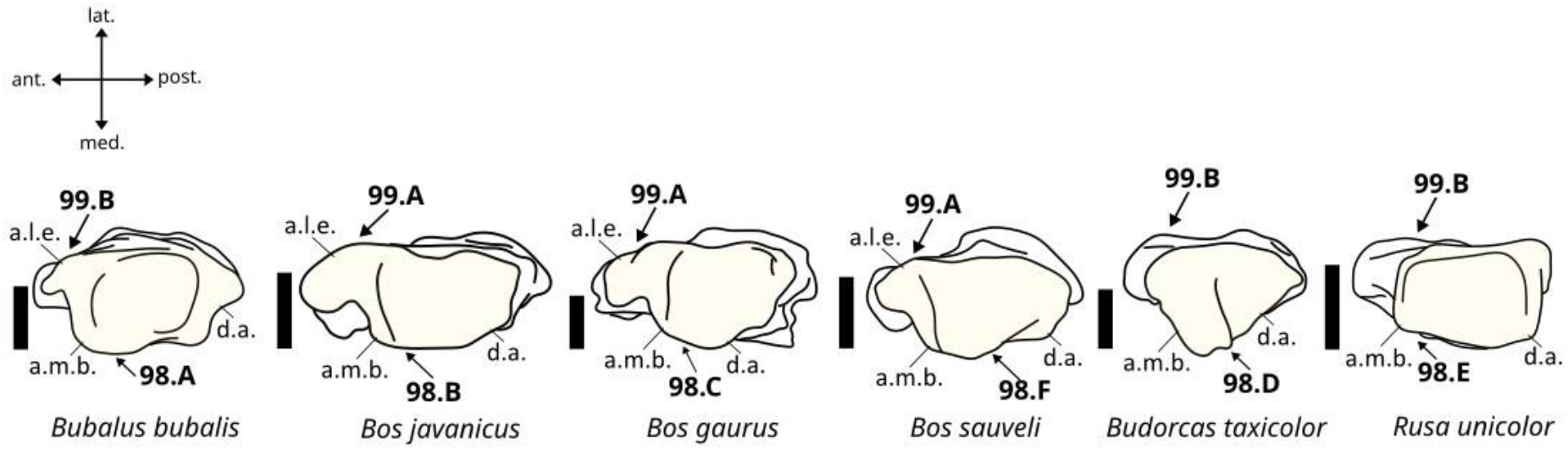
Character states visible on the distal face of the left malleolar bone (characters 98 and 99). Abbreviation: a.m.b.: anteromedial border, d.a.: distal articular surface, a.l.e.: antero-lateral expansion. Scalebar = 1 cm.

A. The articular surface is rectangular in shape, its anteromedial border is expanded in medial direction, and the medial margin is curved.
B. The articular surface is rectangular in shape, the anteromedial border is slightly expanded in medial direction, and the medial margin is straight.
C. The articular surface is trapezoid in shape, the anteromedial border is slightly expanded in medial direction, and the medial margin is curved.
D. The articular surface is triangular in shape, the anteromedial border is expanded in medial direction, and the medial margin is pointed.
E. The articular surface is rectangular and elongated, the anteromedial border does not expand in medial direction, and its medial margin is straight.
F. The articular surface is rectangular in shape, the anteromedial border is relatively expanded in medial direction, and the medial margin is straight.

##### 99. The bone presents a well-developed antero-lateral expansion of its distal articular surface in distal view. (Fig. 60)

A. Yes
B. No

#### Characters of the calcaneus

##### 100. Morphology of the groove on the calcaneal tuberosity in the posterior view. (Fig. 61)

**Figure 61.**
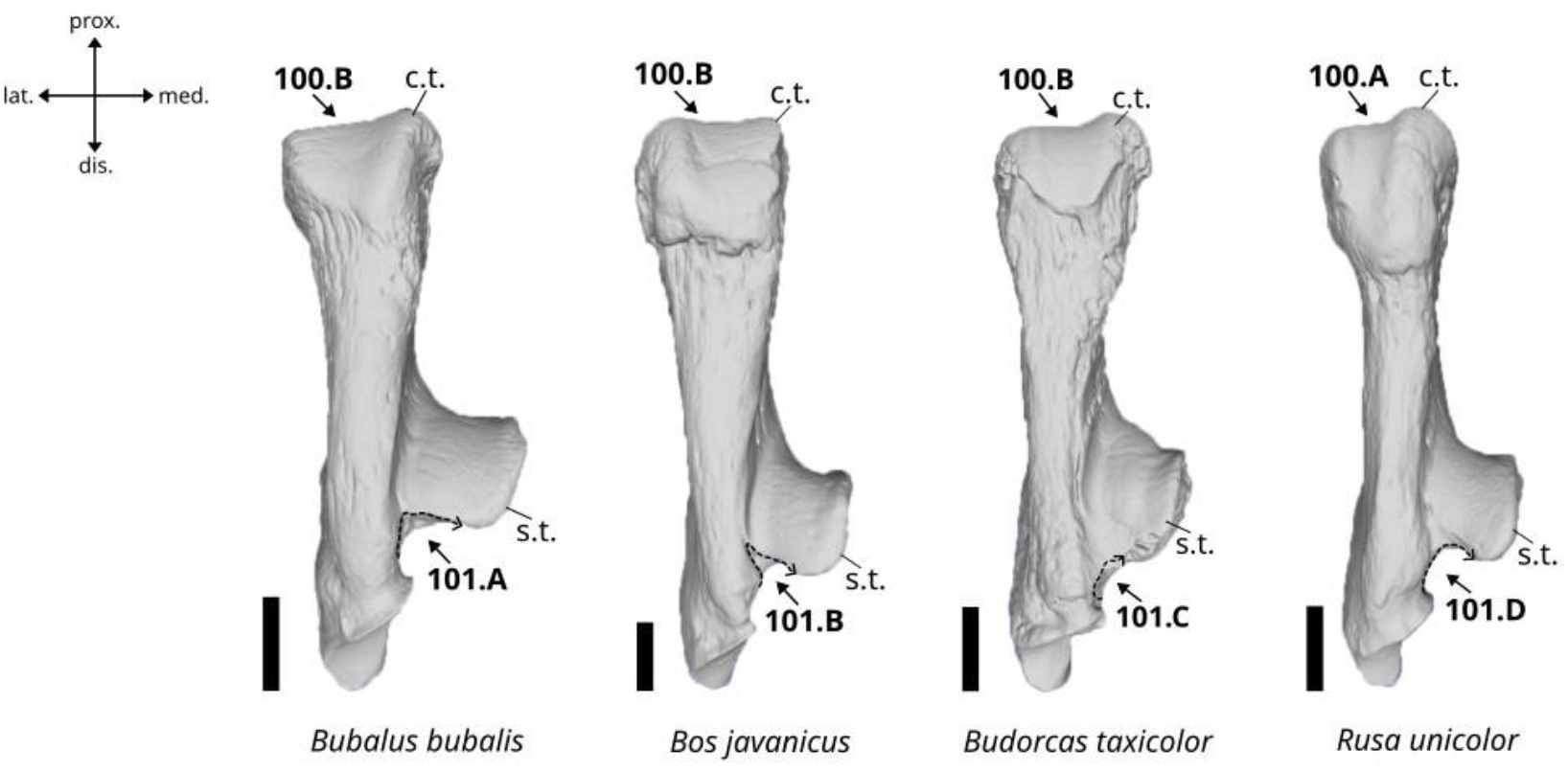
Character states visible on the posterior face of the left calcaneus (characters 100 and 101). Abbreviations: c.t.: calcaneal tuberosity, s.t.: sustentaculum tali. Scalebar = 2 cm.

A. Deep.
B. Shallow.

##### 101. Morphology of the posteromedial margin of the sustentaculum tali in posterior view. (Fig. 61)

A. It is deep and forms a right angle.
B. It is deep and forms an acute angle.
C. It is shallow with a curved lateral margin.
D. It is deep with a curved lateral margin.

##### 102. Occurrence of a notch in the dorsal area of the calcaneal tuberosity in medial view. (Fig. 62)

**Figure 62.**
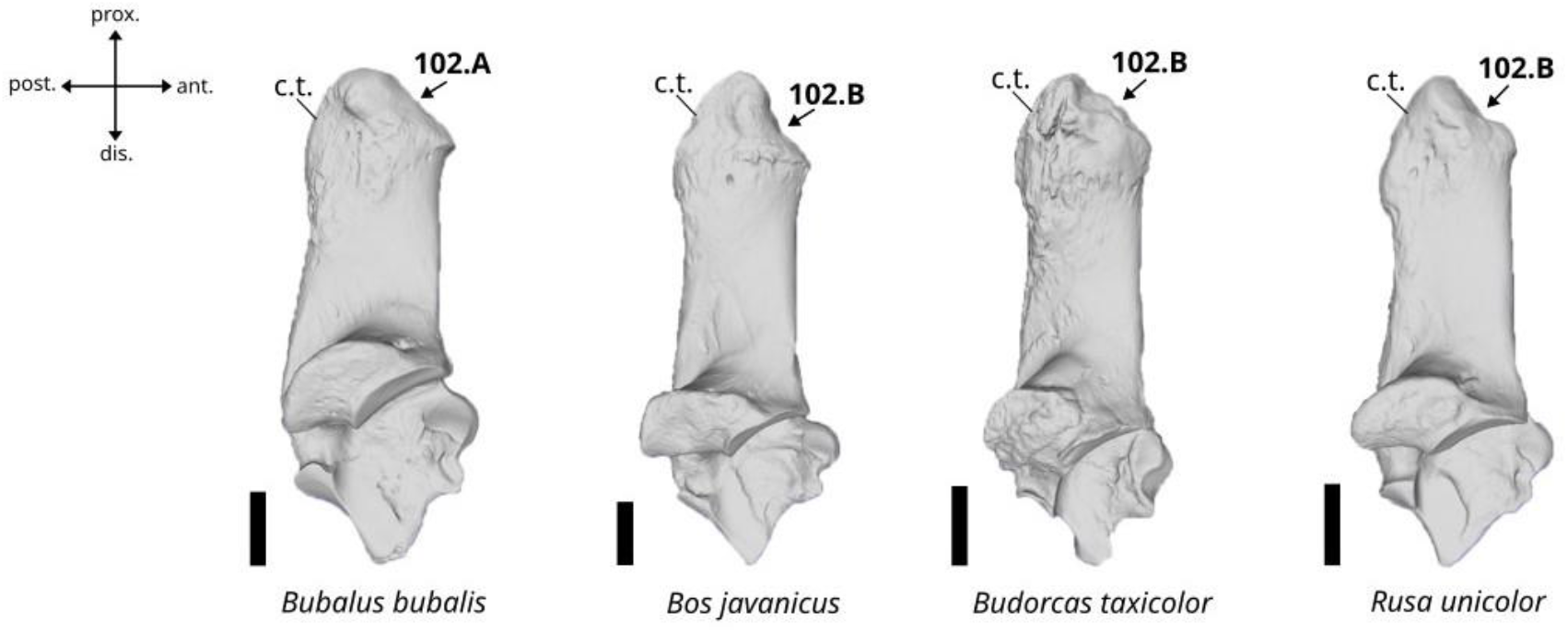
Character states visible on the medial face of the left calcaneus (character 102). Abbreviation: c.t.: calcaneal tuberosity. Scalebar = 2 cm.

A. Absent.
B. Present.

##### 103. Orientation of the articular surface for the malleolar bone in dorsal view. (Fig. 63)

**Figure 63.**
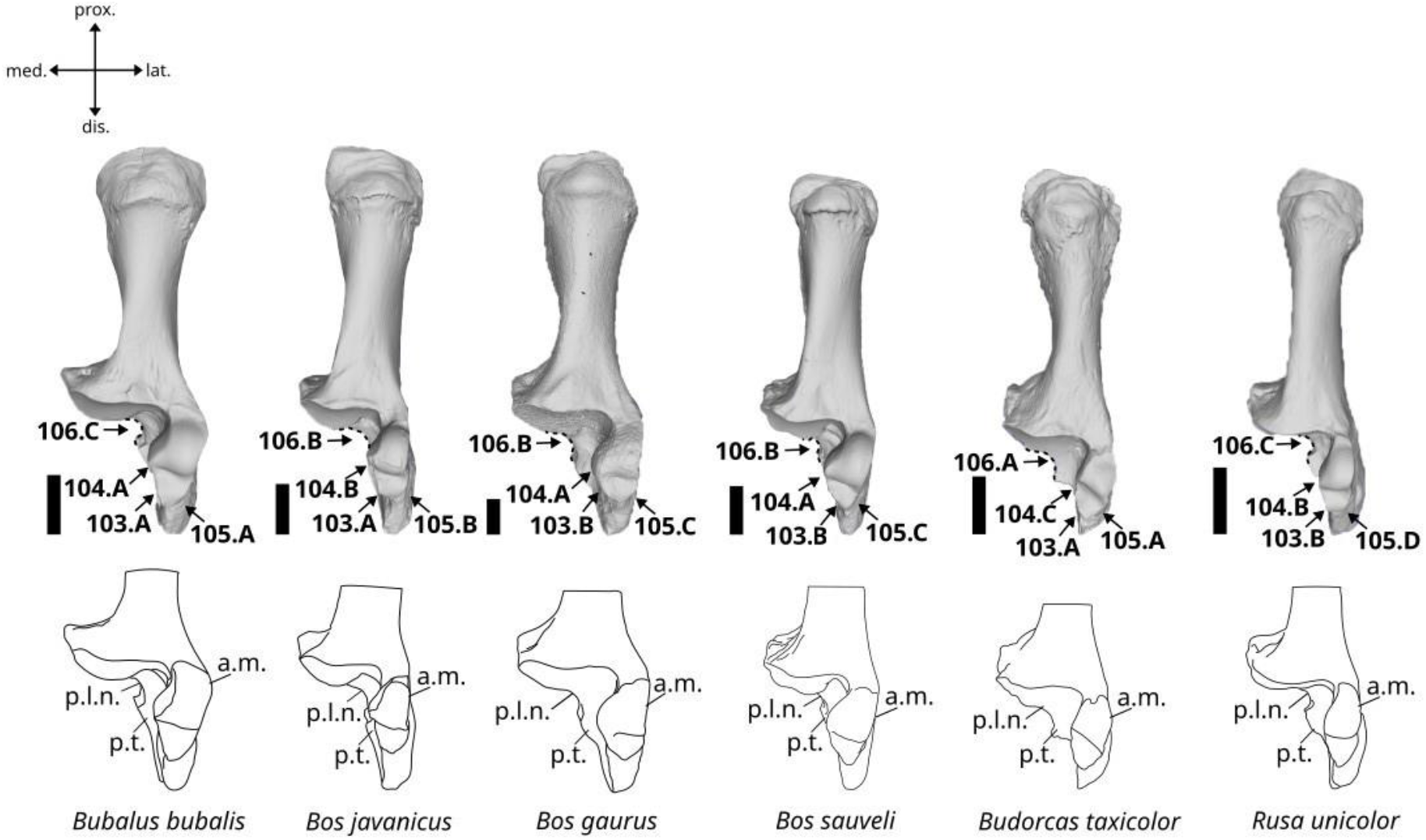
Character states visible on the dorsal face of the left calcaneus (characters 103, 104, 105 and 106). Abbreviations: a.m.: articular surface for the malleolar bone, p.l.n.: posterolateral notch, p.t.: plantar tubercle. Scalebar = 2 cm.

A. The articular surface is inclined in mediodistal direction.
B. The articular surface is straight.

##### 104. Medial extension of the articular surface for the malleolar bone in dorsal view. (Fig. 63)

A. Well-extended
B. Slightly extended with a curved medial margin
C. Slightly extended with a flat medial margin

##### 105. Morphology of the distal area of the articular surface for the malleolar bone in dorsal view. (Fig. 63)

A. Elongated with an acute apex
B. Short and broad with a truncated apex
C. Short and broad with an obtuse apex
D. Short and broad with an acute apex

##### 106. Morphology of the posteromedial wall of the plantar tubercle in anterior view. (Fig. 63)

A. Well-extended in medial direction with a shallow posterolateral notch.
B. Slightly extended in medial direction with a moderately deep posterolateral notch.
C. Slightly extended in medial direction with a deep and angled posterolateral notch.

#### Characters of the astragalus

##### 107. Outline of the laterodistal border in anterior view. (Fig. 64)

**Figure 64.**
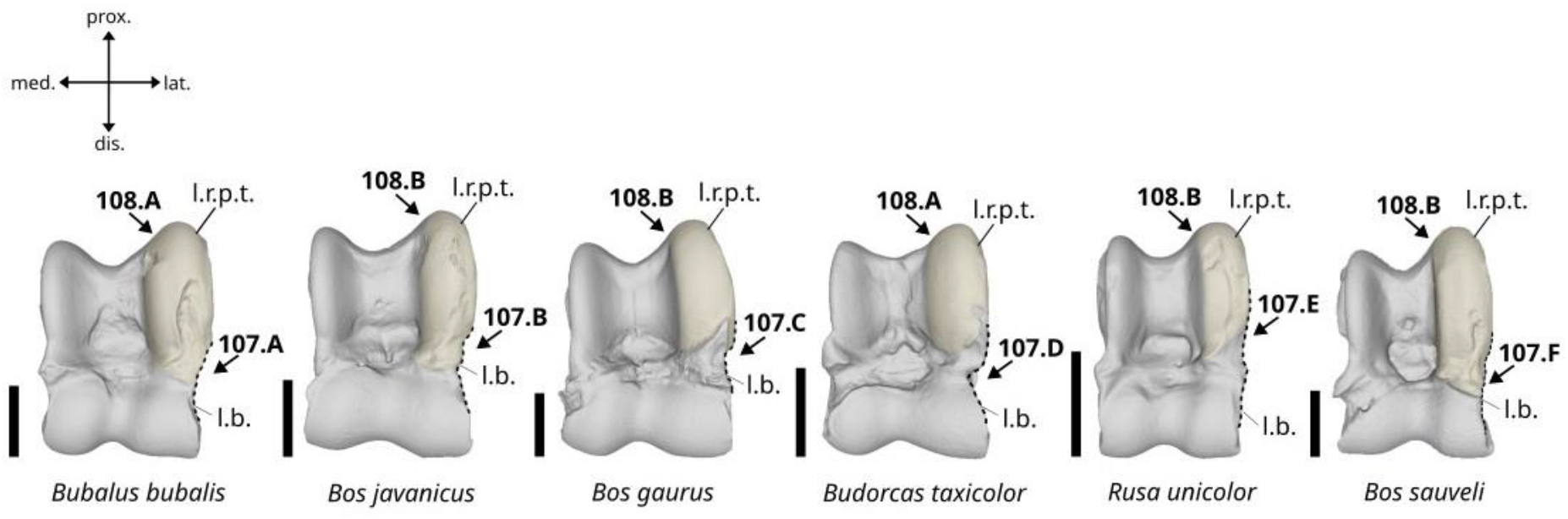
Character states visible on the anterior face of the left astragalus (characters 107 and 108). Abbreviations: l.r.p.t.: lateral ridge of the proximal trochlea, l.b.: laterodistal border. Scalebar = 2 cm.

A. It is concave overall with a long and shallow curved notch.
B. It is concave overall with a sinusoidal outline.
C. It is slightly concave with a short and well-marked notch closer to the lateral ridge of the proximal trochlea.
D. It is slightly concave with a short and deep notch located between the proximal and distal articular surfaces.
E. It is slightly convex with a small pointed medial process.
F. It is nearly flat with a slightly sinusoidal outline.

##### 108. Morphology of the lateral ridge of the proximal trochlea in the anterior view. (Fig. 64)

A. It is wide and short.
B. It is slender and long.

##### 109. Morphology of the laterodistal articular area in posterior view. (Fig. 65)

**Figure 65.**
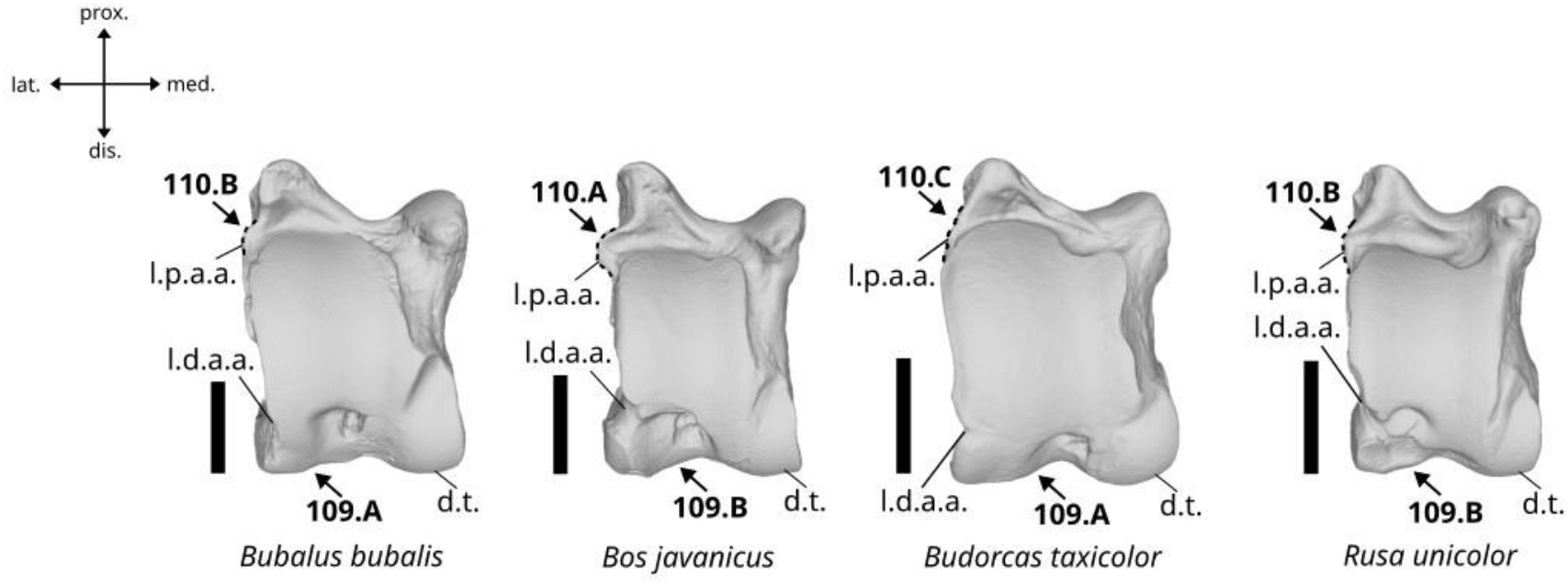
Character states visible on the posterior face of the left astragalus (characters 109 and 110). Abbreviations: l.d.a.a.: laterodistal articular area, l.p.a.a.: lateroproximal articular area, d.t.: distal trochlea. Scalebar = 2 cm.

A. Linked to the articular facet of the distal trochlea.
B. Separated from the articular facet of the distal trochlea.

##### 110. Outline of the latero-proximal area of the bone in posterior view. (Fig. 65)

A. Presenting a well-developed triangular expansion.
B. Presenting a moderately or weakly developed triangular expansion.
C. Flat or slightly curved.

##### 111. Morphology of the medial bulge in medial view. (Fig. 66)

**Figure 66.**
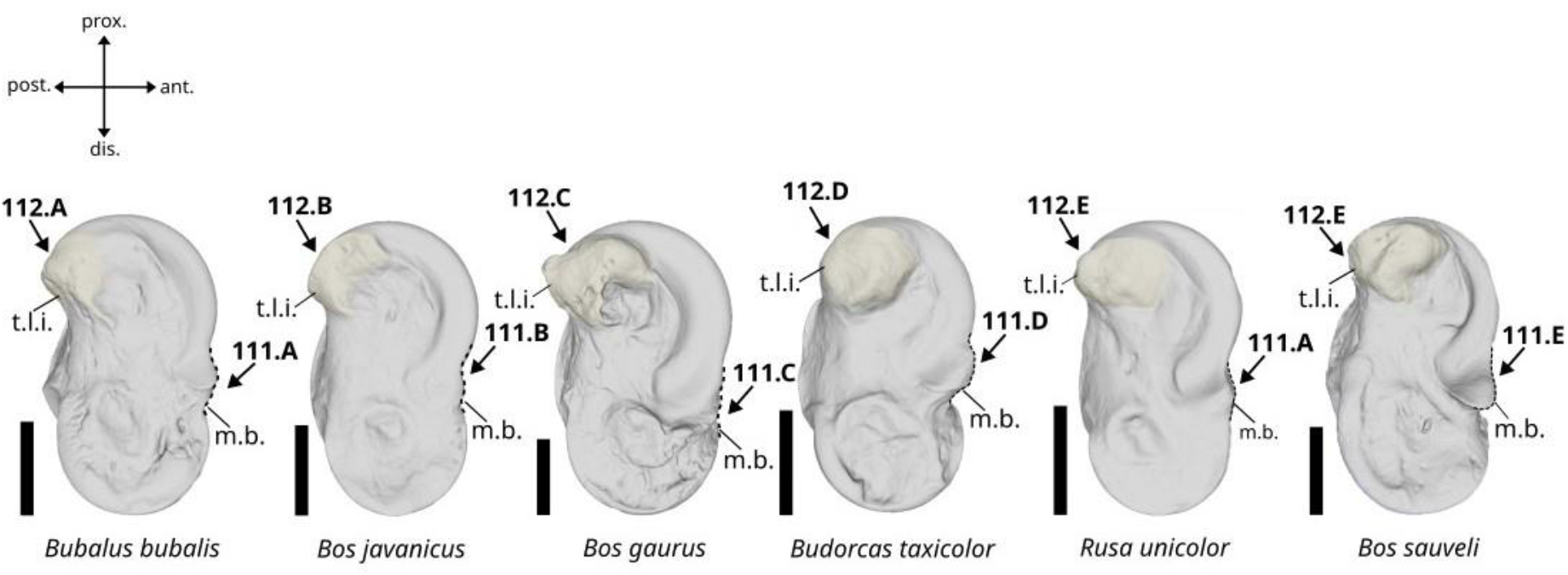
Character states visible on the medial face of the left astragalus (characters 111 and 112). Abbreviation: m.b.: medial bulge, t.l.i.: talotibial ligament insertion. Scalebar = 2 cm.

A. It is well marked with a pointed apex.
B. It is slightly marked with a sinusoidal margin.
C. It is absent.
D. It is well-marked and of truncated triangular shape with a flat anterior margin.
E. It is well-marked with a beak shape oriented in anterodistal direction.

##### 112. Morphology of the talotibial ligament insertion in medial view. (Fig. 66)

A. The ligament’s tuberosity is well developed and oriented in posterior direction with an acute apex.
B. The ligament’s tuberosity is weakly developed and oriented in posterior direction with an acute apex.
C. The ligament’s tuberosity is well-developed and oriented in postero-proximal direction with a beak-shaped apex.
D. The ligament’s tuberosity is very well developed, not projected in any direction, and presents a blunt and rounded apex.
E. The ligament’s tuberosity is well developed and oriented in posterior direction with a curved posterior margin.

##### 113. Morphology of the latero-distal trochlea in lateral view. (Fig. 67)

**Figure 67.**
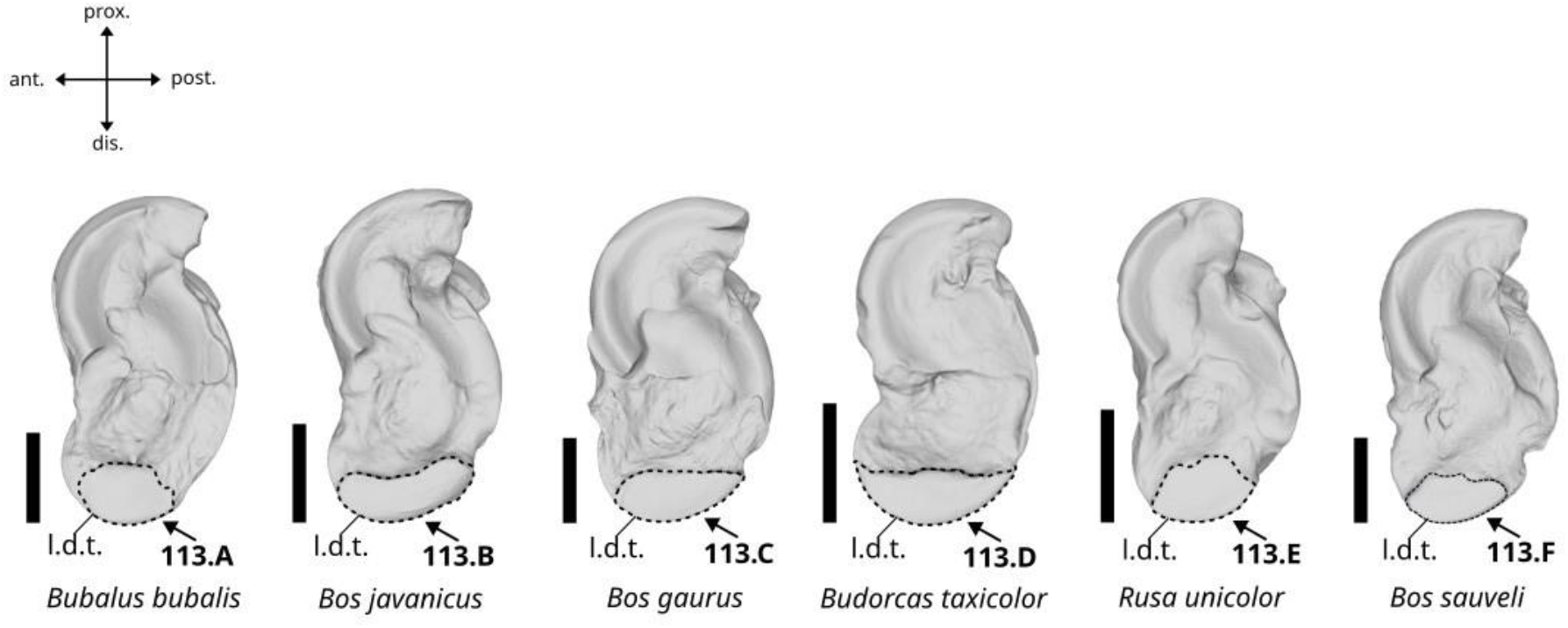
Character states visible on the lateral face of the left astragalus (character 113). Abbreviation: l.d.t.: latero-distal trochlea. Scalebar = 2 cm.

A. It is short, broad, moderately extended in proximal direction, and presents a rounded proximal margin.
B. It is elongated, moderately extended on the anterior and posterior areas of the bone, and presents a concave proximal margin.
C. It is short, wide, moderately extended on the posterior area of the bone, and presents a more or less straight proximal margin.
D. It is elongated, strongly extended on the anterior and posterior areas of the bone, and presents a sinusoidal proximal margin.
E. It is very short, broad, not extended on the anterior and posterior areas of the bone, well-extended in proximal direction, and presents a sinusoidal proximal margin.
F. It is short and broad, poorly extended on the anterior and posterior areas of the bone, moderately extended on the proximal area of the bone, and presents a sinusoidal proximal margin.

#### Characters of the navicular-cuboid

##### 114. Morphology of the median postero-proximal articular facet in antero-proximal view. (Fig. 68)

**Figure 68.**
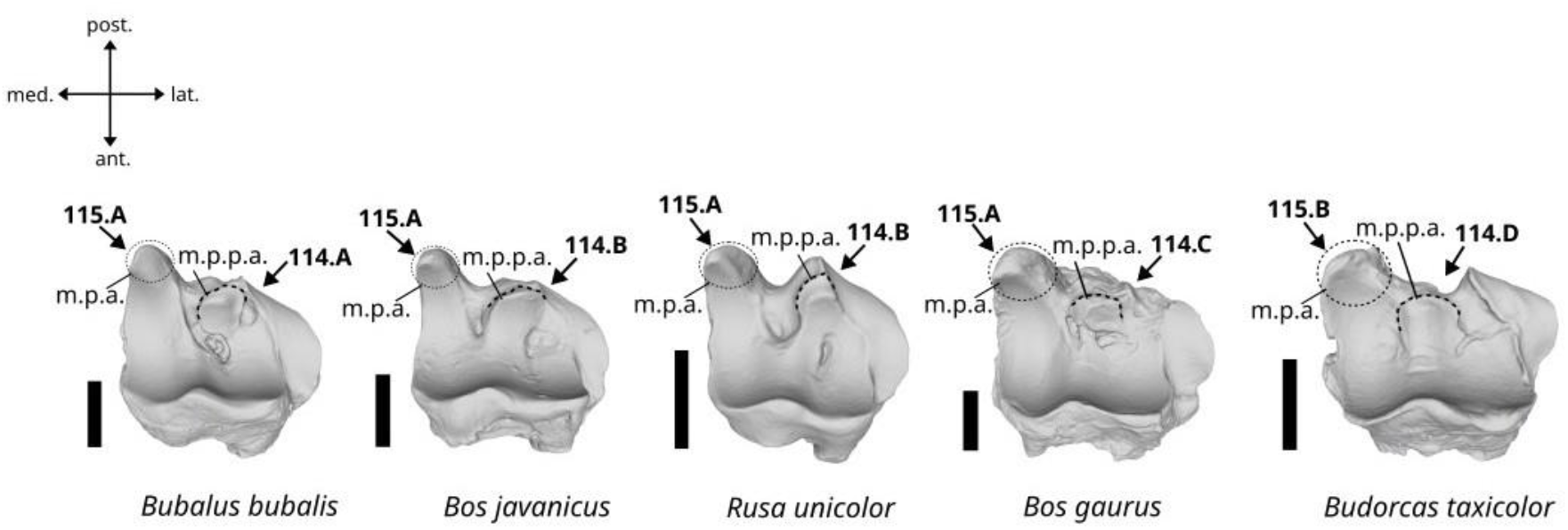
Character states visible on the antero-proximal face of the left navicular-cuboid (characters 114, and 115). Abbreviations: m.p.p.a.: median postero-proximal articular facet, m.p.a.: medio-posterior articular surface. Scalebar = 2 cm.

A. It is high and partly individualized from the rest of the proximal articular surface.
B. It is high and nearly fully in contact with the main proximal articular surface.
C. It is short, and nearly fully individualized from the rest of the proximal articular surface.
D. It is short, and half of its distal area is in contact with the main proximal articular surface.

##### 115. Morphology of the medio-posterior articular surface in antero-proximal view. (Fig. 68)

A. It is well-projected in dorsal direction and presents a pointed apex.
B. It is weakly projected in dorsal direction and presents a blunt apex.

##### 116. Morphology of the articular surface for the calcaneus bone in proximal view. (Fig. 69)

**Figure 69.**
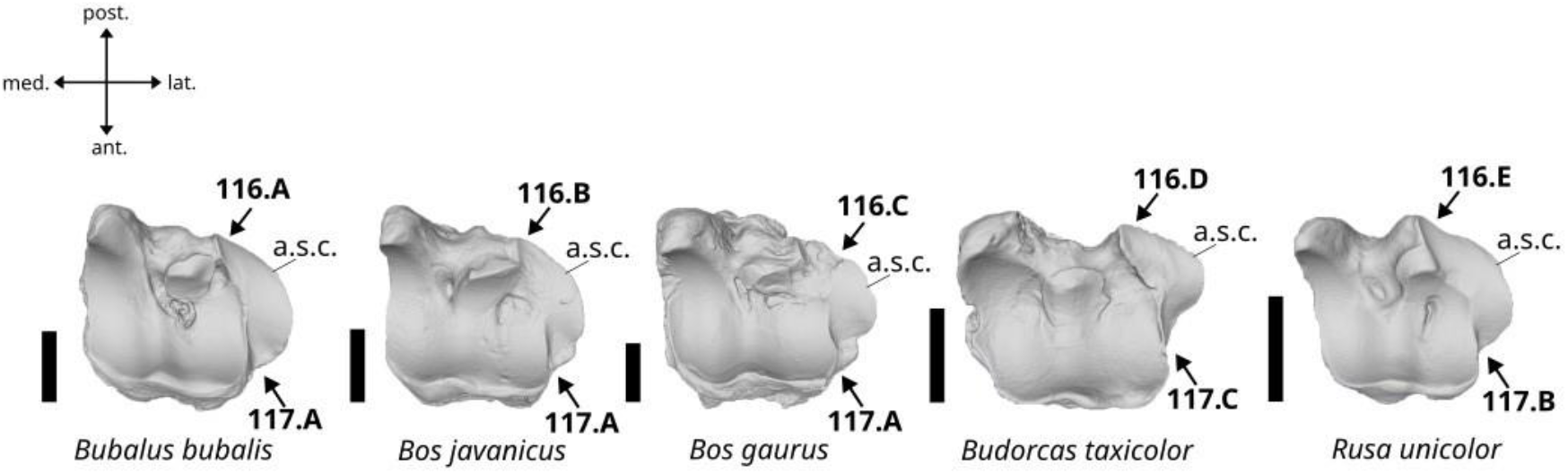
Character states visible on the proximal face of the left navicular-cuboid (characters 116 and 117). Abbreviation: a.s.c.: articular surface for the calcaneus. Scalebar = 2 cm.

A. The posterior area is well-extended in the medial area of the bone, and the whole facet presents a curved lateral margin.
B. The posterior area is well-extended in the medial area of the bone, and the whole facet presents a sinusoidal lateral margin.
C. The posterior area is weakly extended in the medial area of the bone, and the whole facet presents a sinusoidal lateral margin.
D. The posterior area is strongly extended in the medial area of the bone, and the whole facet presents a triangular lateral margin.
E. The posterior area is strongly extended in the medial area of the bone, and the whole facet presents a curved lateral margin.

##### 117. Anterior extension of the articulation for the calcaneus in proximal view. (Fig. 69)

A. It is well-extended toward the anterior limit of the bone.
B. It is moderately extended and reaches about the antero-posterior mid-length of the bone.
C. It is short and restricted to the posterior half of the bone.

##### 118. Size of the posteromedian articular facet in distal view. (Fig. 70)

**Figure 70.**
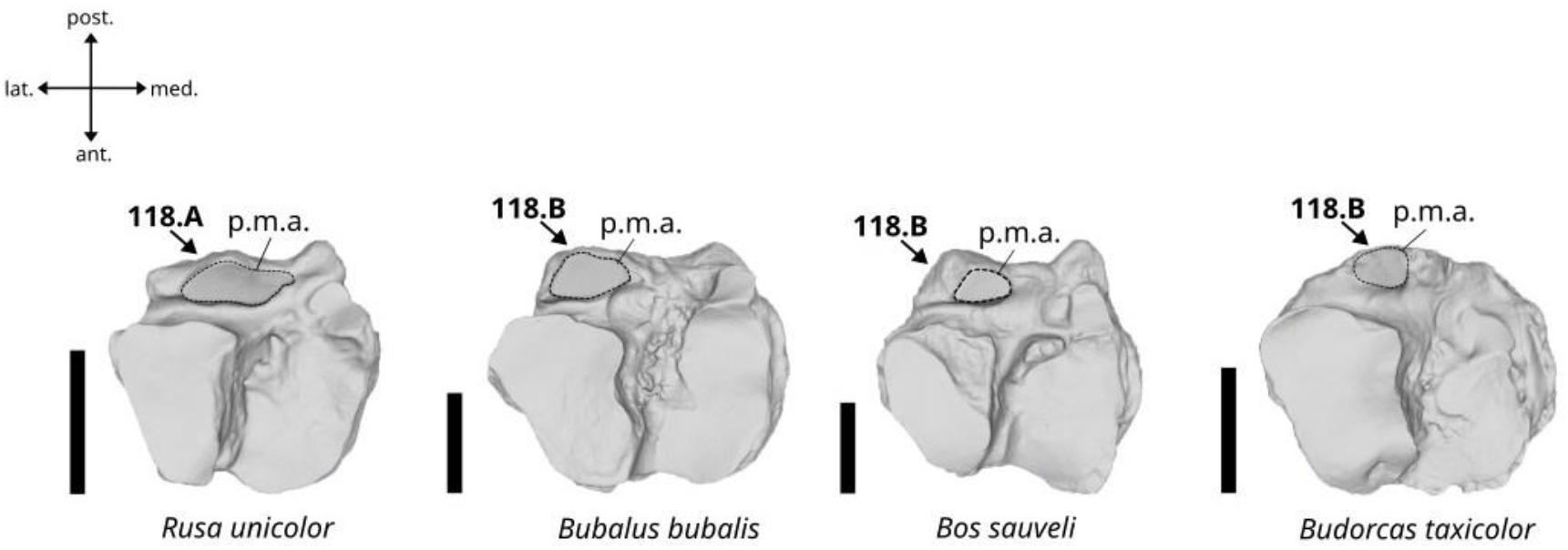
Character states visible on the distal face of the left navicular-cuboid (character 118). Abbreviation: p.m.a.: posteromedian articular facet. Scalebar: 2 cm.

A. Large.
B. Small

#### Characters of the large cuneiform

##### 119. Morphology of the medial border of the proximal articular surface in proximal view. (Fig. 71)

**Figure 71.**
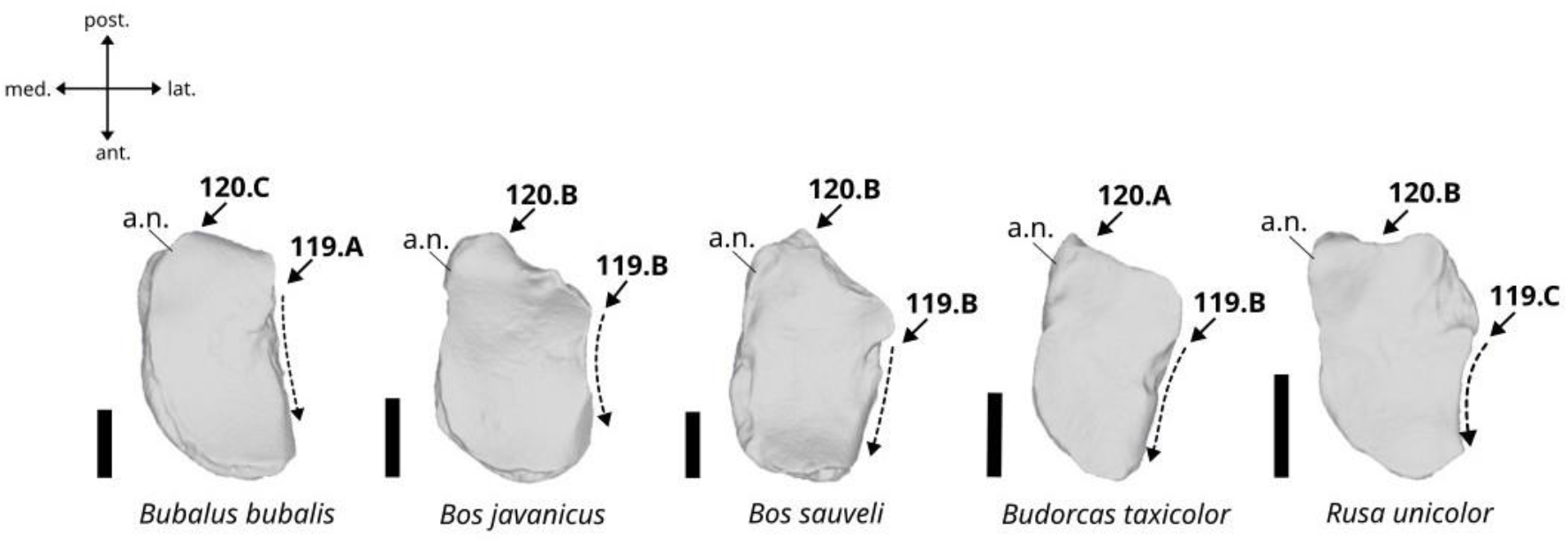
Character states visible on the proximal face of the left large cuneiform (characters 119 and 120). Abbreviation: a.n. articular surface for the navicular-cuboid. Scalebar = 1 cm.

A. It is nearly straight.
B. It is irregular.
C. It is concave.

##### 120. Morphology of the posterior border of the proximal articular surface in proximal view. (Fig. 71)

A. It is pointed and slightly projected in posterior direction.
B. Is more or less triangular with a blunt apex and well-projected in posterior direction.
C. It is convex and not projected in posterior direction.

##### 121. Morphology of the articular surface for the navicular-cuboid in medial view. (Fig. 72)

**Figure 72.**
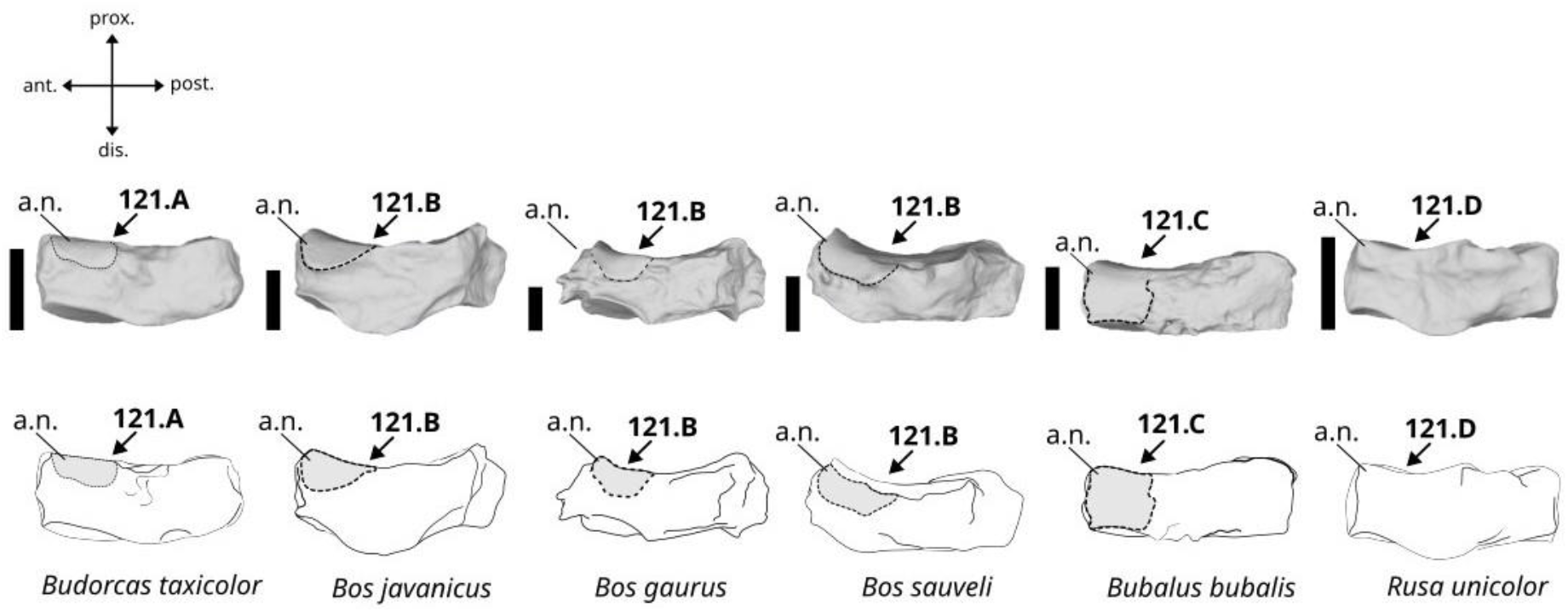
Character states visible on the medial face of the left large cuneiform (character 121). Abbreviation: a.n. articular surface for the navicular-cuboid. Scalebar = 1 cm.

A. Slightly extended in distal direction.
B. Well-extended in distal direction, reaching about half of the height of the bone.
C. Strongly extended in distal direction, and reaching the distal articular surface.
D. Absent.

#### Characters of the metatarsus

##### 122. Morphology of the interosseous canal in anterior view. (Fig. 73)

**Figure 73.**
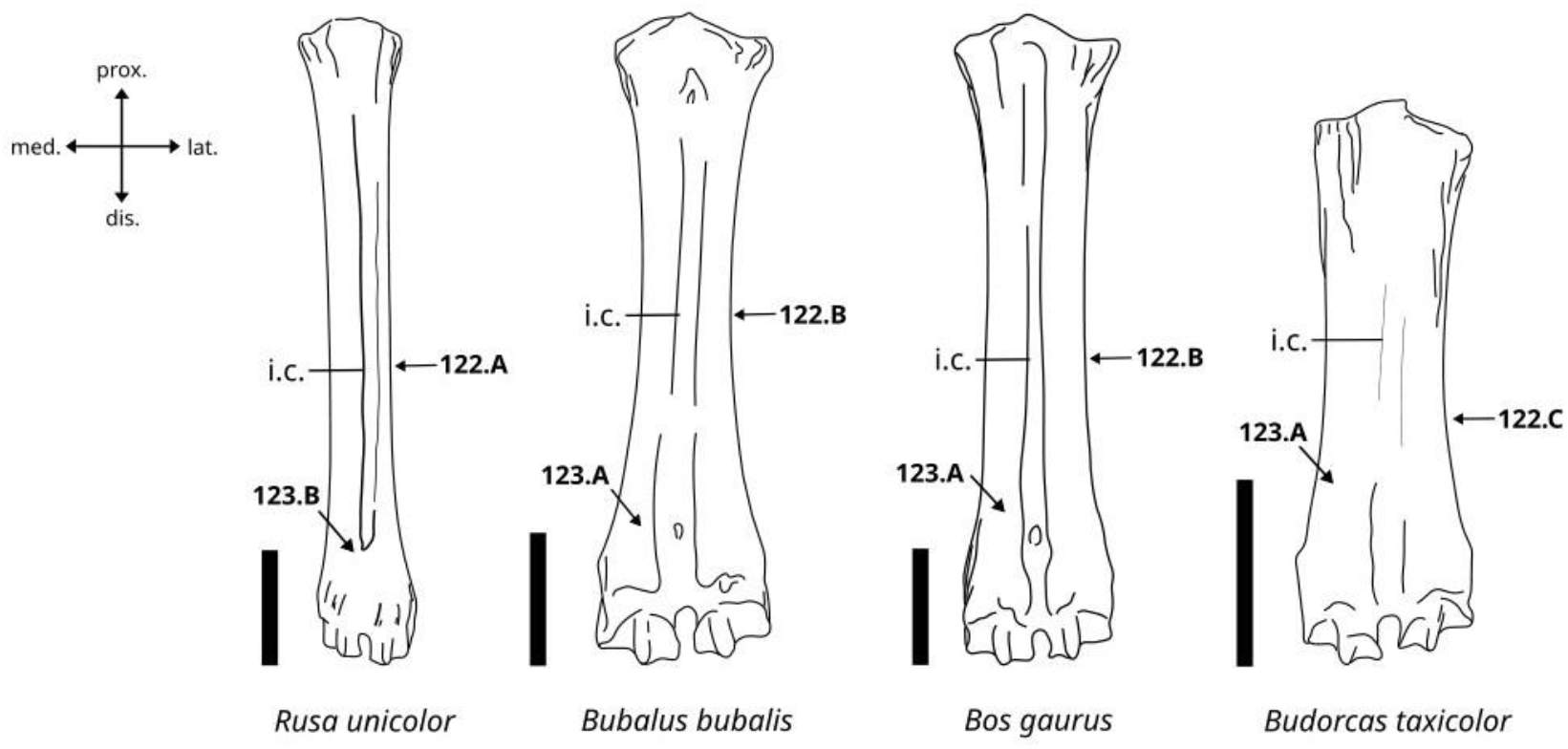
Character states visible on the anterior face of the left metatarsus (characters 122 and 123). Abbreviation: i.c.: interosseous canal. Scalebar = 5 cm.

A. Well-marked, deep, and present along most of the length of the bone.
B. Slightly marked, shallow, and present along most of the length of the bone.
C. Well-marked, deep, and restricted to the distal area of the bone.

##### 123. Morphology of the distal area of the interosseous canal in anterior view. (Fig. 73)

A. The groove is continuously open from the median area to the interarticular incisor.
B. The groove is closed in its distal part.

##### 124. Morphology of the posterolateral tuberosity in proximal view. (Fig. 74)

**Figure 74.**
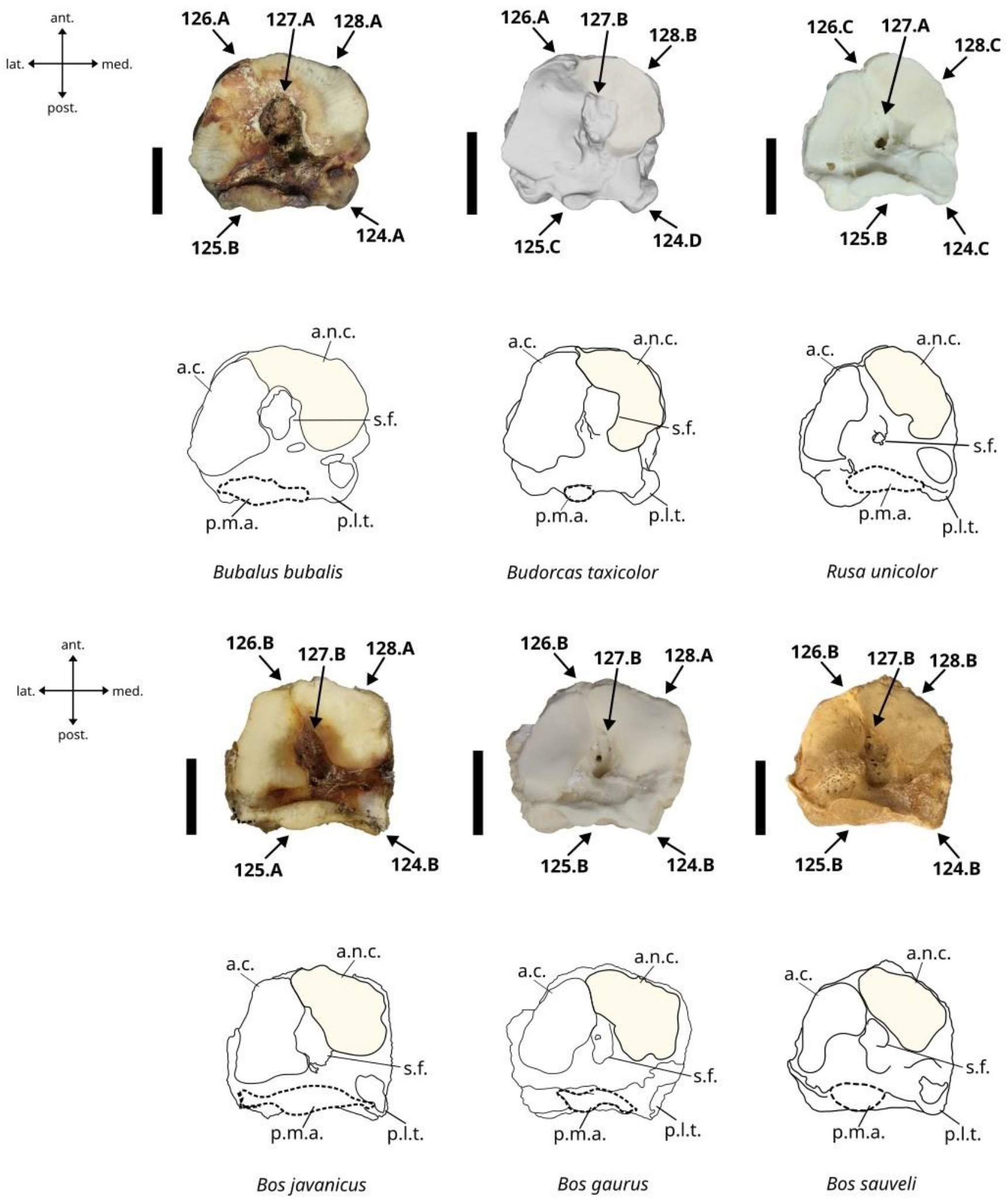
Character states visible on the proximal face of the left metatarsus (characters 124, 125, 126, 127, and 128). Abbreviations: a.n.c.: articular surface for the navicular-cuboid (medial), a.c.: articular surface for the cuneiform (lateral), s.f.: synovial fossa, p.l.t.: postero-lateral tuberosity, p.m.a.: postero-median articular facet. Scalebar = 2 cm.

A. The tuberosity is well-developed with a globular posterior apex.
B. The tuberosity is moderately developed with a pointed posterior apex.
C. The tuberosity is moderately developed with a curved posterior apex.
D. The tuberosity is well-developed with a truncated triangular posterior apex.

##### 125. Shape of the postero-median articular facet in proximal view. (Fig. 74)

A. It is strongly extended in medial and lateral directions.
B. It is moderately extended in medial and lateral directions.
C. It is reduced, small, and circular.

##### 126. Contact between the anterolateral and anteromedial articular surfaces in proximal view. (Fig. 74)

A. They are in contact without any subdivision.
B. They are in contact but remain divided into two facets by a small ridge.
C. They are divided into two facets.

##### 127. Morphology of the synovial fossa in proximal view. (Fig. 74)

A. Deep.
B. Shallow.

##### 128. Expansion of the anterolateral area of the proximal articular surface in proximal view. (Fig. 74)

A. Well-expanded in lateral direction.
B. Moderately expanded in lateral direction.
C. Not expanded in lateral direction.

#### Characters of the first phalanx

##### 129. Morphology of the distal articular surface in dorsal view. (Fig. 75)

**Figure 75.**
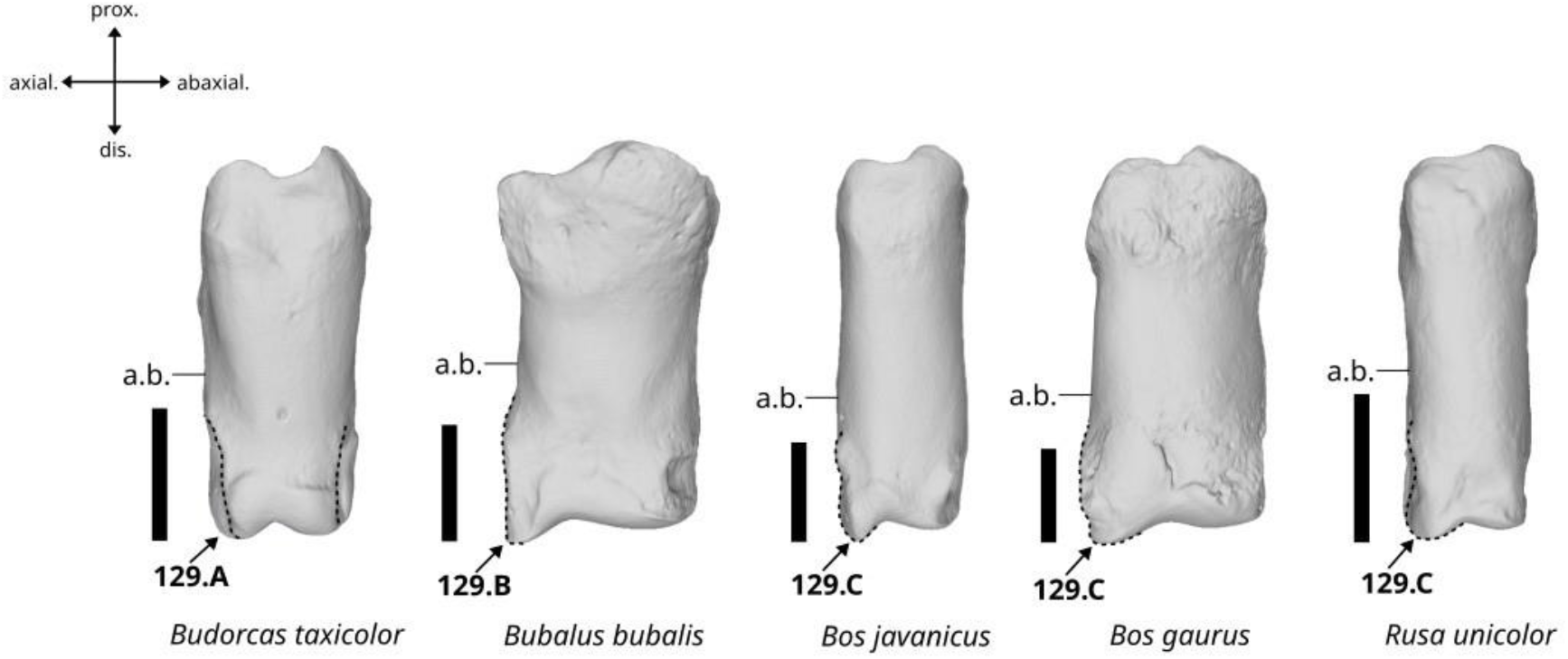
Character states visible on the dorsal face of the first phalanx (character 129). Abbreviation: a.b.: axial border. Scalebar = 2 cm.

A. The abaxial and axial area of the articular surface are of similar development. The axial margin of the articular area is sinusoidal.
B. The axial area presents an axial projection of moderate length pointing in distal direction. The axial margin of the articular area is straight.
C. The axial area presents an axial projection of moderate length pointing in distal direction. The axial margin of the articular area is sinusoidal.

##### 130. Morphology of the palmar margin in axial view. (Fig. 76)

**Figure 76.**
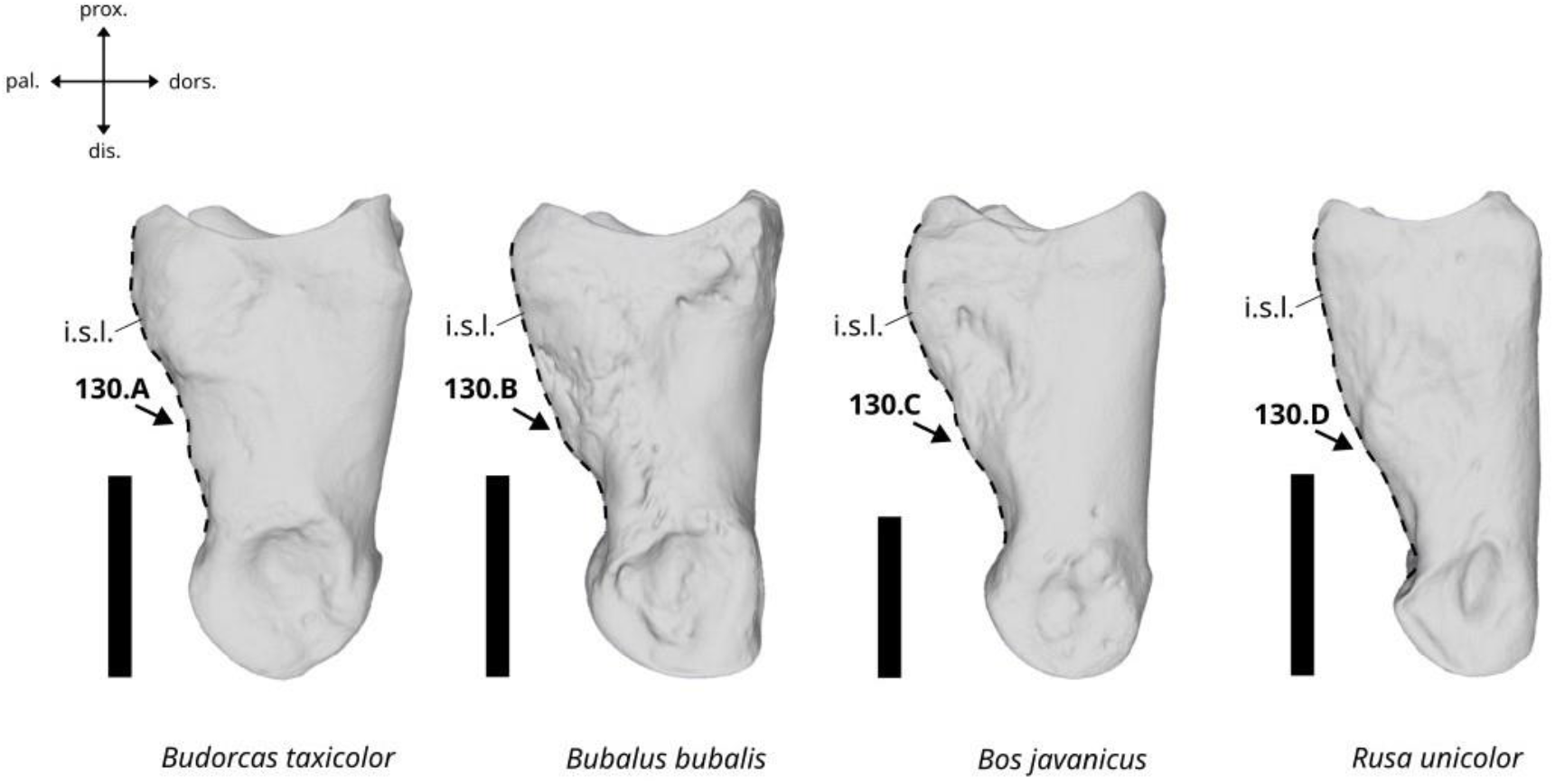
Character states visible on the axial face of the first phalanx (character 130). Abbreviation: i.s.l.: imprint of the sesamoidean ligament. Scalebar = 2 cm.

A. The imprint of the sesamoidean ligament is well-developed. The palmar face presents a sinusoidal distal margin.
B. The imprint of the sesamoidean ligament is well-developed. The palmar face presents a deep distal depression.
C. The imprint of the sesamoidean ligament is well-developed. The palmar face presents a shallow distal depression.
D. The imprint of the sesamoidean ligament is weakly developed. The palmar face presents an oblique distal margin.

##### 131. Morphology of the insertion surface in palmar view. (Fig. 77)

**Figure 77.**
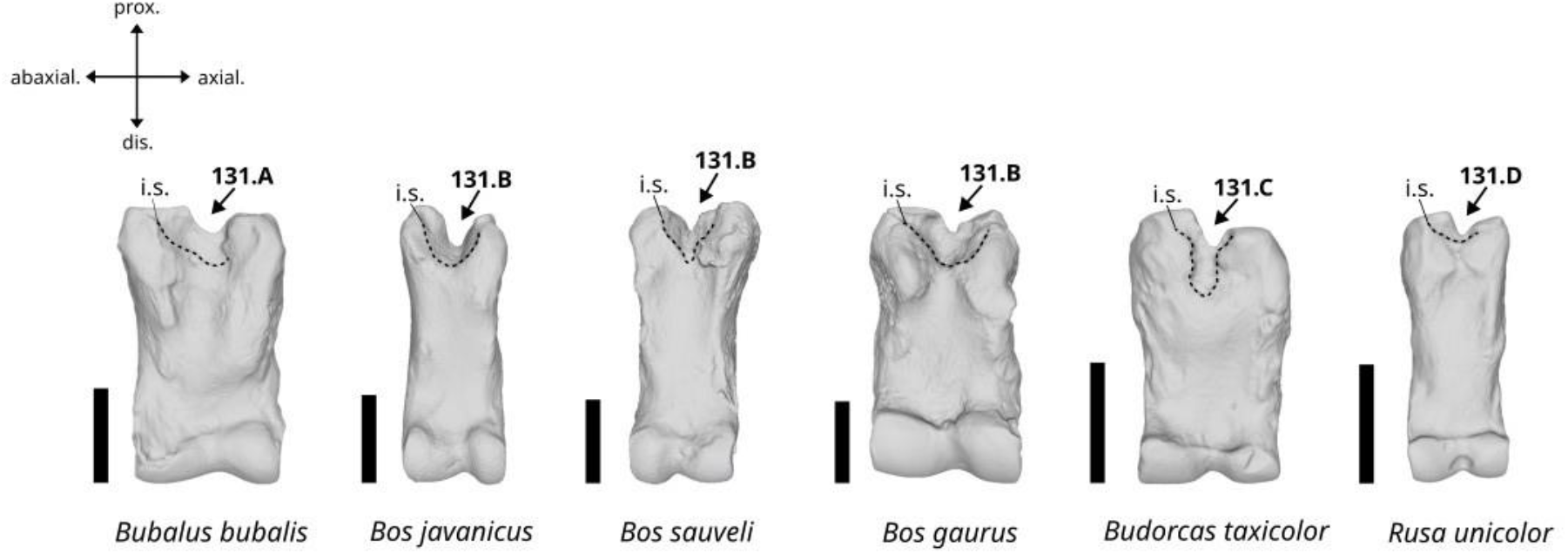
Character states visible on the palmar face of the first phalanx (character 131). Abbreviation: i.s.: insertion surface. Scalebar = 2 cm.

A. It is moderately extended in distal direction with an irregular distal margin.
B. It is moderately extended in distal direction with a more or less curved distal margin.
C. It is well-extended in distal direction with a deep and oblong distal margin.
D. It is weakly extended in distal direction with a curved distal margin.

#### Characters of the second phalanx

##### 132. Shape of the abaxio-proximal margins of the bone in dorsal view. (Fig. 78)

**Figure 78.**
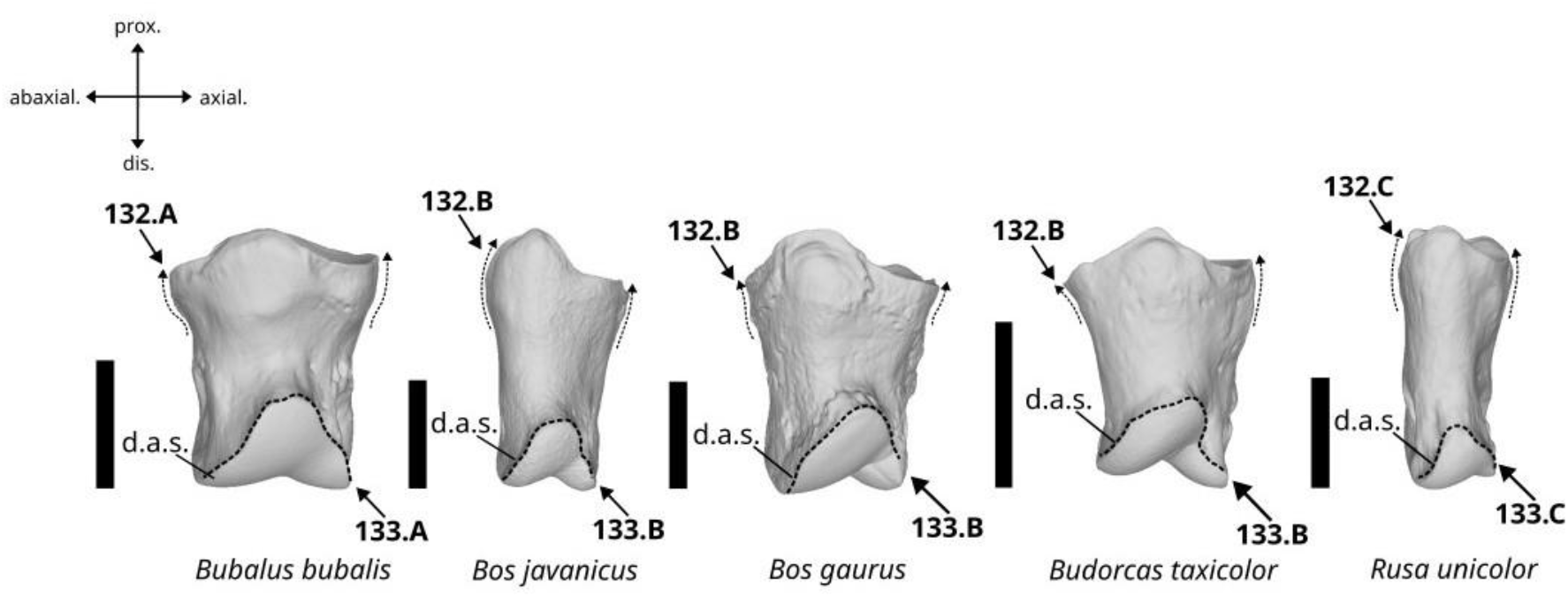
Character states visible on the dorsal face of the second phalanx (characters 132 and 133). Abbreviation: d.a.s.: distal articular surface. Scalebar = 2 cm.

A. The proximal area is strongly enlarged with slightly angular lateral edges.
B. The proximal area is enlarged with more or less curved lateral margins.
C. The proximal area is slightly enlarged with more or less curved lateral margins.

##### 133. Morphology of the distal articular surface in dorsal view. (Fig. 78)

A. The articular surface is broad and triangular with a nearly flat ventral margin.
B. The articular surface is broad and triangular with a strongly notched ventral margin.
C. The articular surface is thin and triangular with a nearly flat ventral margin.

##### 134. Morphology of the abaxial insertion tuberosity in palmar view. (Fig. 79)

**Figure 79.**
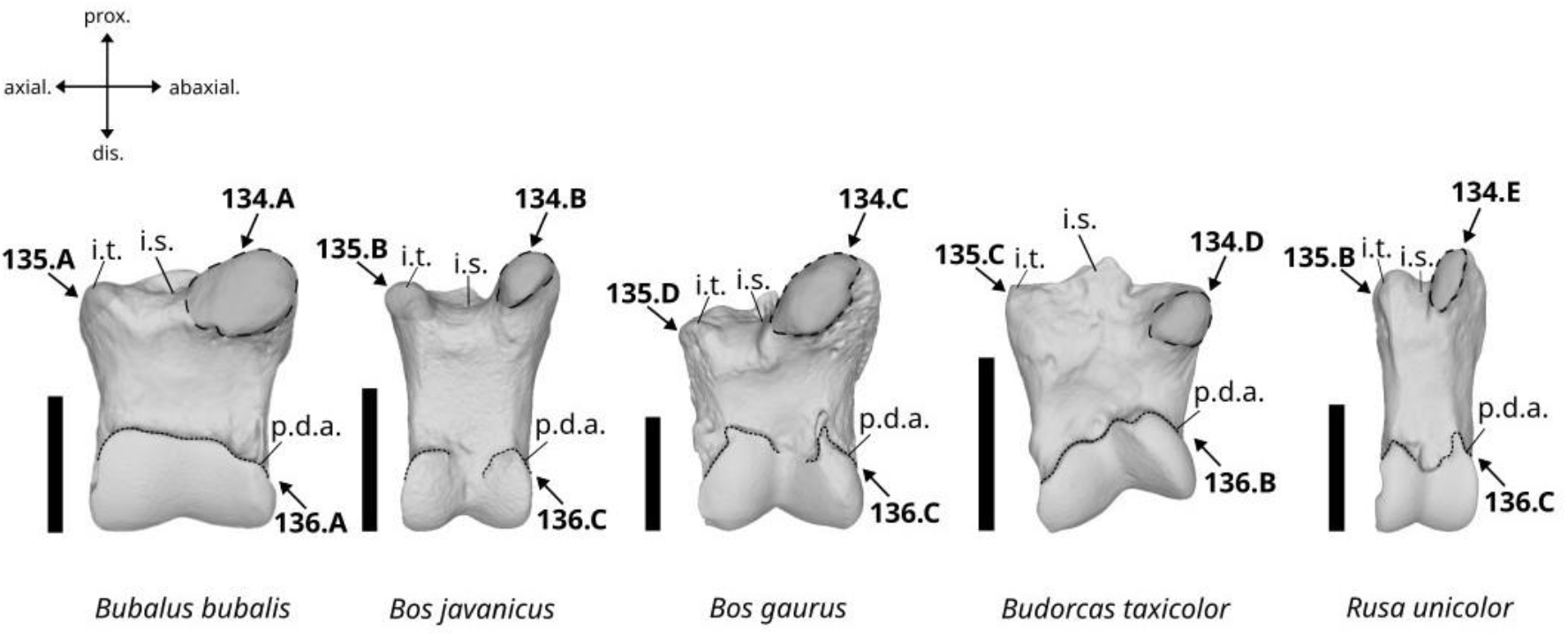
Character states visible on the palmar face of the second phalanx (characters 134, 135 and 136). Abbreviations: i.t.: insertion tubercle, i.s.: insertion surface, p.d.a.: palmarodistal articular surface. Scalebar = 2 cm.

A. The abaxial insertion tuberosity is very well-developed, well-delimited, and oriented in abaxial direction.
B. The abaxial insertion tuberosity is well-developed, poorly delimited, and moderately oriented in abaxio-proximal direction.
C. The abaxial insertion tuberosity is very well-developed, well-delimited, and oriented in abaxio- proximal direction.
D. The abaxial insertion tuberosity is weakly developed, poorly marked, and slightly oriented in abaxial direction.
E. The abaxial insertion tuberosity is weakly developed and poorly marked, and oriented in proximal direction.

##### 135. Morphology of the axial insertion tuberosity in palmar view. (Fig. 79)

A. The axial insertion tuberosity is well-developed with a blunt proximal apex.
B. The axial insertion tuberosity is poorly developed with a blunt proximal apex.
C. The axial insertion tuberosity is present but poorly marked.
D. The axial insertion tuberosity is nearly absent.

##### 136. Morphology of the palmarodistal articular surface in palmar view. (Fig. 79)

A. It is moderately extended on the antero-palmar surface of the bone, and with a more or less straight posterior margin.
B. It is moderately extended on the antero-palmar surface of the bone, and with a sinusoidal posterior margin.
C. It is well-extended on the antero-palmar surface of the bone, and with a posterior margin presenting a more or less deep notch in its median area.

##### 137. Morphology of the posterior border of the proximal articular surface in proximal view. (Fig. 80)

**Figure 80.**
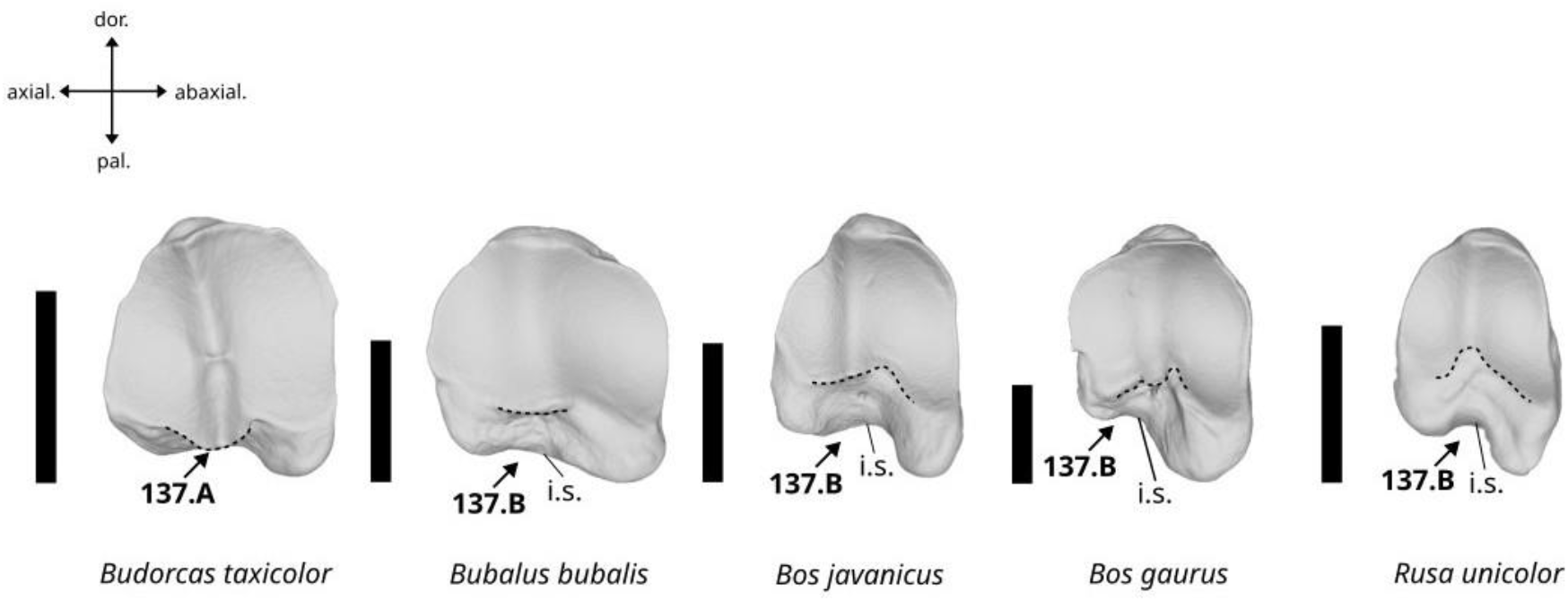
Character states visible on the proximal face of the second phalanx (character 137). Abbreviation: i.s.: insertion surface – Scalebar = 2 cm.

A. It is strongly extended in the postero-palmar direction, beyond the insertion surface of the bone.
B. It is slightly extended in the postero-palmar direction, not beyond the insertion surface of the bone.

#### Characters of the third phalanx

##### 138. Morphology of the extensor process in axial view. (Fig. 81)

**Figure 81.**
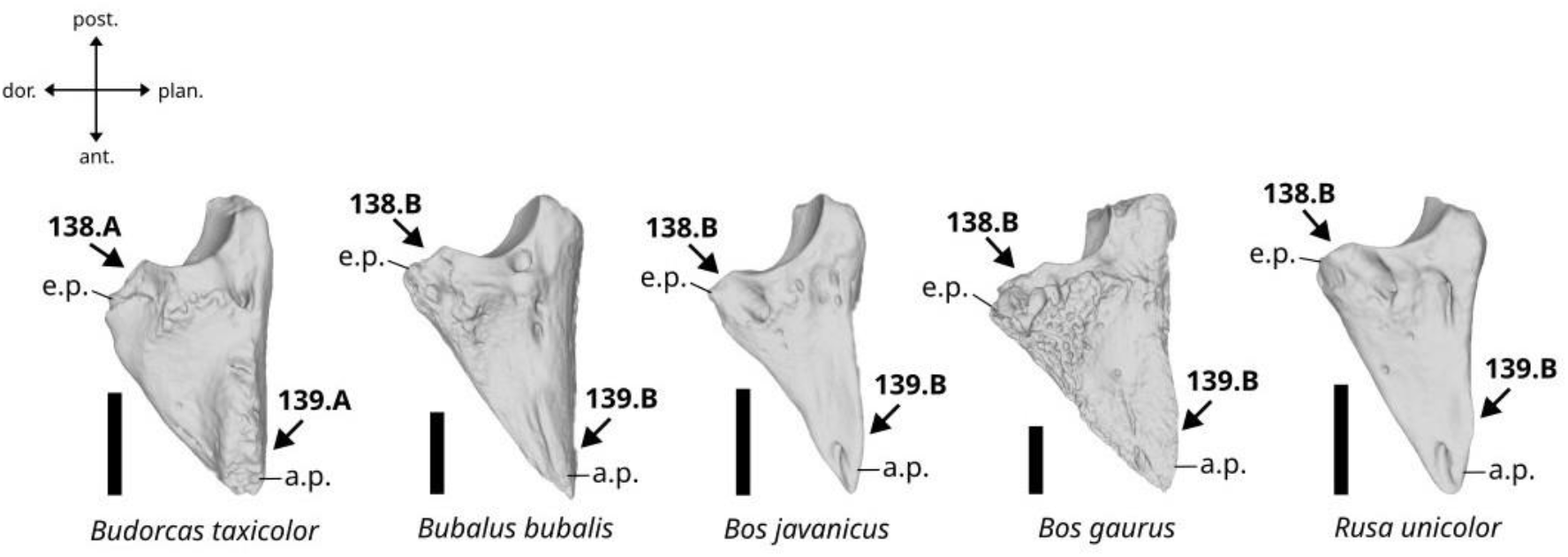
Character states visible on the axial face of the third phalanx (characters 138 and 139). Abbreviations: e.p.: extensor process, a.p.: apex of the phalanx. Scalebar = 2 cm.

A. It is truncated and short.
B. It is triangular and well-projected in postero-dorsal direction.

##### 139. Morphology of the apex of the phalanx in axial view (Fig. 81)

A. It is truncated and wide.
B. It is elongated and pointed.

##### 140. Morphology of the solar surface in palmar view. (Fig. 82)

**Figure 82.**
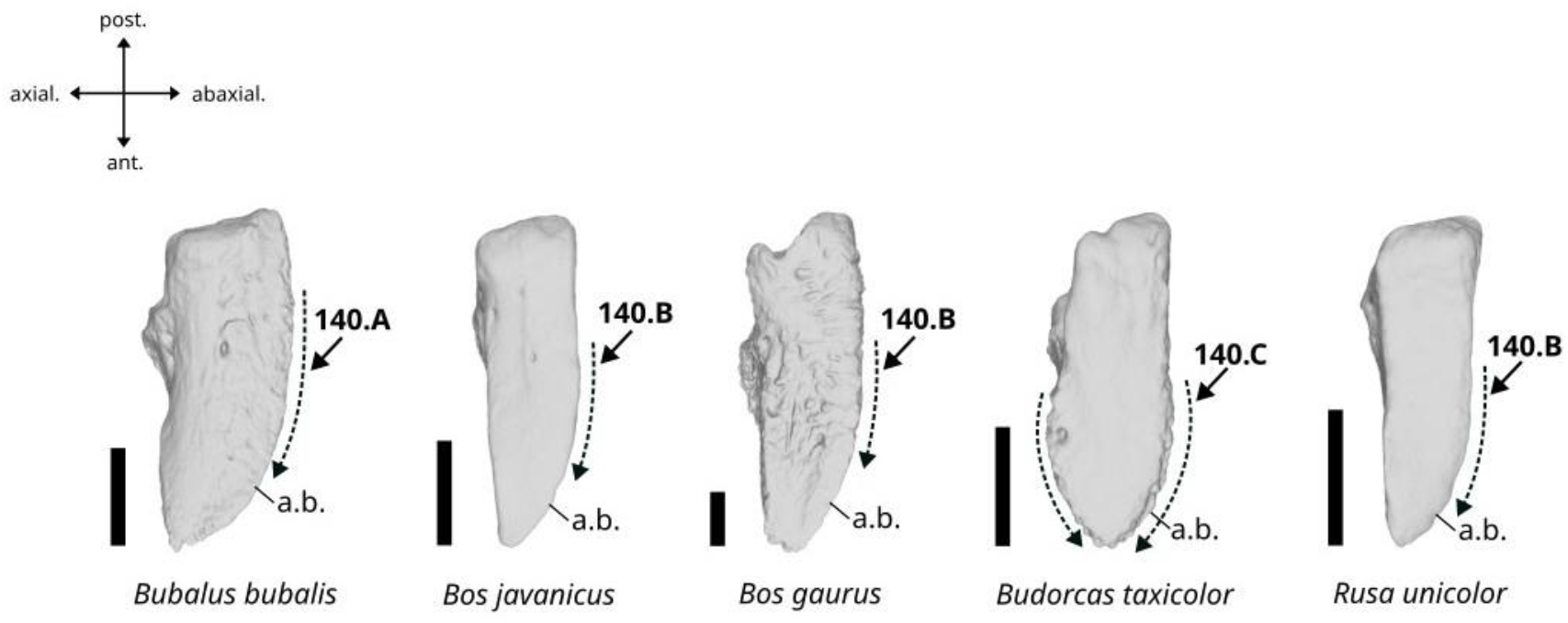
Character states visible on the palmar face of the third phalanx (character 140). Abbreviation: a.b.: abaxial border. Scalebar = 2 cm.

A. The axial border is more or less straight, and the abaxial border is curved.
B. The axial border is more or less straight, and the abaxial border is nearly straight.
C. The axial border is sinusoidal, and the abaxial border is convex, giving to the anterior area an ovate shape.

##### 141. Morphology of the abaxial border of the proximal articular surface in proximal view. (Fig. 83)

**Figure 83.**
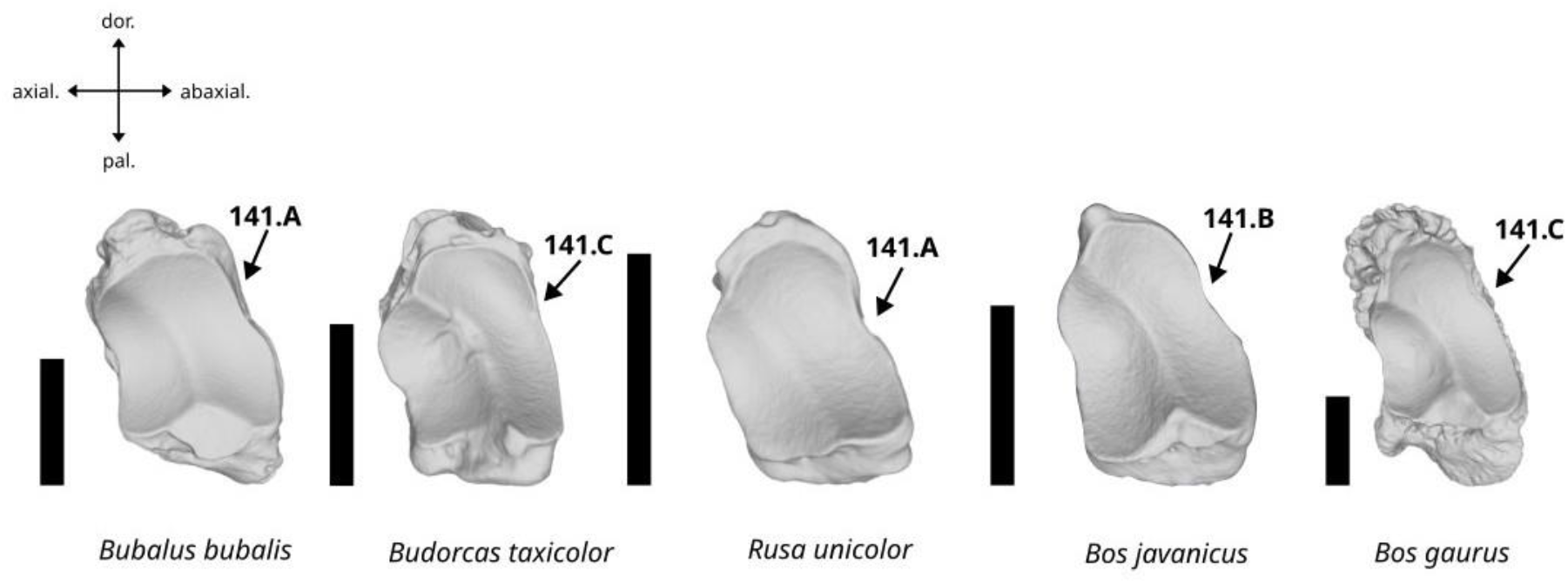
Character states visible on the proximal face of the third phalanx (character 141). Scalebar = 2 cm.

A. The abaxial border presents a deep notch.
B. The abaxial border presents a shallow notch.
C. The abaxial border do not present any notch.

#### Characters of the abaxial large sesamoid

##### 142. Shape of the bone in lateral view. (Fig. 84)

**Figure 84.**
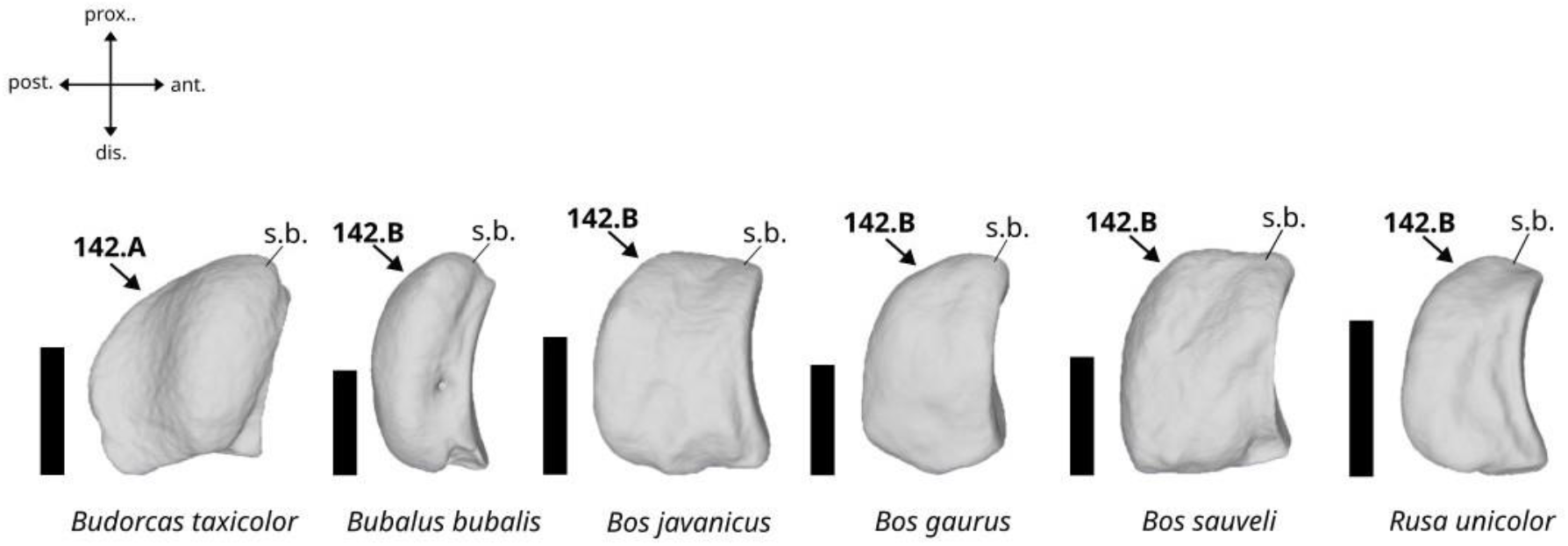
Character states visible on the abaxial face of the abaxial large sesamoid (character 142). Abbreviation: s.b.: sesamoid’s body. Scalebar = 1 cm.

A. Elongated.
B. Short.

##### 143. Morphology of the articular surface in anterior view. (Fig. 85)

**Figure 85.**
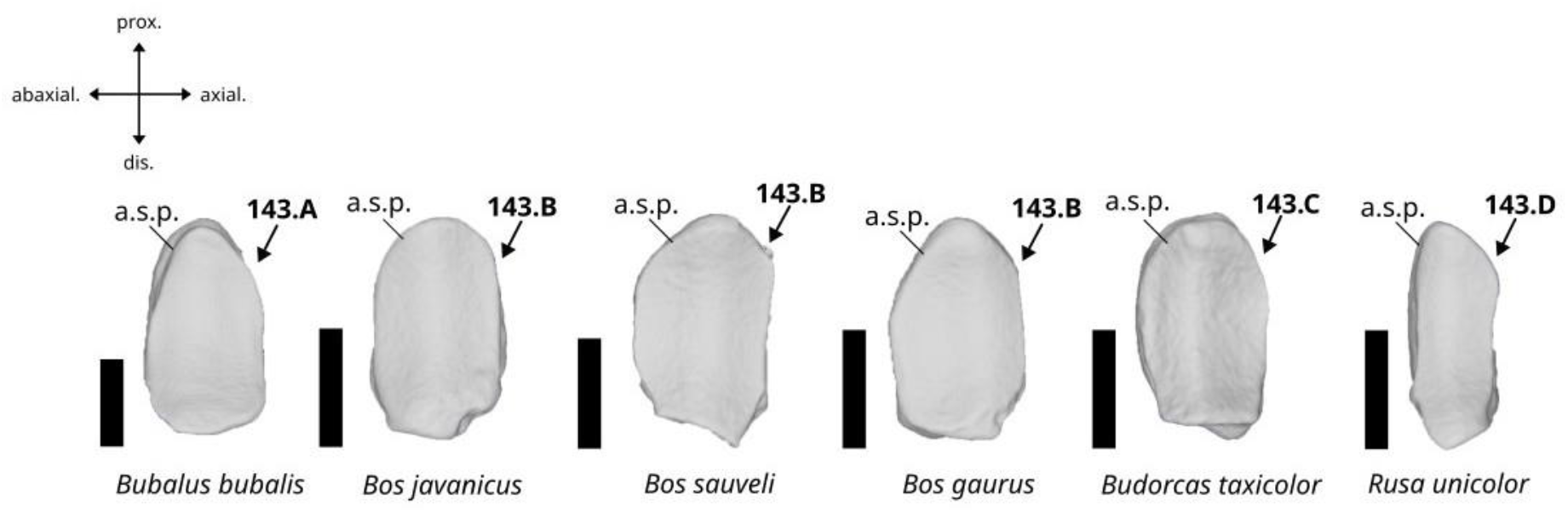
Character states visible on the anterior face of the abaxial large sesamoid (character 143). Abbreviation: a.s.p.: articular surface for the phalanx. Scalebar = 1 cm.

A. The articular surface is oval and broad. Its distal margin is curved.
B. The articular surface is oval and broad. Its distal margin is straight and oblique.
C. The articular surface is broad and oval. Its distal margin is straight and perpendicular to the main axis of the bone.
D. The articular surface is oval and slender. Its distal margin presents a pointed distal apex.

#### Characters of the axial large sesamoid

##### 144. Morphology of the ventral margin of the sesamoid’s body in lateral view. (Fig. 86)

**Figure 86.**
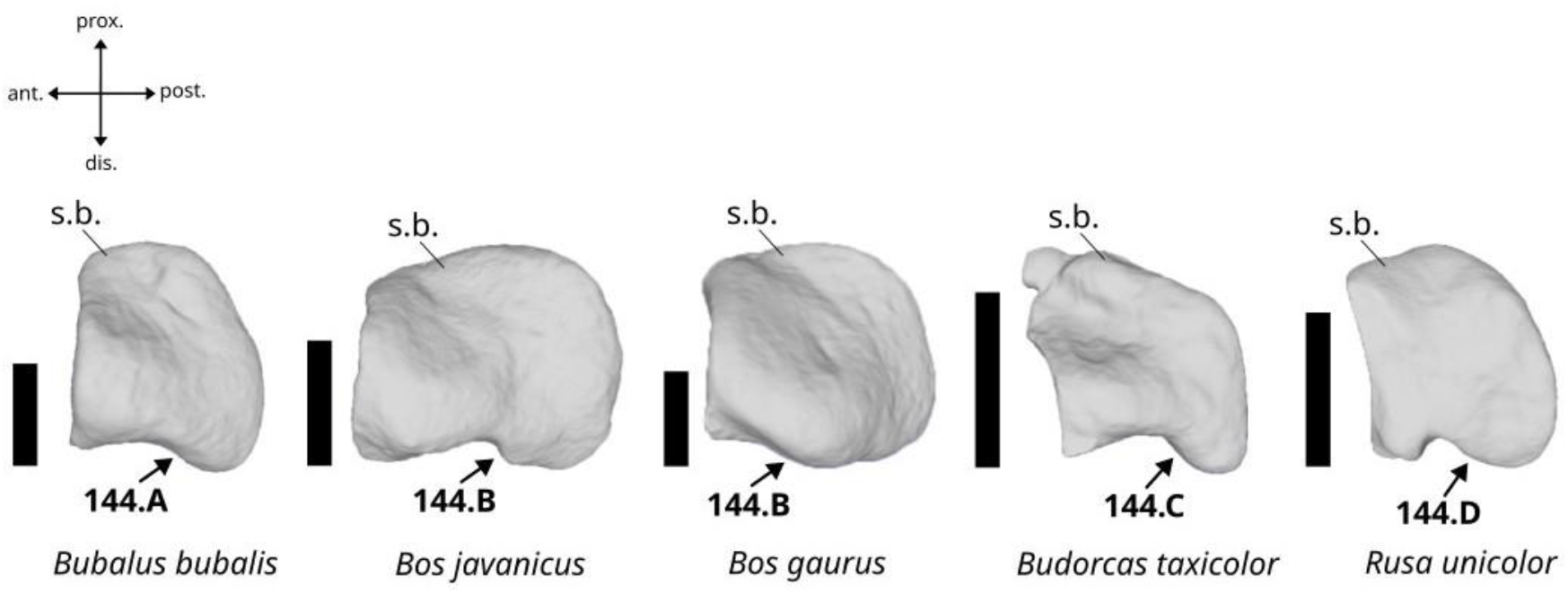
Character states visible on the axial face of the axial large sesamoid (character 144). Abbreviation: s.b.: sesamoid’s body. Scalebar = 1 cm.

A. It is slightly and uniformly curved, and slightly projected in distal direction.
B. It is sinusoidal and slightly projected in distal direction.
C. It is strongly curved on half of its length and strongly projected in distal direction.
D. It presents a small and deep notch in its axial area and is slightly projected in distal direction.

##### 145. Morphology of the anterior articular surface in anterior view. (Fig. 87)

**Figure 87.**
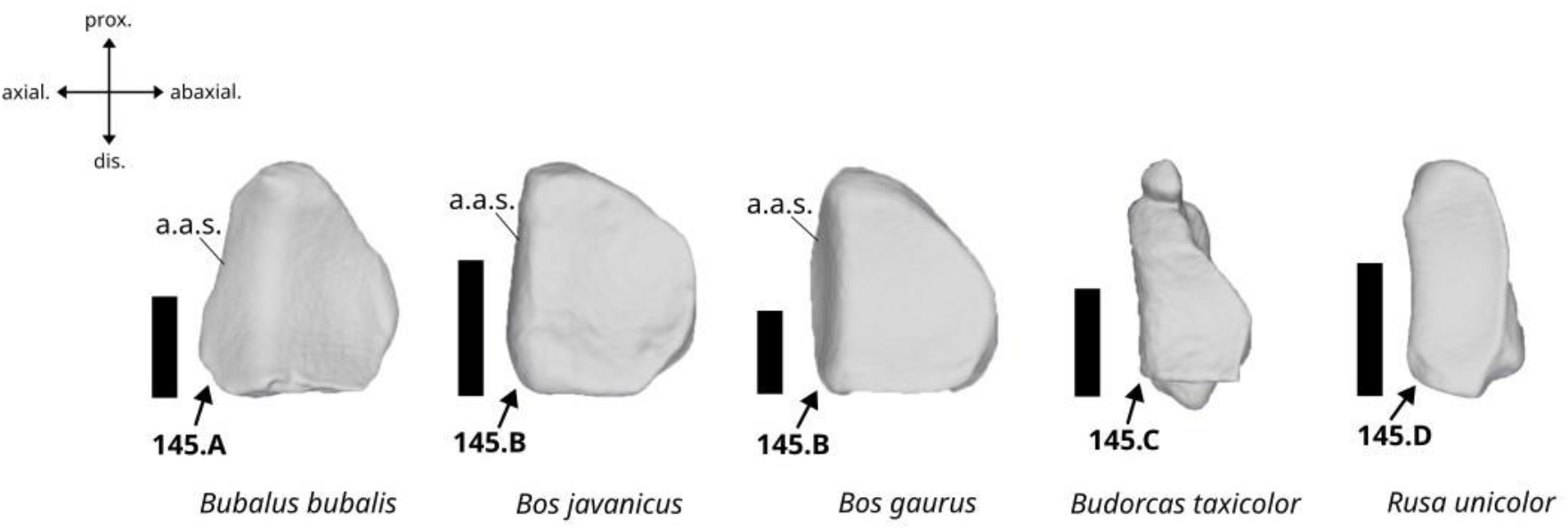
Character states visible on the anterior face of the axial large sesamoid (character 145). Abbreviation: a.a.s.: axial articular surface. Scalebar = 1 cm.

A. The axial articular surface is well-developed, with more or less curved axial margin.
B. The axial articular surface is weakly developed, with more or less straight axial margin.
C. The axial articular surface is absent, and the axial margin of the abaxial articular surface is straight.
D. The axial articular surface is absent, and the axial margin of the abaxial articular surface is concave.

#### Characters of the small sesamoid

##### 146. Shape of the sesamoid’s body in posterior view. (Fig. 88)

**Figure 88.**
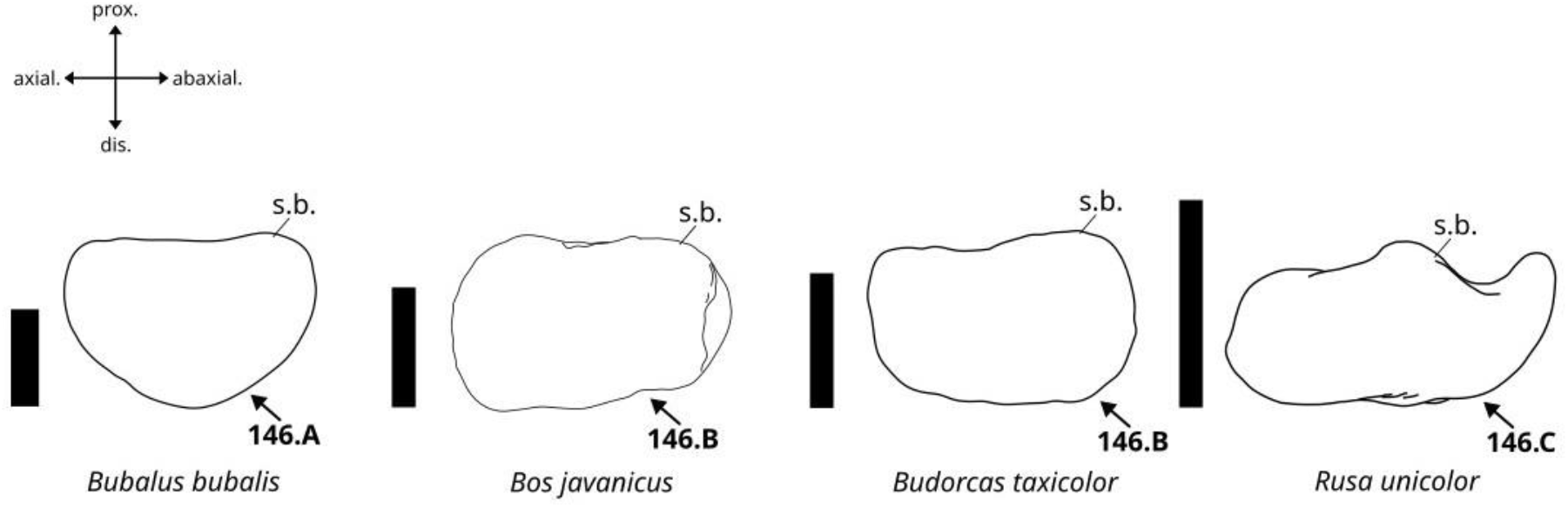
Character states visible on the posterior face of the small sesamoid (character 146). Abbreviation: s.b.: sesamoid’s body. Scalebar = 1 cm.

A. Robust and semi-circular.
B. Robust and more or less rectangular.
C. Thin, elongated, and more or less rectangular.

##### 147. Morphology of the anterior articular surface in anterior view. (Fig. 89)

**Figure 89.**
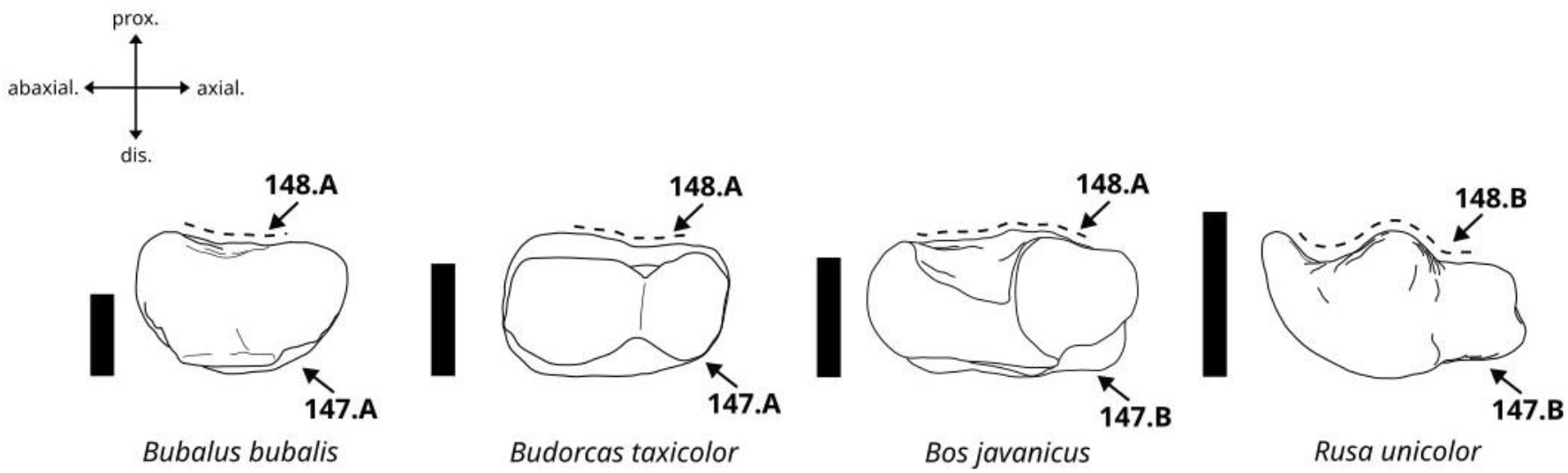
Character states visible on the anterior face of the small sesamoid (characters 147, and 148). Scalebar = 1 cm.

A. The articular facets are shallow and poorly individualized.
B. The articular facets are well-individualized and separated by a well-marked eminence.

##### 148. Morphology of the proximal border of the anterior articular surface in anterior view. (Fig. 89)

A. More or less straight.
B. Sinusoidal.

### II) Variability of the character states across the taxa

The frequency of the recorded character states across different taxonomic ranks (Supplementary Material 05; Chantasri et al., 2026) shows very limited polymorphism between the taxa (**Fig. 90**), and many characters present very strong discriminative power, with frequencies above 95%. In the dataset, counting each character state reaching a frequency of at least 95% within the taxon they characterize, 52 characters carry a state unique to a family, 128 to a subfamily, 177 to a genus, and 168 to a species. These character states are, for most of them, unique to several ranks, because one family (Cervidae), two subfamilies (Caprinae and Cervinae), and three genera (*Budorcas*, *Rusa*, and *Bubalus*) are each represented by a single species in our dataset. For higher taxonomic ranks, it thus only makes sense to fully consider the character states unique to the taxonomic ranks represented by more than one species (three in this case): Bovidae (7 character states: 5:B, 9:B, 32:B, 47:B, 100:B, 123:A, 148:A), Bovinae (13 character states: 18:A, 21:A, 22:A, 27:A, 29:C, 49:C, 59:A, 66:B, 72:A, 77:A, 83:B, 85:A, 122:B), and *Bos* (20 character states: 24:B, 31:C, 35:B, 36:B, 38:C, 40:C, 50:B, 58:B, 65:C, 67:B, 80:B, 92:B, 93:B, 97:B, 126:B, 130:C, 131:B, 143:B, 144:B, 145:B) (Figure 90).

**Figure 90.**
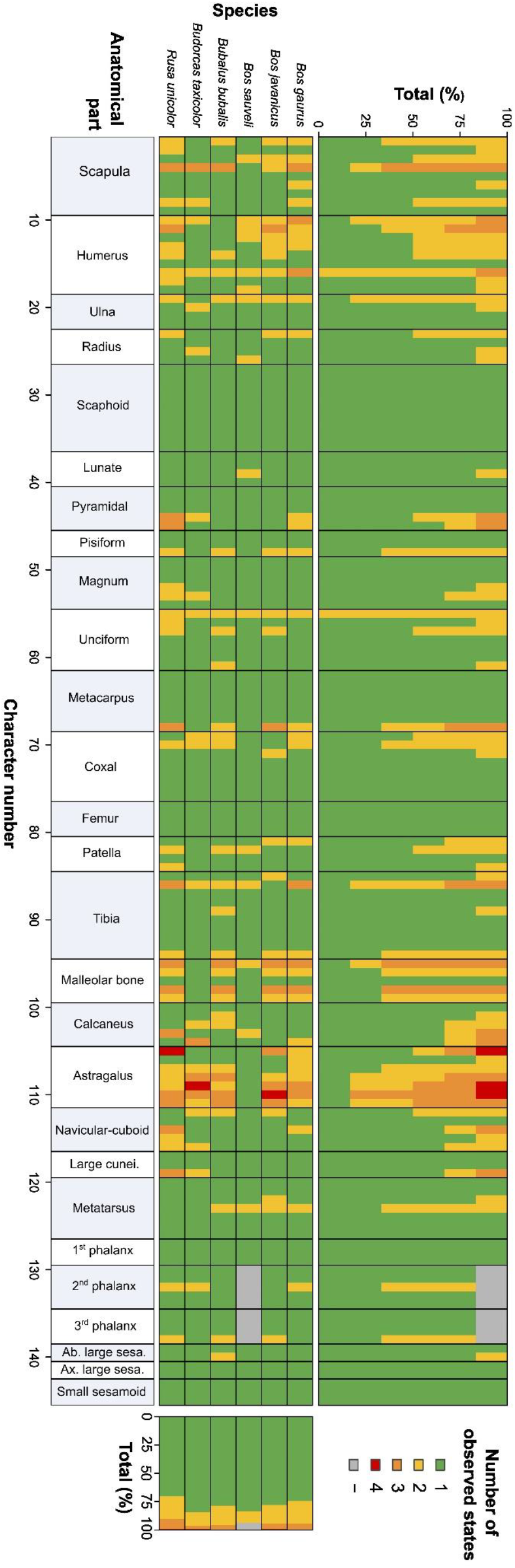
Number of observed states for each character in each taxon.

The character states unique to a species are most numerous for species that are the sole representative of a higher taxonomic rank, such as *Rusa unicolor* (45 character states); *Budorcas taxicolor* (70 character states); and *Bubalus bubalis* (42 character states) (See Supplementary Material 06 for the full list; Chantasri et al., 2026) (**Fig. 91**). Conversely, the character states unique to the species within *Bos* are much scarcer, with only two for *Bos gaurus* (17:C, 52:C), one for *Bos javanicus* (25:C), and eight for *Bos sauveli* (17:F, 32:A, 48:F, 52:F, 98:F, 107:F, 111:E, 113:F) — although the fact that this last species is only represented by three specimens makes this result potentially fragile.

**Figure 91.**
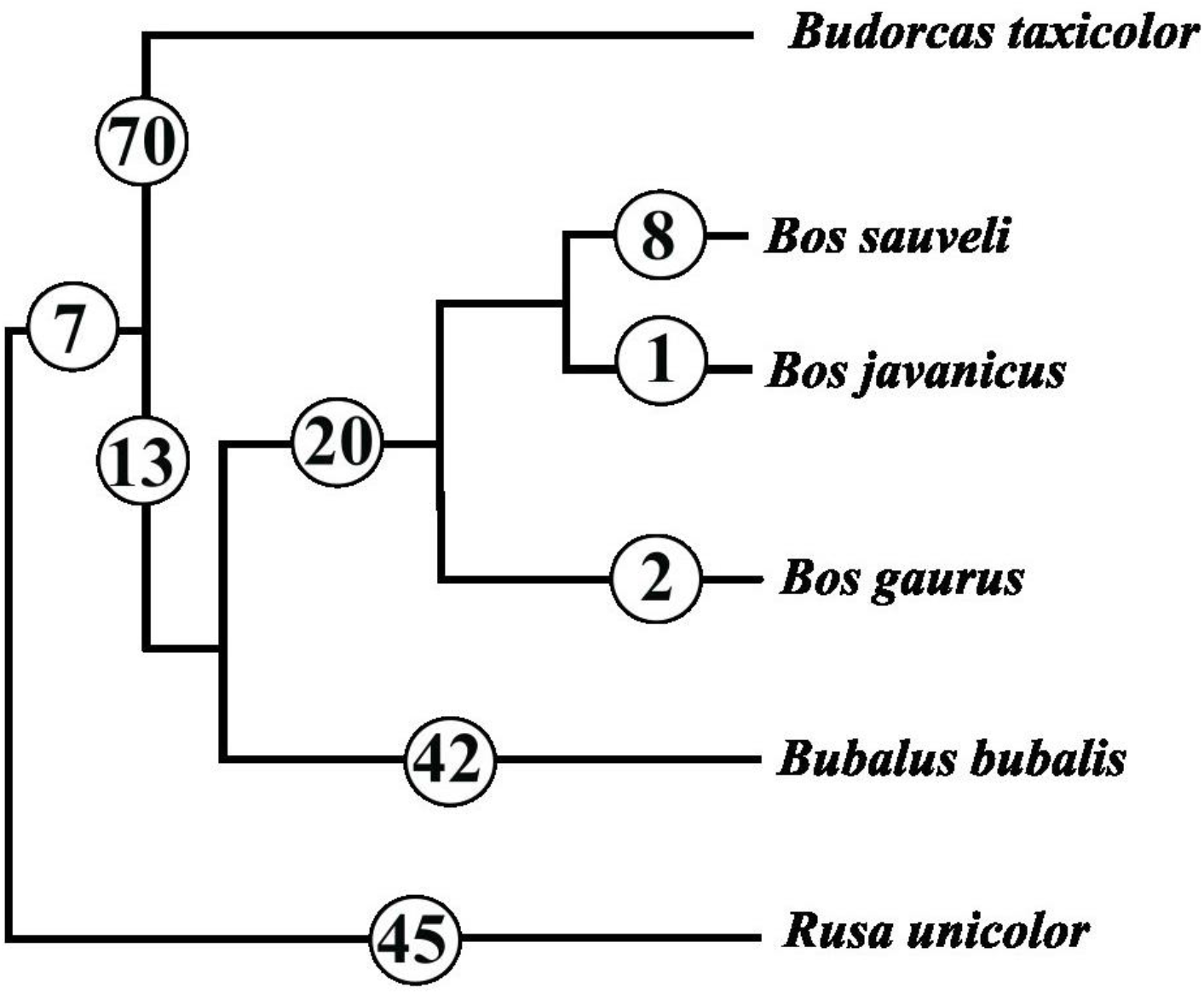
Distribution of the autapomorphic and synapomorphic character states listed in the text and in Supplementary Material 06 (Chantasri et al., 2026). The backbone tree is a composite of Fernández & Vrba (2005) for the higher phylogeny, Calamari (2021) for the Bovidae, and Sinding et al. (2021) for the position of *Bos sauveli*.

Taken as a whole, 7 characters allow for the full discrimination of the two families represented (5, 9, 32, 47, 100, 123, 148), 10 for the three subfamilies represented (**21**, **22**, **27**, **29**, **49**, **66**, **72**, **77**, **83, 122**), 12 for the four genera represented (**24**, **31**, **40**, **65**, **67**, **80**, **93**, **97, 130**, **131**, **144**, **145**), and none for the six species represented.

The maximum frequencies are extremely high overall and show comparatively little variation between characters and taxa. When all taxa are considered together, the mean maximum frequency is 94.8% (median = 100%), with a standard deviation of 12.0% and values ranging from 33.3% to 100%. This reflects the fact that 80.0% of the character states exhibit maximum frequencies above 95%, indicating high stability, while only 8.1% fall below 70% and are thus strongly variable. The lowest values are not randomly distributed: they are concentrated in the characters located on a few specific bones, which show mean maximum frequencies far lower than the others. Indeed, while most bones present mean maximum frequencies equal to 100% (5/27) or above 95% (12/27), only a few elements present means of 90–94% (7/27), and only three bones have a mean below 90% (the humerus, the malleolar bone, and the astragalus). Among these, the astragalus performs exceptionally poorly, with a mean of only 81.8%. It is worth noting that all the characters coded on the sesamoid bones were recorded on very small samples of specimens (between 22 % and 13.8 % of the complete dataset). Their results should therefore be treated with much caution. The axial large sesamoid is absent from the RF and NB accuracy estimates as its characters were not scored enough times to be included. The distribution of these scores among the taxonomic units investigated reflects the number of unique character states observed in each of them (Table 01). All monospecific taxonomic ranks perform very well (means above 90%), while the units composed of 5 (Bovidae) to 3 (Bos) taxa perform progressively better as the number of species they include decreases (from means of 66% to 89%) (Table 01).

**Table 01.** Mean across bones of the highest character-state frequency observed for each character. Values are reported for all the higher taxonomic ranks encountered in this study.

| Bone | Total | *Rusa*<br>Cervidae | *Budorcas*<br>Caprinae | *Bubalus* | Bovidae | Bovinae | Bos |
| --- | --- | --- | --- | --- | --- | --- | --- |
| First phalanx | 100.0% | 100.0% | 100.0% | 100.0% | 47.2% | 60.0% | 100.0% |
| Femur | 100.0% | 100.0% | 100.0% | 100.0% | 72.1% | 80.0% | 100.0% |
| Axial large sesamoid | 100.0% | 100.0% | 100.0% | 100.0% | 71.4% | 76.9% | 100.0% |
| Scaphoid bone | 100.0% | 100.0% | 100.0% | 100.0% | 68.1% | 77.2% | 99.1% |
| Small sesamoid | 100.0% | 100.0% | 100.0% | 100.0% | 87.0% | 87.9% | 100.0% |
| Magnum bone | 99.0% | 97.6% | 96.2% | 100.0% | 58.8% | 71.4% | 91.1% |
| Lunatum bone | 98.6% | 100.0% | 100.0% | 100.0% | 57.0% | 71.6% | 98.3% |
| Third phalanx | 98.4% | 97.1% | 100.0% | 98.6% | 69.0% | 76.3% | 90.0% |
| Second phalanx | 98.2% | 98.9% | 93.3% | 100.0% | 61.4% | 62.8% | 91.2% |
| Pyramidal bone | 97.4% | 92.7% | 96.4% | 100.0% | 58.7% | 64.4% | 88.9% |
| Metacarpus | 97.3% | 95.2% | 100.0% | 97.3% | 60.0% | 78.3% | 92.7% |
| Abaxial large sesamoid | 97.2% | 100.0% | 100.0% | 83.3% | 88.6% | 92.9% | 100.0% |
| Pisiform bone | 97.0% | 96.3% | 100.0% | 93.3% | 70.6% | 78.3% | 84.5% |
| Metatarsus | 96.8% | 100.0% | 100.0% | 96.6% | 65.5% | 77.2% | 89.0% |
| Tibia | 96.7% | 94.1% | 98.3% | 97.3% | 66.6% | 70.6% | 88.2% |
| Coxal bone | 96.4% | 98.9% | 92.5% | 93.5% | 72.6% | 76.2% | 97.0% |
| Large cuneiform | 95.5% | 80.4% | 92.9% | 100.0% | 56.7% | 62.5% | 100.0% |
| Calcaneus | 94.9% | 91.6% | 94.1% | 97.9% | 60.4% | 70.1% | 87.7% |
| Radius | 94.5% | 99.0% | 95.8% | 100.0% | 59.7% | 62.0% | 69.1% |
| Unciform bone | 94.3% | 97.2% | 96.9% | 87.2% | 71.9% | 85.3% | 91.2% |
| Navicular-cuboid | 93.3% | 71.5% | 92.9% | 98.9% | 64.4% | 75.3% | 83.5% |
| Ulna | 93.1% | 92.7% | 95.8% | 92.7% | 81.1% | 88.2% | 92.7% |
| Patella | 91.9% | 79.2% | 100.0% | 97.2% | 84.0% | 87.5% | 88.3% |
| Scapula | 91.8% | 89.4% | 93.3% | 93.7% | 65.7% | 67.3% | 82.2% |
| Humerus | 88.1% | 92.7% | 95.9% | 94.4% | 60.2% | 63.1% | 68.6% |
| Malleolar bone | 85.9% | 78.3% | 100.0% | 86.0% | 48.8% | 49.8% | 67.2% |
| Astragalus | 81.8% | 73.7% | 75.3% | 83.2% | 49.3% | 54.1% | 63.5% |
| **Mean** | **95.5%** | **93.2%** | **96.7%** | **96.0%** | **65.8%** | **72.9%** | **89.0%** |

Although the overall performance of the raw characters was strong enough to make them useful, we chose to perform additional analyses in order to further strengthen them and to align their use with the same protocol as the one described in the study of the smaller ruminants.

### III) Results of the Random Forest and Naïve Bayes approaches

The analyses we performed combined several characters from single bones but did not combine different bones to simulate an application to the archaeological record, where the bones are most often found isolated. Because they have been registered on a small number of specimens and taxa, sesamoid bones are excluded from all the following inter-bones comparisons.

In general, both the Random Forest (RF) and Naïve Bayes (NB) approaches performed very similarly in our tests. The ability of the analyses to correctly identify a bone is strongly influenced by the taxonomic rank considered. Indeed, correct family-level identification can be achieved on average in 93.8% (NB) or 94.5% (RF) of cases. For subfamily-level identification, this percentage slightly decreases to 92.8%/93.1% (NB and RF), then to 89.2%/89.0% (NB and RF) for genus-level identification, and to 76.0% (NB) or 75.7% (RF) for species-level identification.

The number of characters considered (**Fig. 92-A**) also has a strong impact on the reliability of the results. Indeed, shifting from one to six input characters improves identification reliability for every taxonomic rank (+35.0% for species level, +26.9% for genus level, +18.4% for subfamily level, and +14.2% for family level). There are also differences between bones, as not all anatomical parts have the same discriminatory power. The mean identification accuracy is 87.3% if the sesamoids are excluded, with only three bones showing accuracies lower than 80% (3^rd^ Phalanx, Ulna, and Navicular-cuboid), and then growing numbers of bones performing better, including 11 elements with accuracies above 90% (**Fig. 92-B**). Sesamoids are excluded here because the results they provide are extremely fragile, given the small number of specimens on which these characters could be recorded; the axial sesamoids could not be included in these analyses at all, having been scored on too few specimens (N = 15).

**Figure 92.**
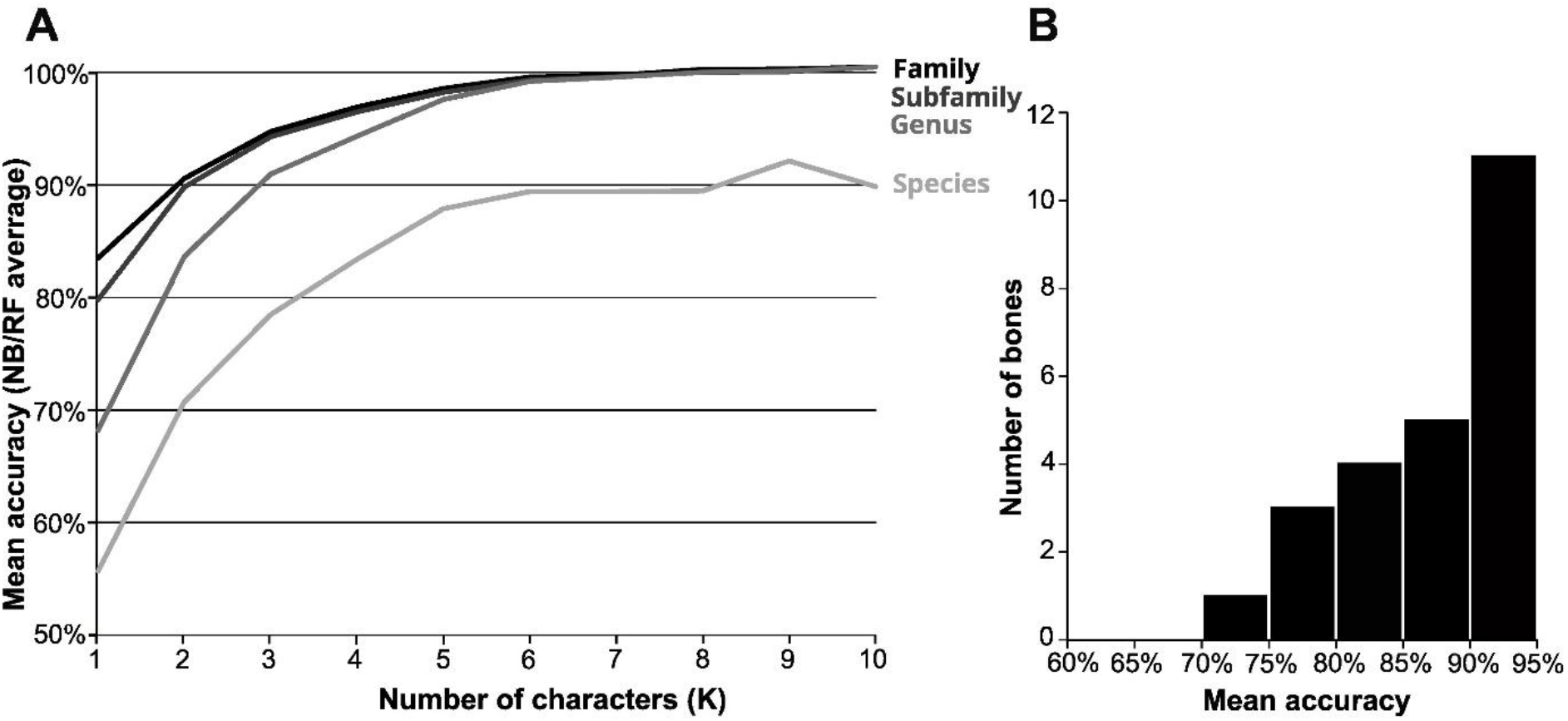
A) Mean accuracy of the identifications for a given number of characters (K) and a given taxonomic rank. Characters are not mixed between several bones. B) Mean accuracy of the considered anatomical parts.

The complete list of mean accuracies for the different ranks, bones, and numbers of characters k is reported in Supplementary Material (Supplementary Material 07; Chantasri et al., 2026). It provides a reliability estimate for each possible case.

There are significant disparities between the identification accuracies of the different taxa, mainly because of the taxonomic coverage of the study that is strongly imbalanced, and because some of our species are documented by a small number of specimens.

At the family rank, Cervidae are identified with 88.5% (RF) or 84.7% (NB) accuracy, while Bovidae present accuracies above 99% for both methods. At the subfamily rank, Bovinae (98.9% RF, 98.9% NB) are better identified than Caprinae (90.7% RF, 90.5% NB), and Cervinae (89.0% RF, 87.0% NB). All the genera present very good accuracies with *Bos* being the easiest to identify (97.0% RF, 97.8% NB), followed by *Budorcas* (91.6% RF, 91.0% NB), *Rusa* (89.9% RF, 88.9% NB), and *Bubalus* (89.6% RF, 88.7% NB). Regarding species, those being the sole representatives of their family, subfamily, or genus are identified fairly well: *Rusa unicolor* (95.3% RF, 95.1% NB), *Bubalus bubalis* (93.9% RF, 94.5% NB), and *Budorcas taxicolor* (91.1% RF, 91.6% NB). However, the identification becomes way less accurate for the species of *Bos,* with still correct accuracy for *Bos gaurus* (77.0% RF, 78.3% NB) but lower accuracy for *Bos javanicus* (58.5% RF, 57.2% NB) and for *Bos sauveli* (25.7% RF, 34.2% NB). The random thresholds being 50% for families, 33% for subfamilies, 25% for genera, and 16.6% for species, it is only really approached by *Bos sauveli*, and to a lesser extent by *Bos javanicus,* which are morphologically close to each other and, in the case of *Bos sauveli,* poorly represented in the dataset with only three specimens.

## Discussion

### General remarks regarding the reliability of the described characters for taxonomic identification

The analyses of characters initially defined on a small set of specimens and subsequently tested on a larger comparative sample provide several insights into character performance and taxonomic discrimination. In contrast to the results reported for the medium-sized ruminants (Mithong et al., 2026), the testing phase indicates that the discriminatory power of most characters described here remained very strong when assessed across the full comparative sample. Indeed, 80% of the recorded character states showed a maximum state frequency above 95% within their respective taxon. This result is broadly consistent with previous studies that used osteological characters to distinguish morphologically different taxa in samples considering a limited number of species (Bochaton et al., 2016; Hanot & Bochaton, 2018; Zhang et al., 2022). This overall pattern, however, does not hold uniformly across the dataset. Character stability and classification accuracy both decrease markedly for the three species of the genus *Bos*. These three species are the only ones that are not the sole representative of their genus in the dataset and are, in addition, all closely related (Hassanin & Ropiquet, 2007; Sinding et al., 2021). These species are thus much closer morphologically to one another than all the other species of the dataset are. The overall excellent results of this study with respect to the studies performed on the medium and small taxa are mainly explained by the strong morphological disparity and the low number of species included in the present dataset. This mirrors the challenges faced by any anatomical identification regardless of the method used and does not question the quality of the data acquisition. However, this makes the two studies difficult to compare, and some taxa will remain much easier to identify reliably than others just because the pool of possibilities strongly differs between the different size and taxonomic groups.

Despite the excellent performance of our character set, to keep our analytical protocol consistent with that applied to the small and medium-sized ruminants (Mithong et al., 2026), we also implemented Random Forest and Naïve Bayes classifiers to combine several characters and quantify identification reliability at every taxonomic rank. This approach provided further support for the strong performance of our character set at most taxonomic ranks, while also explicitly quantifying the more limited reliability observed among the species of *Bos*, an outcome that a character-by-character assessment alone might have obscured. When a combination of characters is considered, it becomes possible to identify all the taxa with more or less uncertainty. Only the identification of *Bos sauveli* remains impossible in most instances.

### How to use our characters to study zooarchaeological and paleontological material

The characters described and tested in this study have been defined on a limited number of taxa and thus could not be used to identify bone pieces not belonging to these taxa. The user should thus make sure the bone they are trying to identify belongs to a member of Ruminantia, a basic step that can be done with a straightforward comparison with the pictures included in this paper, 3D models of any taxa available online (e.g., on Morphosource.org), or a comparative skeleton if available. As indicated in the Material and Methods section of this paper, the small and medium-sized Southeast Asian Ruminantia species have been excluded from the present sample and analyzed separately in the previous paper of this series (Mithong et al., 2026). The user should first make certain that they are not trying to identify a member of this latter group (genera *Axis, Capricornis, Cervus, Elaphodus, Moschus, Muntiacus, Naemorhedus, Pseudois*, and *Rucervus).* In case of doubt, this could be assessed using the morphometric data from the first paper of this series (Seesod et al., 2026).

The good discriminative performances of the characters described in this study could allow the user to use them independently, but using the web application associated with this paper will automatically provide the estimation of reliability associated with each identification in addition to allowing the combination of several characters through our Naïve Bayes model. The standalone HTML application (Bochaton et al., 2026) can be run directly from any web browser and includes all the character descriptions. Once inside the application, the user selects the bone and inputs any number of observable character states from a single anatomical element. Combinations of characters from different anatomical elements are not allowed, as the models were generated and validated separately for each element. After input, the application provides attribution probabilities at the family, subfamily, genus, and species levels. Each taxonomic rank is calculated independently and involves a different number of possible taxa.

Because character states are not always unique to a single species, results could be ambiguous, reflecting morphological proximity among taxa and sheer morphological variability. Sampling imbalance may also influence the relative attribution probabilities, as illustrated by *Bos sauveli*, which is represented by only three specimens. It is difficult to define a strict threshold for ambiguity; however, given that our dataset comprises relatively few taxa at each rank (two families, three subfamilies, four genera, and six species), the corresponding expected probability under a random attribution is already comparatively high (50%, 33.3%, 25%, and 16.7%, respectively). How much a probability above that baseline is worth is addressed below.

The tests reported in the Material and Methods section show that an attribution is correct in more than 90% of cases from 80% upwards at the family, subfamily and genus ranks, and from 90% upwards at the species rank (Supplementary Material 04, Chantasri et al., 2026). In order to retain the most precise attribution that can be relied upon, the following rules should be applied: 1) retain the lowest rank whose attribution reaches the threshold set for that rank, and 2) record the element as Ruminantia indet. where none does. In our tests, the species rank is reached for 64.9% of the character combinations tested, the genus rank for 86.1%, and an attribution retained under these rules proved correct in 99.7% of cases at every rank. As indicated by our tests, neither the margin separating the two most likely taxa at a given rank nor the mutual consistency of the attributions returned at the four ranks adds any reliability once these thresholds are applied. These rules are applied automatically by the application, which highlights the lowest rank that can be retained, or highlights none where no attribution meets its threshold, and which displays beside every rank the reliability observed for the probability returned.

If necessary, the character-based results can be complemented with the metrical criteria developed in the first paper of this series (Seesod et al., 2026) to help resolve ambiguous attributions using size data. This may be particularly useful for morphologically similar taxa that differ in size, such as the species of the genus *Bos*. In such cases, it should be clearly stated that the specific identification is based on size criteria and is therefore subject to the potential biases previously discussed (Seesod et al., 2026). Whatever approach is adopted, it should be clearly described in the Material and Methods section of the study.

## Conclusion

The results presented here show that the postcranial skeleton of large-sized Southeast Asian ruminants carries reliable taxonomic information, supporting reliable identification down to the genus level and, for half of the taxa, to the species level. The main limitation concerns the closely related species of the genus *Bos*, particularly the poorly represented *Bos sauveli*, for which improved species-level discrimination will likely require additional comparative material and/or complementary approaches, including biomolecular methods. Combined with the osteometric protocol of Seesod et al. (2026), this work provides a reproducible framework for identifying the postcranial remains of large-bodied ruminants in continental Southeast Asia, bones for which comprehensive postcranial identification criteria have been lacking, despite their frequent occurrence in the region’s zooarchaeological record. Our approach clarifies the limits of what can reasonably be achieved through morphological identification alone, while providing a complementary toolkit to perform taxonomic identification in a repeatable and independently assessable manner. This approach can also help identify which specimens are best suited for further analysis, particularly biomolecular approaches, whether to gather additional information (e.g., isotopic analyses) or to refine taxonomic identification further, such as ancient DNA. Given the important role played by large bovines and cervids in the subsistence strategies of past hunter-gatherer and early farming populations across the region, this protocol can facilitate ongoing and future zooarchaeological and paleontological studies, while also providing a template that could be extended to other biodiverse tropical settings facing similar identification challenges. The approach could also be adapted to other regions and taxa, as an essential step towards reproducibility and transparency in zooarchaeological taxonomic identification, and one that will make such data more relevant to broader studies of past environments and human-environment interactions, which depend on reliable baseline taxonomic data, particularly in biodiversity-rich areas that remain poorly documented from this perspective.

## Acknowledgements

The authors are grateful to the museums, universities, and staff members who supported this study. We especially thank the Muséum national d’Histoire naturelle (Paris, France) and Joséphine Lesur; the National Museum of Natural History (Washington, D.C., USA) and Darrin Lunde; the Zoologische Staatssammlung München (Munich, Germany) and Anneke H. van Heteren; the Museum für Naturkunde (Berlin, Germany) and Christiane Funk; and the Royal Belgian Institute of Natural Sciences (Brussels, Belgium) and Olivier Pauwels.

## Data, scripts, code, and supplementary information availability

Supplementary information is available online: https://doi.org/10.23708/76NWAY; Chantasri et al., 2026.

## Author contributions

**Conceptualization**: Sirikanya Chantasri and Corentin Bochaton; **Data curation**: Sirikanya Chantasri and Corentin Bochaton; **Formal analysis**: Sirikanya Chantasri, Corentin Bochaton, and Julien Claude; **Funding acquisition**: Corentin Bochaton; **Data acquisition**: Sirikanya Chantasri; **Supervision**: Corentin Bochaton; **Writing original draft**: Sirikanya Chantasri, and Corentin Bochaton; **Writing – review & editing**: All the authors.

